# Chromatin binding of survivin regulates glucose metabolism in the IFN-γ producing CD4^+^ T cells

**DOI:** 10.1101/2021.10.05.463166

**Authors:** Malin C. Erlandsson, Karin M.E. Andersson, Nina Y. Oparina, Venkataragavan Chandrasekaran, Anastasios Damdimopoulos, Maria-Jose Garcia-Bonete, Zakaria Einbeigi, Sofia T. Silfverswärd, Marcela Pekna, Gergely Katona, Maria I. Bokarewa

## Abstract

Interferon-gamma (IFNγ) producing T cells develop metabolic adaptation required for their effector functions in tumour biology, autoimmunity and antiviral defence.

Using sorted CD4^+^ cells we demonstrated that glycolytic switch and high glucose uptake in IFNγ-producing cells was associated with survivin expression. Inhibition of survivin restored glycolysis by upregulating the transcription of phosphofructokinase PFKFB3 and reducing glucose uptake. Integration of the whole-genome sequencing of the chromatin immunoprecipitated with survivin with transcription changes in CD4^+^ cells after survivin inhibition revealed co-localization of survivin, IRF1 and SMAD3 in the regulatory elements paired to the differentially expressed genes. Western blot demonstrated direct binding of survivin to IRF1 and SMAD3. Functionally, inhibition of survivin repressed IFNγ signalling and activated SMAD3-dependent protein remodelling, which resulted in the effector-to-memory transition of CD4^+^ cells. These findings demonstrate the key role of survivin in IFNγ-dependent metabolic adaptation and identify survivin inhibition as an attractive strategy to counteract these effects.

## Introduction

Interferon-gamma (IFNγ) producing T-helper cells are fundamental for immune surveillance and exhibit unique protective properties in cell-mediated and humoral immune responses that control intracellular invasion of pathogens and unwanted tumour and tissue growth. IFNγ producing cells are effector cells that migrate, proliferate and produce signal molecules at the site of inflammation to mobilize immunity and to resolve inflammation (*1, 2*). Inability to achieve resolution converts the IFNγ-producing cells into facilitators of adaptive immune responses contributing to chronic autoimmune inflammation. In this role, IFNγ counteracts Th17 and Th2 dependent inflammation, controls TGFβ dependent fibrotic processes and exerts tissue-protective effects restricting expansive tissue growth. These IFNγ dependent mechanisms are shared between the canonical autoimmune diseases type I diabetes mellitus (T1D), multiple sclerosis (MS), systemic lupus erythematosus (SLE) and rheumatoid arthritis (RA) and are central for their pathogenesis (*3*). The program of the Genome-Wide Association Studies (GWAS) revealed strong connection between autoimmune disease development and treatment response and the genetic polymorphism within and close to principal mediators of IFNγ signalling, the IFN response factors (IRFs) and IFN-stimulated genes (ISGs)(*4*). Concordant changes in the expression signature of ISG across autoimmune pathologies shown by functional comparative analysis of the blood leukocytes and target tissues warrants broad application of IFN-targeting strategy (*5-9*). However, the dual role of IFN protection against pathogens and auto aggression, limits the benefits of available IFN targeting therapies, and a more selective approach is needed.

Expression of ISGs is initiated by activation of IFNγ receptor, production of IRFs and binding of the IRF-specific regulatory elements on the chromatin. Analysis of the IRF-bound chromatin revealed their functional specificity evidenced by the ability to cooperate with multiple transcription factors (TF) (*10*). The collaborative effects of IRFs and TFs of the E-twenty-six specific (ETS) and the basic leucine zipper (bZIP) protein families regulates the impact of IFNγ in pro-inflammatory effector functions and in anti-TGFβ fibrotic processes that fuels autoimmunity. The exact molecular mechanisms to balance between these effects of IFNγ have not been fully mapped.

To be efficient in their effector functions and persistently respond to stimuli from the outside, IFNγ-producing cells require energy. Thus, IFNγ-producing cells undergo metabolic adaptation by switching glucose metabolism from the citric acid cycle to pentose phosphate pathway alternative of glycolysis thereby increasing the availability of nucleotides, amino acids, and fatty acids (*11, 12*). This dramatically increases glucose consumption to sustain the increased energy demands of the IFNγ-producing cells, which manifest in their effector phenotype and high migration capacity (*13-15*). The inhibition of anabolic adaptation by pharmacological or dietary interventions shows promising results in animal models of autoimmune diseases, however the therapeutic window and long-term safety of pharmacological modulation of glycolysis modification in the clinic remains to be determined.

Metabolic adaptation of T cells varies between the autoimmune conditions. In RA, a redistribution of energy production is achieved by controlling expression of 6-phosphofructo-2-kinase/fructose-2,6-bisphosphatase 3 (PFKFB3), which generates and breaks down fructose-2,6-bisphosphate, an allosteric regulator of phosphofructokinase 1 (*16*). Downregulation of PFKFB3 activates glucose-6-phosphate dehydrogenase (G6PD) and 6-phosphogluconolactonase (PGLS) and directs the glucose processing into the pentose phosphate pathway. In T1D, MS and in SLE, the cells experience no energetic starvation and rely on the pyruvate kinase dependent hyperproduction of lactate (*17-20*). Despite these differences, inhibition of PFKFB3 in experimental MS, graft-versus-host disease, and T1D rebalances the metabolic alterations and arrests the effector ability of T cells (*17, 21*).

Survivin, coded by *BIRC5* gene, is the proliferation-supporting protein widely expressed in solid and haematological malignancies (*22, 23*). In autoimmune conditions, survivin has been attracting attention in a variety of diseases including autoimmune arthritis (*24*), psoriasis (*25*), and MS (*26*). Targeting survivin in experimental and clinical autoimmunity was efficient in reducing inflammation, proliferation, and tissue damage (*27-31*), while its role in cell metabolism has never been assessed.

Survivin is highly expressed in the bone marrow cells and is required for healthy haematopoiesis and for maintenance of the CD34^+^ stem cell population (*32*). Survivin is an essential for T cell development. Conditional deletion of survivin in thymocytes affects the transition from the double negative to the double positive thymocyte developmental stage resulting in the reduction of mature CD4^+^ and CD8^+^ T cell populations (*33*). Survivin deletion also disrupts the formation of a functional T cell receptor and causes inability to mount an adequate immune response upon antigen challenge (*34*). In mature T cells, survivin expression declines and re-appears during critical adaptive immune responses requirering phenotype transition such as the acquisition of the effector phenotype of CD4^+^ T cells and maintenance of virus-specific CD8^+^ memory T cells (*35, 36*). Differentiation into the regulatory CD25^+^FOXP3^+^ and the follicular CXCR5^+^BCL6^+^ T cells has been shown directed by survivin (*36-38*).

In dividing cells, survivin acts within the chromosomal passenger complex sensing mitotic spindle tension and protecting chromosomal cohesion (*39, 40*). During interphase, survivin localizes to the nucleus and cytoplasm with distinct biological functions (*41*). In cytoplasm, survivin interacts with the mitochondrial protein Diablo, which prevents its release from mitochondria and the activation of apoptosis. In the nucleus, survivin interacts with histone subunits, retinoblastoma transcriptional activator 1, cyclin dependent kinases 4 and 2 (*42, 43*). Although there are some functional indications for the role of survivin in the regulation of gene expression (*37, 44-46*), the overall function of survivin on the chromatin is not clear.

In this study, we present experimental evidence for the involvement of survivin in the regulation of the effector phenotype in IFNγ-producing Th1 cells. By ChIP-seq analysis of the survivin bound chromatin and transcriptional analysis of the IFNγ-activated CD4^+^ cells, we demonstrate that survivin is enriched on the regulatory elements (RE) of the genes sensitive to survivin inhibition. Survivin is recruited to the RE containing IRF-binding motifs through its direct interaction with IRF1 and SMAD3. On the chromatin, survivin controls the transcriptional programs governed by these two proteins by inhibition of PFKFB3. Since survivin also promotes glucose uptake, the pentose phosphate pathway is the only channel available for glucose. Inhibition of survivin restored the PFKFB3 dependent metabolism of CD4^+^ cells, improved control over the IFNγ-dependent effector activity and activated the SMAD3 dependent protein remodelling in CD4^+^ cells.

## Results

### Survivin is an essential determinant of IFNγ producing CD4 effector cells in RA patients

Flow-cytometry analysis of ConA stimulated CD4^+^ cells from the peripheral blood of 22 RA patients demonstrated that the effector (T_EFF_) cells defined as CD62L^neg^CD45RA^+/-^CD27^neg^ had high intracellular levels of survivin. These survivin-producing cells comprised up to 9.2% [5.4-16.4] of T_EFF_ population, which was the largest of CD4^+^ cell subsets expressing survivin (Figure 1A). The intensity of survivin signal in those T_EFF_ was also the highest.

**Figure 1.**
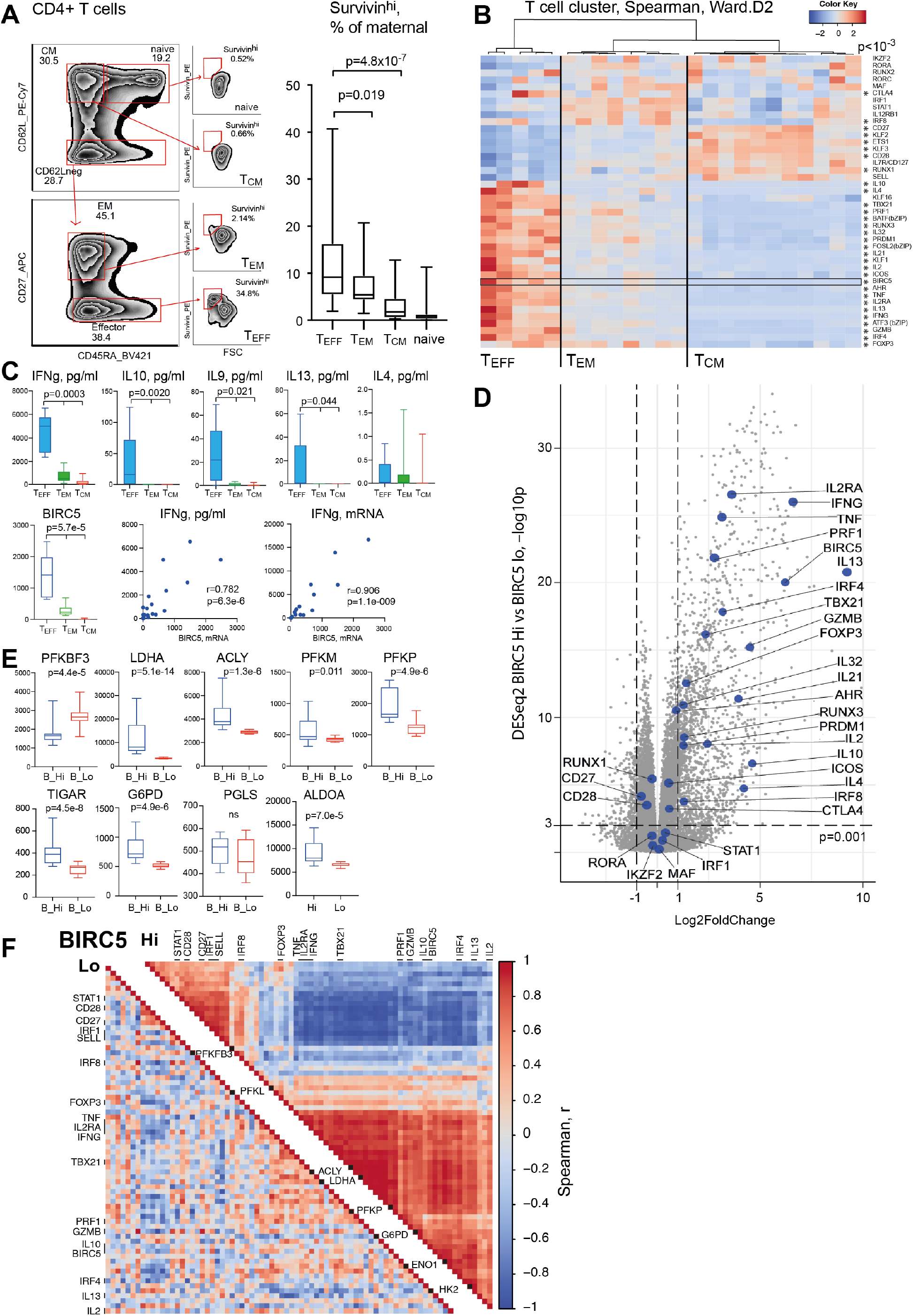
Survivin is essential for the phenotype of IFNγ producing CD4^+^ T cells. **A**. Gating for survivin^hi^CD4^+^ cells within naïve (CD62L^hi^CD45RA^+^), central memory (CM, CD62L^hi^CD45RA^neg^), effector memory (EM, CD62L^neg^CD45RA^neg^CD27^hi^), and effector (EFF, CD62L^neg^CD45RA^+/-^CD27^neg^) populations. Box plot of the frequency of survivin^hi^CD4^+^ cells (RA patients, n=22). P-values are obtained by the Wilcoxon test. Differential expression analysis by RNAseq of CD4^+^ cells stimulated with aCD3 for 48h (RA patients, n=24). P-values are obtained by DESeq2. **B**. Heatmap of Th core signature genes clustered by non-parametric Spearman distance and Ward.D2 linkage. (*) p<0.001 between T_EFF_ and T_CM._ **C**. Box plots of cytokines by protein levels in supernatants (p-values by Kruskal-Wallis test) and RNAseq (p-values by DESeq2) of CD4^+^ cells as above. **D**. Volcano plot of protein-coding genes in *BIRC5*^hi^ and *BIRC5*^lo^ cells (median split). Th1 signature genes are indicated. **E**. Box plots of normalized RNAseq values of the key glycolytic enzymes in *BIRC5*^hi^ and *BIRC5*^*lo*^ cells. **F**. Correlation matrix of Th1-signature genes and glycolytic enzymes in *BIRC5*^hi^ and *BIRC5*^lo^ cells. Unsupervised clustering was done by corrplot Bioconductor, R-studio.

To investigate the phenotype of the survivin-producing T_EFF_ cells, primary CD4^+^ cells from 24 RA patients activated with anti-CD3 antibodies were subjected to RNA-seq analysis. We performed unsupervised clustering of CD4^+^ samples by the expression of 41 core signature genes characteristic for T-helper subsets (*47*), and identified the samples that were predominantly composed of T_EFF_ cells (Figure 1B). These cells produced high levels of Th1 signature genes including *TBX21, EOMES, IL2RA* and cytokines IFNγ, IL9, IL10, IL13 (Figure 1C). The levels of *BIRC5* mRNA, coding for survivin, gradually increased from T_CM_ CD4^+^ cluster to T_EM_ (p=1.2×10^−4^) and to T_EFF_ (p=1.15×10^−18^) clusters with strong correlation to *INFg* mRNA (r=0.906, p=10^−9^) and protein (r=0.782, p=10^−6^) levels (Figure 1C). The comparative analysis of RNAseq from CD4^+^ cells with high (*BIRC5*^hi^) and low (*BIRC5*^lo^) expression of survivin showed difference for classical features of IFNγ-producing Th1 cells including the lineage specific cytokines, core transcription factors and cytotoxic molecules perforin and granzyme B (Figure 1D).

Glucose availability and metabolism plays a key role in regulation of IFNγ production and cytotoxicity of Th1 cells (*48*). We found that *BIRC5*^hi^ and *BIRC5*^lo^CD4^+^ cells differed in the expression of glucose metabolism enzymes (Figure 1E). Specifically, *BIRC5*^hi^CD4^+^ cells were deficient in the key regulator of glucose processing *PFKFB3* leading to shunting of glucose to the pentose phosphate pathway as evidenced by increased expression of glucose-6-phosphate dehydrogenase (*G6PD*) and 6-phosphogluconolactonate (*PGLS*), increased expression of ATP citrate lyase (*ACLY*), indicating increased fatty acid metabolism, and up-regulation of *TIGAR* (TP53 induced glycolysis regulatory phosphatase). In agreement with its central role in glucose metabolism, *PFKFB3* expression and its expression relative to *G6PD* and *LDHA* correlated inversely to *BIRC5, IFNγ*, transcription factors *TBX21, EOMES, AHR* and cytotoxic molecules perforin and granzyme B (Supplementary Figure S1B). The correlation matrix of the signature Th genes and glycolysis markers revealed clear distinctions in biological processes within *BIRC5*^hi^ and *BIRC5*^lo^CD4^+^ cells (Figure 1F). The associations between high expression of *BIRC5*, IFNγ-dependent CD4^+^ cell phenotype and utilization of glucose point to the role of survivin in regulation of glycolysis in IFNγ-producing CD4^+^ cells.

### Chromatin bound survivin is predicted to regulate the carbohydrate metabolism

Previous reports on survivin binding to specific regions of the genomic DNA indicated its potential role in regulation of gene transcription (*37, 44, 45*). This prompted us to investigate the chromatin binding of survivin on the whole genome scale.

We performed survivin ChIP-seq analysis of 12 individual CD4^+^ cell cultures pooled in 4 independent replicates. Peak calling of the sequenced DNA material revealed 13704 nonredundant peaks with enrichment significance (adjusted p<10^−5^). These survivin-ChIP peaks were widely distributed across the human genome with specific enrichment in the regions with promoter function (Figure 2A). About 40% and 70% of the survivin-ChIP peaks were located within 10kb and 100kb, respectively, from known genes. The peaks were frequently colocalized with the chromatin areas occupied by transcriptional regulators, promoters and enhancers located in 10-100kb vicinity, while CTCF binding sites and insulator regions were located at a shorter distance of 1-10kb (Figure 2B). Interval co-localization analysis for survivin-ChIP peaks with 0, 1, 10, 50kb flanks demonstrated statistically significant enrichment of the prevalent DNA elements (Figure 2C).

**Figure 2.**
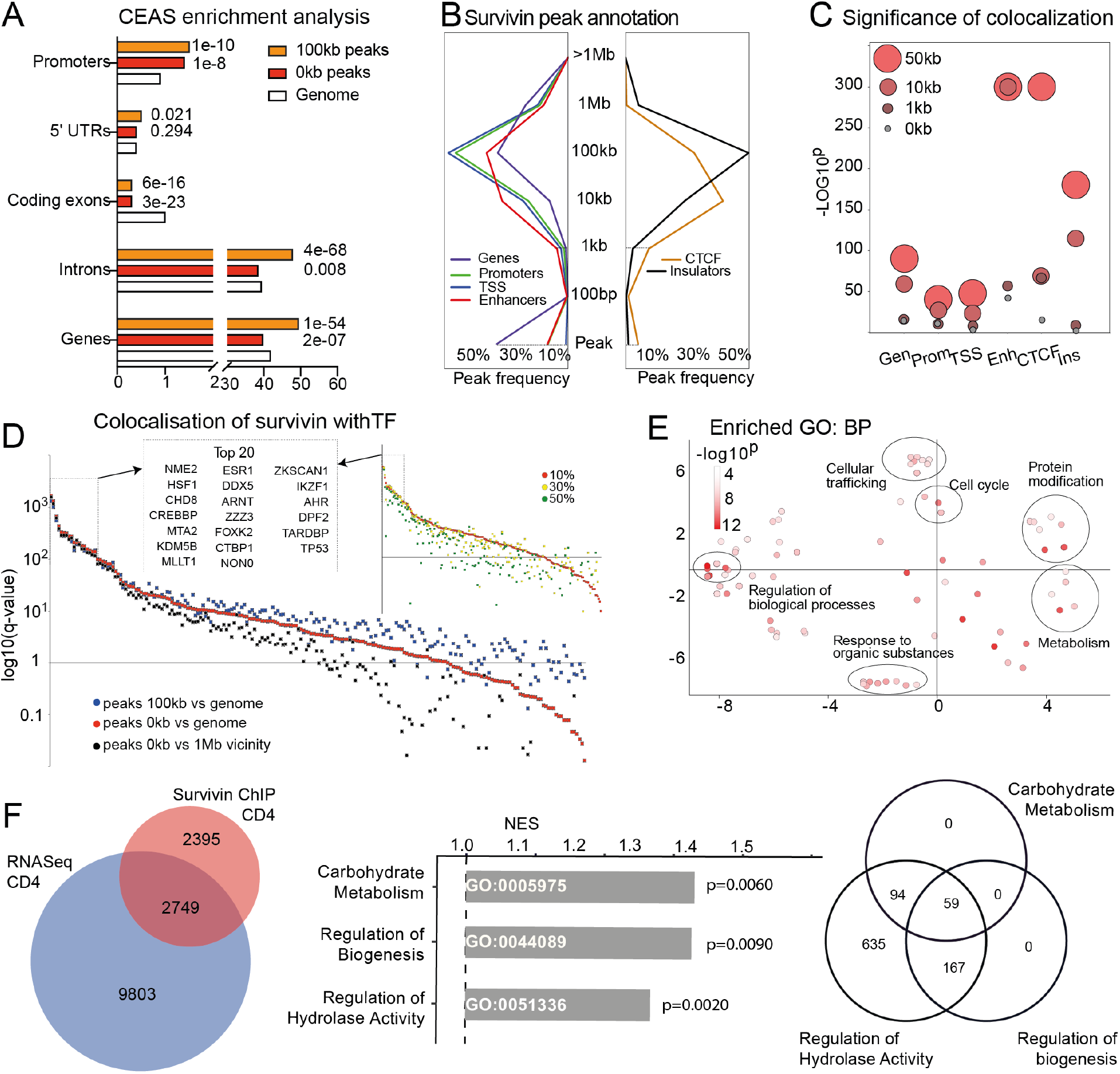
Survivin binding to chromatin is predicted to regulate carbohydrate metabolism. Survivin-ChIPseq was performed from CD4^+^ T cells (n=12). Survivin peaks (enrichment against input, p<10^−5^) were mapped to hg38 human genome. **A**. Bar plots of survivin peaks distribution (red bars 0kb, orange bars 100kb flanks) compared to the genome (open bars). **B**. Position of survivin peaks and colocalization with DNA elements. **C**. Dot plot of two-tailed p-values obtained by Fisher’s exact test. **D**. Dot plot of individual q-values for colocalization of survivin and TFs peaks by ChIPseq (ReMap2020, n=1034) (red dots, 0kb flanks, 10% overlap; blue dots, 100kb flanks; black dots, 1Mb genome neighborhood). Only TFs with >100 events are shown. **Insert**. Dot plot of q-values for colocalization by overlap >10% (red dots), >30% (yellow dots), and >50% (green dots). **E**. Semantic similarity map of biological processes (GO:BP) regulated by TFs colocalized with survivin. Functional annotation was done in MetaScape. **F**. Venn diagram of the protein-coding genes annotated to survivin peaks and those identified by RNAseq of CD4^+^ cells. Bar plot presents normalised enrichment score (NES) for GO:BP and p-values identified in GSEA. Venn diagram of the three enriched GO:BP.

Next, we characterized the TF landscape of the survivin bound chromatin regions by annotating survivin-ChIP peaks to the global ChIPseq dataset for 1034 human transcriptional regulators integrated in the ReMap database (*49*). We identified 146 TF candidates significantly enriched across the settings of survivin-ChIP peaks with 0kb and 100kb flanks (Figure 2D). For comparison, we analysed areas within 1Mb of the peaks. In each dataset, the minimal threshold for an overlapping peak was set to 10%, equally for survivin and other TF candidates. We observed a tight agreement between the TF candidates identified by these two approaches (Figure 2D). Notably, the q-significance of association between TFs and survivin increased in the regions of 0 to 100kb and lower for the local 1Mb vicinity comparison. Subsequently, increasing the minimal peak overlap from 10% to 30% and 50% revealed highly specific TF candidates co-localized with survivin-ChIP peaks (Figure 2D, insert). Among the top TFs, we noted those regulating glucose and insulin metabolism including CREBBP, KDM5B, DDX5, ARNT, FOXK2, CTBP1, IKZF1, and AHR.

Since the survivin-colocalized TF were found with no pre-selection by tissue specificity, we analysed their functional enrichment via MetaScape. TFs co-localized with survivin were compared to the background of all TF ChIPseq in the ReMap database. This approach identified the biological processes potentially regulated by survivin (Figure 2E) including functional groups for regulation of cellular trafficking, cell cycle, protein modification and metabolism. Other functional groups regulate response to organic substances including glucose and carbohydrates, pro-inflammatory cytokines and chemokines (Figure 2E. Supplementary Table 2S). To determine the physiological role of survivin-ChIP peaks in CD4^+^T cells, we compared the protein-coding genes annotated to the survivin-ChIP peaks with those transcribed in CD4^+^ cells (RNAseq, normalized raw counts above 0.5). We identified 2749 protein-coding genes potentially controlled by survivin (Figure 2F) and their subsequent analysis in GSEA revealed significant enrichment for the carbohydrate metabolic processes (GO:0005975), regulation of cellular biogenesis (GO:0044089) and hydrolase activity (GO:0051336) (Figure 2F), which show substantial overlap.

Analysis of survivin-ChIP peaks distribution suggests that survivin is enriched within or in the vicinity of the chromatin areas occupied by *cis*-regulatory elements. Functional annotation of survivin-ChIP peaks to the nearest genes and co-localized TFs indicates the role of survivin in the regulation of carbohydrate metabolism.

### Survivin restricts PFKFB3 expression and changes the metabolic requirements of CD4^+^ cells

To gain experimental evidence for the role of survivin in the associated biological processes, we inhibited survivin in freshly isolated CD4^+^T cells using YM155, a specific synthetic inhibitor of survivin function (*44*). Cell were activated with IFNγ for the final 2h. RNAseq analysis of the YM155-treated (0 and 10 nM) CD4^+^ cells identified the complete set of the DEGs (DESeq2, p<0.05). Comparison of DEGs to the list of protein-coding genes common for the survivin-ChIPseq and RNAseq demonstrated that 11.8% and 4.5% of the genes were sensitive to survivin inhibition for 24h and 72h, respectively (Figure 3A). Using the curated TRRUST database of gene regulatory relationship, we identified the central metabolism regulators MYC, SP1 and HIF1a as the upstream transcriptional regulators governing these survivin sensitive DEGs both after 24h and 72h of survivin inhibition. Other effects were attributed to the activity of JUN, NF-kB, RELA, SMAD4, ETS1 (24h) and later, to IRF1 and the class II trans-activator (72h) (Figure 3B).

**Figure 3.**
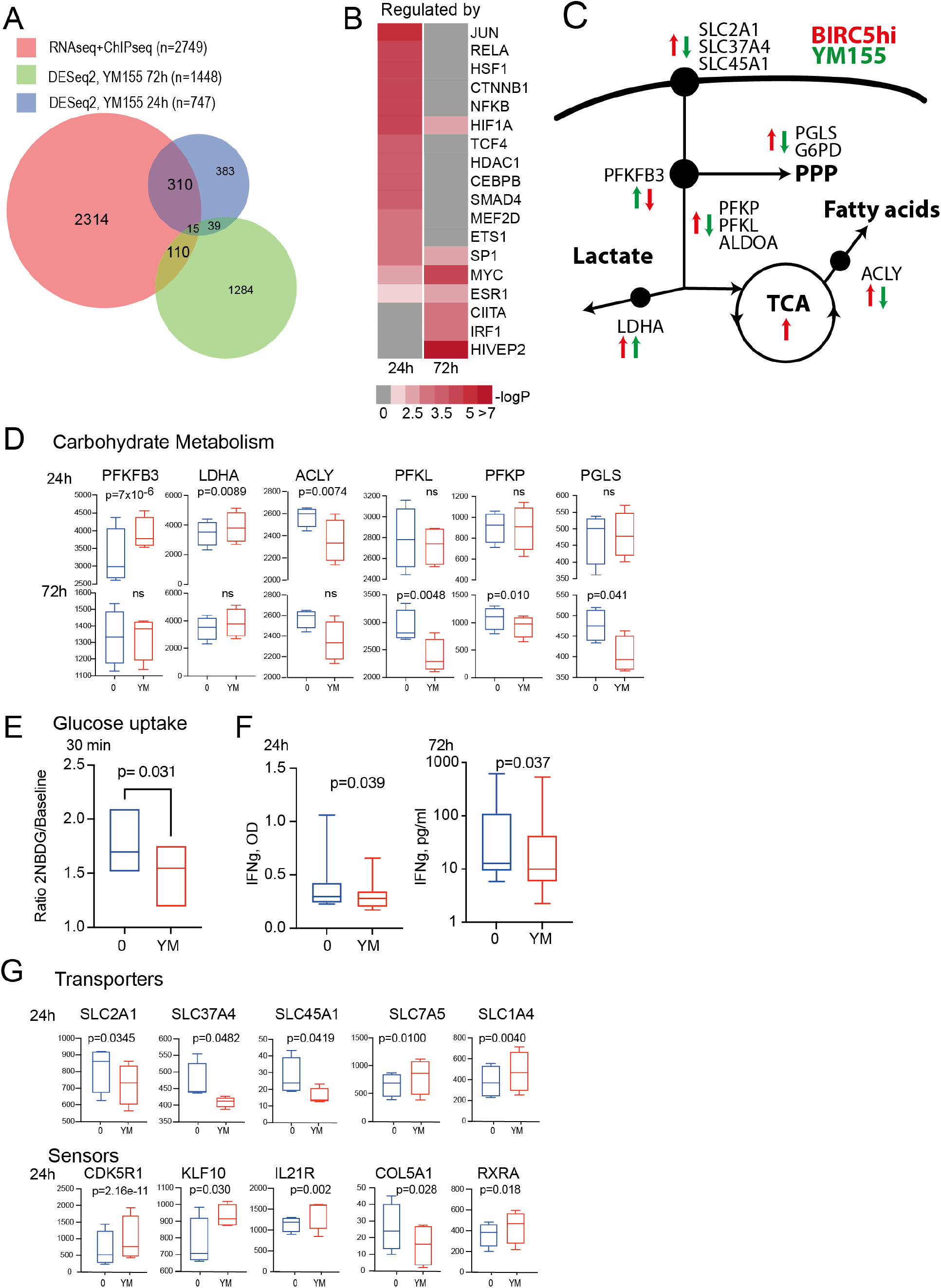
Survivin controls glycolysis through the phosphofructokinase metabolic axis. RNAseq of CD4^+^ T cells (n=4) treated with aCD3 and survivin inhibitor YM155 (0 and 10 nM) for 24h or 72h, activated with IFNγ during last 2h. P-values were obtained by the paired DESeq2. **A**. Venn diagram of protein-coding genes expressed in CD4^+^ cells, annotated to survivin peaks, and DEG in YM155-treated cells. **B**. Heatmap of upstream regulators of DEGs annotated by TRUSST database. **C**. Schematic illustration of glucose metabolism. Arrows indicate DEGs in BIRC5^hi^ cells (red) and in YM155-treated cells (green). **D**. Box plots of normalized RNAseq values of glycolytic enzymes. **E**. Box plots of 2NBD-glucose uptake by YM155-treated CD4^+^ cells, normalized to baseline. **F**. Box plots of IFNγ production in supernatants of aCD3/YM155-treated cells. P-values are obtained by the paired Wilcoxon’s sign rank test. **G**. Box plots of the expression of sugar transporters and sensors by normalized RNAseq values.

Next, we performed a detailed analysis of the enzymes involved in cellular glucose utilization (Figure 3C). RNAseq analysis of YM155-treated CD4^+^ cells revealed a rapid increase in the expression of mRNA coding for *PFKFB3*, the rate-limiting enzyme of pyruvate formation, and *LDHA*, which catalyses pyruvate conversion into lactate (Figure 3D). The reduced expression of mRNA coding for *PGLS* and *ACLY* (Figure 3D) indicates downregulation of the pentose phosphate pathway and fatty acid metabolism, occurred later. These observed effects of survivin inhibition *in vitro* are in line with the alterations in carbohydrate metabolism and glycolysis observed in the IFNγ-producing CD4^+^ cells from RA patients (Figure 1F).

To assess the role of survivin in the regulation of glucose uptake by CD4^+^ cells, we measured accumulation of the fluorescent dye 2NBD-glucose in CD4^+^ cell activated with aCD3/IFNγ. We found that YM155-treated CD4 cells had a significantly reduced uptake of 2NBD-glucose compared to sham-treated cells (Figure 3E). RNA-seq analysis of YM155-treated CD4^+^ cells demonstrated a significant decrease in the expression of the main sugar transporters *SLC2A1*, coding for GLUT1, glucose-6-phosphate transporter *SLC37A4* and H^+^ sugar transporter *SLC45A1*, while the expression of amino acid transporters *SLC7A5* and *SLC1A4* required for cell remodelling, and the expression of sugar sensors *CDK5R1, KLF10, IL21R* and *RXRA* was increased (Figure 3G).

### Inhibition of survivin activates TGF-beta/SMAD3 signalling and promotes memory-like transition of CD4^+^T cells

The attenuation of glucose metabolism resulted in a significant reduction of IFNγ production by YM155-treated CD4^+^ cells for 24h and 72h (Figure 3F). Analysing DEGs in GSEA, we found a significant limitation of IFNγ-dependent processes (Supplementary Figure 3Sb). After 24h, this included canonical ISG that control the clonal expansion receptors *IL2RA, SLAMF7* and *IL10RA*, and cytotoxic *PRF1*, joint homing receptors *CX3CR1, ITGB3, ICAM2*, and cytokines *CXCL8* and *IL1β* (Figure 4A). After 72h, the downregulation of ISGs was more pronounced and involved multiple IRF1-dependent genes such as *SOCS1, SOCS4*, and MHC class I, and canonical ISGs related to autoimmunity *IFNAR1, IFI35, IFITM2*, ubiquitin-like *ISG15* and *ISG20, LY6E, OAS1, ODF3B, RSAD2, LAMP3* and *GAS6*, also *IRF7 and IRF2BPL* (Figure 4B).

**Figure 4.**
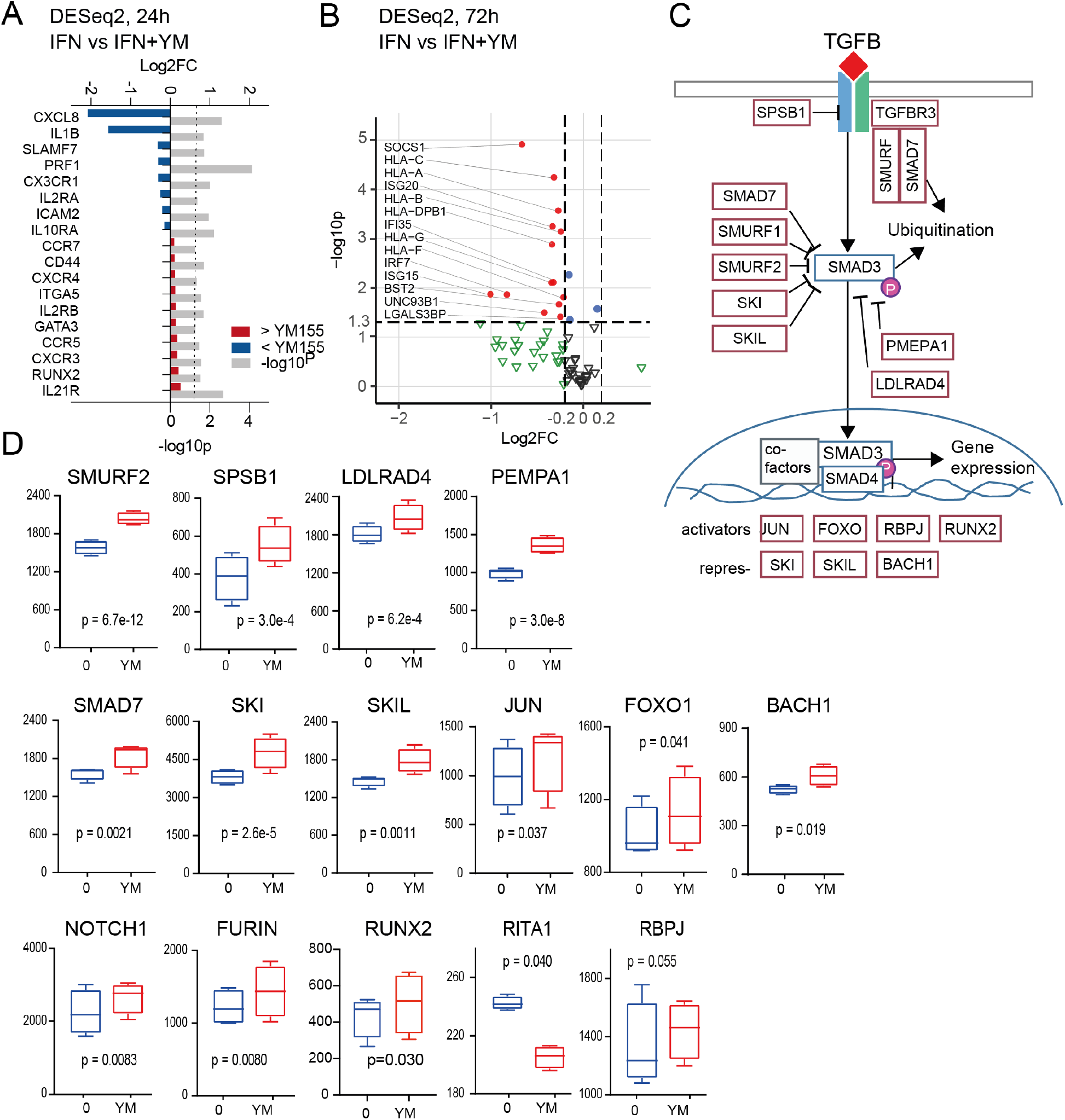
Inhibition of survivin activates TGFβ/SMAD3 signalling. RNAseq of CD4^+^ T cells (n=4) treated with aCD3 and survivin inhibitor YM155 (0 and 10 nM) for 24h or 72h, activated with IFNγ during last 2h. P-values were obtained by the paired DESeq2. **A**. Forest plot of the enrichment and p-values of the IFN-sensitive DEGs in YM155-treated cells for 24h. **B**. Volcano plot of the IFN-sensitive DEGs in YM155-treated cells for 72h. Clinically relevant IFN-sensitive genes are indicated by red dots. **C**. Schematic picture of the TGFβ/SMAD signalling pathway. **D**. Box plots of the key regulators of TGFβ signalling in YM155-treated cells for 24h. **E**. Box plots of the proteins of NOTCH signalling in YM155-treated cells for 24h.

Given that TGFβ-dependent mechanisms often counteract pro-inflammatory and cytotoxic properties of IFNγ (*50, 51*). We next assessed the effects of survivin inhibition on the expression of genes coding for the transcriptional regulators within the canonical TGFβ-pathway (Figure 4C). Among the top 77 DEGs (median expression level by basemean 546.4, unadjusted p-value below 0.005. Supplementary Figure 4S), we found a significant increase in the expression of a) E3 ubiquitin ligases *SMURF2, SPSB1*, and *SIAH3, LDLRAD4* and *PMEPA1* that facilitate proteolysis required for T cell reprogramming; b) *SMAD7* and its co-repressors *SKI* and *SKIL* that physically interact with the receptor-activated SMADs; c) the chromatin binding co-factors to SMAD3, *JUN, FOXO1* and *BACH1* (Figure 4D).

In addition, increased mRNA levels of *FOXO1* and *NOTCH1*, furin-like convertase (*FURIN*) that cleaves Notch protein to initiate its trafficking to cell membrane, reduced expression of Notch signalling repressor *RITA1*, together with moderately higher expression of *RBPJ* (Figure 4E) show that survivin inhibition induced the activation of NOTCH1 and FOXO1 pathways, that are controlled by the glycolytic enzymes PFKFB3 (*52, 53*) and LDHA (*54*), respectively. Additionally, the inhibition of survivin resulted in a significant upregulation of TFs RUNX2 and GATA3, which supported the development of the memory-like phenotype. Consequently, CD4 cells acquired higher transcription of the surface receptors *CD44, IL21R, ITGA5, CXCR3* downstream of NOTCH1, and the FOXO1 target genes including *IL2RB, CCR5, CCR7*, and *CXCR4* (Figure 4A). Taken together, these results indicate that the up-regulation of PFKFB3 and inhibition of the IFNγ program by survivin inhibition enables the maintenance of TGFβ signal and promotes the transcriptional activity of NOTCH and FOXO1 required for transition from effector to memory-like phenotype of CD4^+^ cells

### Survivin-ChIP peaks contain IRF-binding motif and survivin co-precipitates with IRF1 and SMAD3

To characterize the survivin bound chromatin sequences, we performed a motif analysis of genomic regions covered by the survivin-ChIP peaks. The analysis for the *de novo* motif prediction in the survivin-ChIP peaks was done using JASPAR database of TF binding sites in 4 independent ChIPseq replicates. It revealed a significant enrichment with IRF-binding motifs in all the samples. Among the IRF motifs dominated those suitable for binding of IRF1 and IRF8, both containing the conserved GAAA repeat (Figure 5A). Further analysis of the survivin-ChIP peak sequences revealed the enrichment of composite motifs bZIP/AP1:IRF (AICE motif, GAAAnnnTGAc/gTCA) and ETS/SPI1:IRF (EICE motif, GGAAnnGAAA). Additionally, we observed the co-localization of AICE and EICE motifs within a single survivin-ChIP peak and the presence of multiple binding sites for these motifs. The ISRE binding motif (GRAASTGAAAST), suitable for simultaneous binding of two IRFs, was infrequent, but it was still significantly enriched within the survivin-ChIP peaks compared to the whole genome (Figure 5A).

**Figure 5.**
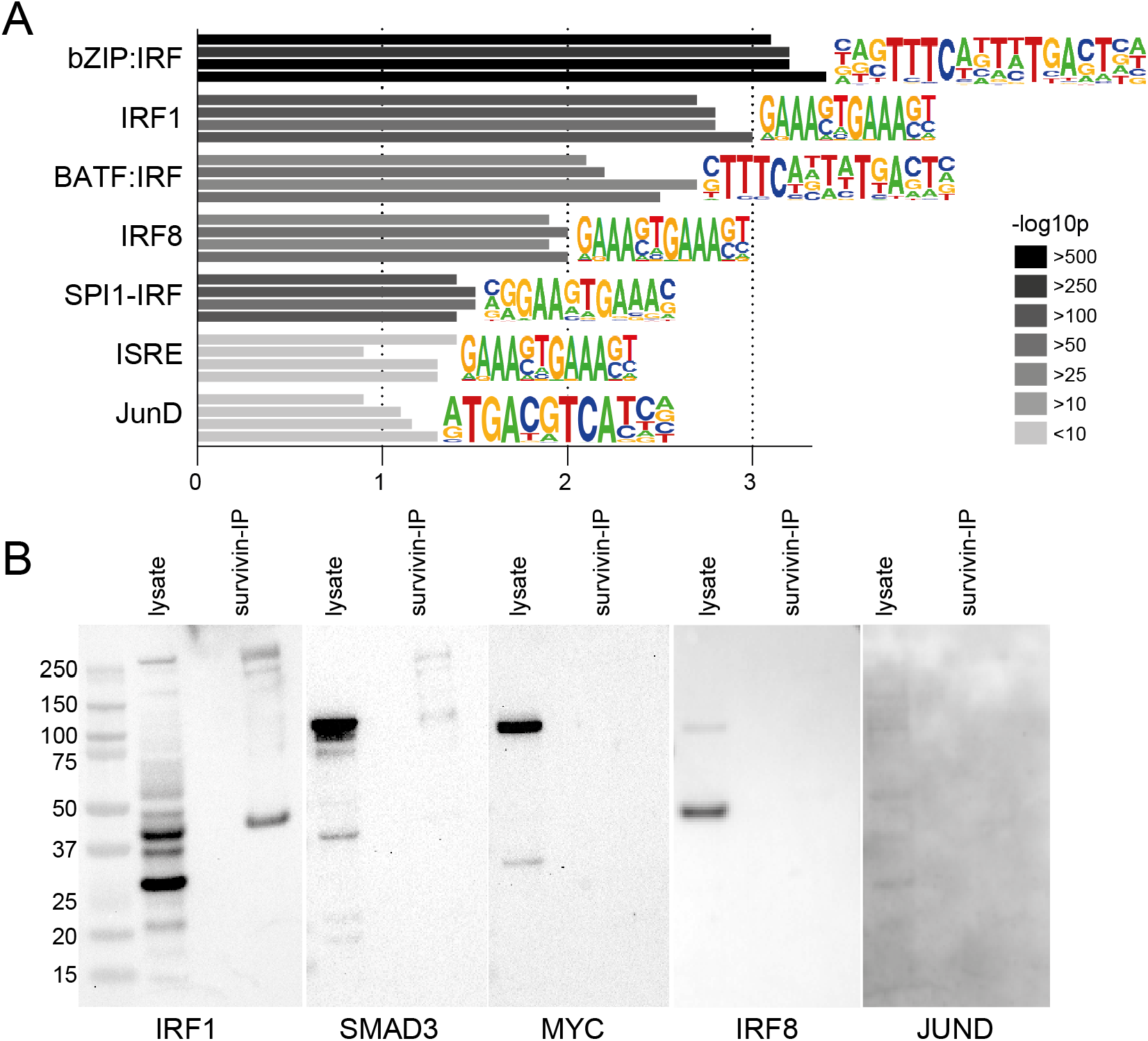
Survivin binding to IRF1 and SMAD3 recruits it to the chromatin. Survivin-ChIPseq was performed from CD4^+^ T cells (n=12) pooled into 4 independent samples. Survivin-ChIP peaks (enrichment against input, p<10^−5^) were mapped to hg38 human genome and analysed by HOMER program. *De novo* motif search was done in JASPAR database. **A**. Bar plots of the enrichment and significance for IFNγ-relevant DNA motifs in survivin-ChIP peaks. **B**. Western blots of THP1 cell lysate before and after affinity immunoprecipitation with survivin, stained for IRF1, SMAD3, c-MYC, IRF8 and JUND.

To provide the evidence of physical interaction between survivin and the predicted partners, we performed survivin co-immunoprecipitation using anti-survivin antibodies on lysate of THP1 cells followed by affinity isolation, protein denaturation and separation by electrophoresis. Western blotting of the membrane with specific antibodies revealed the presence of IRF1 and SMAD3 precipitating together with survivin (Figure 5B). However, neither IRF8, nor JUN, JUND could be visualised by western blotting in the survivin precipitated material from two independent experiments.

Taken together, these findings demonstrate that survivin is recruited to the chromatin containing the sequence specific IRF motifs through its binding to IRF1 and SMAD3. This provides the molecular evidence for a critical role of survivin in IFNγ-signalling observed in the functional experiments (Figures 3 and 4).

### IRF1 and SMAD3 are close partners of survivin in regulation of gene transcription

Since we anticipated location of the gene expression associated survivin-ChIP peaks in the *cis*-REs of the candidate genes, we prepared the complete list of 969 REs paired to the top 77 protein-coding DEGs (Supplementary Figure 4S) using the likelihood score for the enhancer-gene pairing (*55*). Among those, 117 REs contained survivin-ChIP peaks in 0-10kb vicinity. The remaining 852 REs had no survivin-ChIP peaks (Figure 6A,B). The characteristics of the survivin-containing REs were not different from the remaining REs with respect to GH-score, length/size of REs and distance to TSS (Supplementary figure 6S).

**Figure 6.**
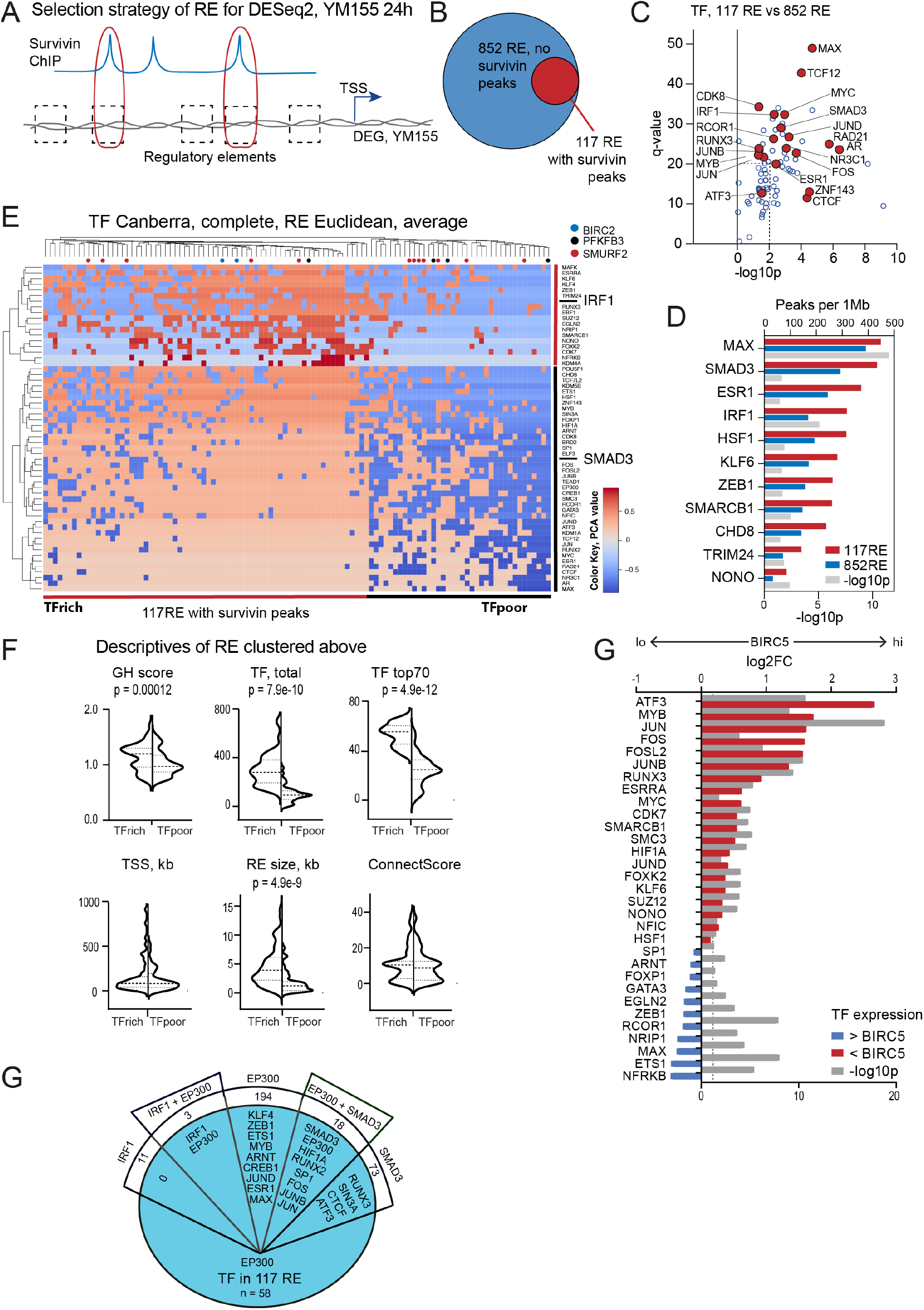
Predicted survivin partners in gene regulation. **A**. Selection of regulatory elements (RE) paired to protein-coding DEGs. **B**. Venn diagram of all REs (n=969) and survivin-containing REs (0-10kb flanks, n=117). **C**. Scatter plot of TFs enriched within survivin-containing REs against remaining REs (X-axis) and the genome (Y-axis). TFs present in >75% of 117 REs are indicated. **D**. Forest plot of TF density by ChIPseq peaks within survivin-containing REs (blue), remaining REs (open) and significance (red) between these two groups. **E**. Heatmap of principal component analysis of TFs within each survivin-containing RE. Clustering of REs by Euclidean distance, and of TFs by Canberra distance. Only TFs expressed in CD4^+^ cells (n=58) are used. **F**. Violin plots of specific characteristics of TF-rich and TF-poor survivin-containing REs. P-values were obtained by unpaired Student’s *t-*test. **G**. Protein-protein interaction by the BioGrid of IRF1, SMAD3 and EP300. **H**. Forest plot of the enrichment and p-values of TFs within survivin-containing REs between BIRC5^hi^ and BIRC5^lo^ CD4^+^ cells (RA patients, n=24). RNA-seq were analysed by DESeq2.

Among TFs co-localized with survivin-ChIP peaks (10% overlap, 0kb flanks) (Figure 2D), we identified 70 TFs with ChIPSeq peaks significantly more prevalent in the survivin-containing REs compared to the genome and to the remaining REs (all, p<10^−3^), 58 of those TF were expressed in CD4^+^ cells (Figure 6C). Analysis of density distribution of the TFs within the survivin-containing REs identified SMAD3 and IRF1 among the most frequent and densely present TFs compared with the remaining REs paired to the DEGs (Figure 6D).

To study internal relationship between the identified TFs, we analysed distribution and clustering of TFs across the REs of these genes. Principal component analysis followed by unsupervised clustering of those survivin-containing REs (Figure 6E) did not reveal any gene-specific clustering of REs to occur. The REs clustered by total density of TFs and further by association of TFs with IRF1 or SMAD3 (Figure 6E), which suggested engagement of survivin in TF complexes with distinct protein composition. The REs densely populated by TFs were significantly longer, which made them appropriate for accommodation of bulky complexes of diverse TF (Figure 6F).

Analysis of protein-protein interactions of human IRF1 and SMAD3 via the BioGrid database provided no evidence for a direct interaction between these proteins. We identified histone acetyltransferase EP300 and glycogen synthetase kinase GSK3B as the only common interactors of IRF1 and SMAD3. EP300/CREB1 complex that interacts and recruits TF to distant enhancers was present among the top 58 TF colocalized with survivin in the survivin-containing REs (Figure 6E,G) and could explain clustering of the REs by density of TFs observed in the PCA. Notably, a remarkable number of the top TFs were interacting with either IRF1, SMAD3 or EP300. Some of the proteins were differentially expressed in *BIRC5*^hi^CD4^+^ cells of RA patients (Figure 6H) and/or were sensitive to survivin inhibition by YM155 in CD4^+^ cells (Supplementary Figure S5C,D).

### Survivin exerts specific pattern of transcriptional regulation

To explore the mode of survivin specific transcriptional regulation, we analysed the chromatin regions containing the genes and REs highly sensitive to survivin inhibition and critical for the regulation of glycolysis in the IFNγ-producing CD4^+^ cells by survivin.

*PFKFB3* appeared as the main target of the survivin-dependent metabolic effects in CD4^+^ cells. We identified 4 survivin-ChIP peaks associated with 5 high-scored REs paired to *PFKFB3* (Figure 7A). Three REs were clustered ∼20kb upstream of *PFKFB3* and two additional REs were detected 90kb and 100kb downstream. We could see that both the upstream and downstream REs contained ChIPSeq peaks for IRF1 and SMAD3 grouped together with the survivin-ChIP peaks (Figure 6E). The chromatin regions containing the REs were functionally connected to each other and to the *PFKFB3* promoter according to the integrated annotation of RE connections in the GeneHancer database. In CD4^+^ cells, the cluster of upstream REs partly covers the active *PFKFB3* promoter, while the downstream REs are located in the repressed/poised region of chromatin. The importance of these REs for *PFKFB3* expression and their role in T cell mediated disorders is supported by GWAS studies. In the vicinity of the upstream REs, there was a dense accumulation of SNPs associated with latent autoimmune diabetes, IL2RA levels and thyroiditis. SNPs near the downstream REs were annotated to T1D, RA, thyroiditis and celiac disease (Figure 7A. Supplementary table 7S).

**Figure 7.**
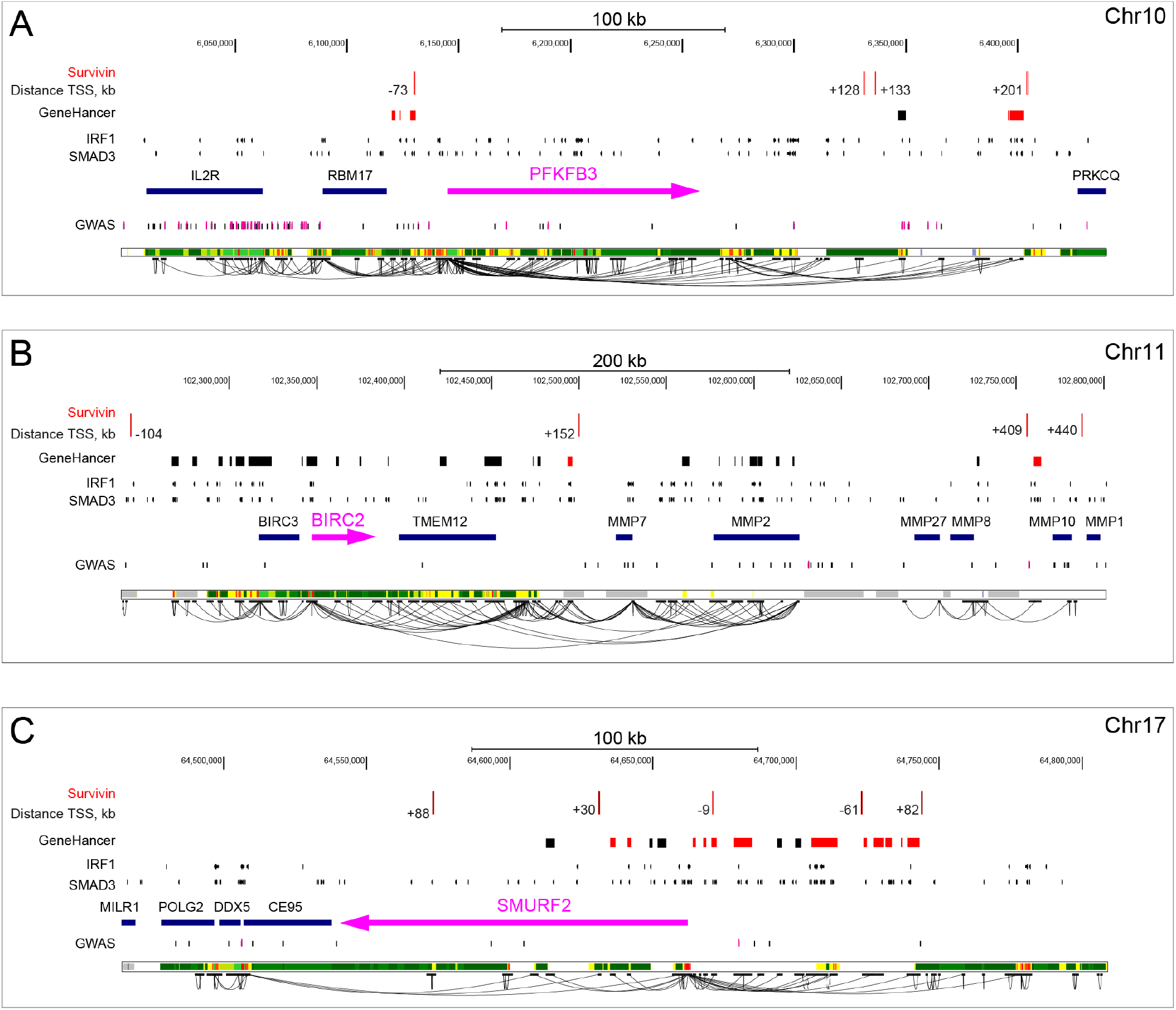
Regulatory pattern of survivin bound to the chromatin in CD4^+^ T cells. Genomic maps of the gene of *PFKFB3* **(A)**, *BIRC2* **(B)** and *SMURF2* **(C)**. Position of the canonic gene transcript is shown in magenta, arrow indicates the transcription orientation. Survivin-ChIP peaks positions are shown by red dashes at the top of each loci. Distance to TSS is indicated. Regulatory elements (REs) paired to the gene by GeneHancer are indicated by boxes. Integrated annotation of RE connections by the GeneHancer are shown by solid lines. Red boxes indicate REs <10kb to survivin peak. Position of ChIPSeq peaks for IRF1 and SMAD3 by ReMap2020 are shown by dashes. Positions of GWAS SNPs associated with metabolic and autoimmune triads are indicated by dashes according to NHGRI GWAS catalog. Functional chromatin segmentation for activated CD4^+^ cells (RoadMap ChromHMM. E042:CD4^+^CD25^-^ IL17^+^_PMA-Ionomyc_stimulated_Th17_Primary_Cells) is shown at the bottom of each map. Green blocks cover actively transcribed areas; yellow blocks depict enhancers; red blocks depict active promoters; grey blocks depict repressed (dark grey) and poised (light grey) areas.

Another gene that was significantly upregulated by survivin inhibition was *BIRC2*, which potentially contributes to the regulation of glycolysis-dependent cell survival and growth (*56, 57*). Figure 7B shows an extended region of ∼550 kb containing 26 REs paired to and surrounding *BIRC2*. Among those REs, only 2 candidate REs are associated with 3 survivin-ChIP peaks. Both REs are located downstream of *BIRC2*, ∼100 kb and ∼400kb from TSS, respectively. Despite the distant location, the REs are paired to *BIRC2* with the GH-score of 1.56 and 10.95, respectively, and illustrate the long-range interaction between the survivin-containing REs and the *BIRC2* promoter region. Both REs contained multiple IRF1 and SMAD3 ChIPseq peaks. According to the functional chromatin segmentation in CD4^+^ cells, both REs are located within the repressed/poised chromatin and putatively activated after survivin inhibition. A search for the GWAS traits in the extended *BIRC2* region identified only two SNPs with connection to autoimmunity. The one close to the distant survivin-containing RE was associated with multiple sclerosis.

Yet another gene that regulates cell cycle stability and aerobic glycolysis is *SMURF2. SMURF2* showed the highest expression change after survivin inhibition in our study (Supplementary Figure 4S). The pattern of *SMURF2* REs is different. The *SMURF2* REs form a dense cluster of 25 REs adjacent to each other that spans the region of ∼100kb upstream of TSS and covers the gene start (Figure 7C). Of these REs, 12 are close to 4 survivin peaks and interspersed 5 other REs that were not associated with survivin. An additional survivin peak is located within *SMURF2*, outside of any RE. Only one of 12 survivin-associated REs is located in the active chromatin region ∼50kb upstream of the TSS. The remaining REs are found in the repressed/poised chromatin. By similarity with *PFKFB3* and *BIRC2* loci, these REs are predicted to have alternating functional state controlled by survivin. The *SMURF2* paired REs vary in their properties and represent the whole spectrum of qualities with respect to size/length, density of TFs, and presence of survivin partners IRF1 and SMAD3 (Figure 6E). In contrast to the *PFKFB3* and *BIRC2* loci, the upstream region of *SMURF2* contains no other genes, but is rich with clustered REs. All but one survivin-associated REs in the *SMURF2* locus were annotated to the chromatin inactive in CD4^+^ cells. According to functional chromatin annotation of the NIH Roadmap Project, activity of those REs is coordinated across the cells presenting a higher order regulatory unit. Therefore, a cooperative activation of those clustered REs could be expected after survivin inhibition and presents a plausible mechanism for the pronounced upregulation of *SMURF2* expression demonstrated in this study. Such functional associations for complex, RE-rich *SMURF2* locus were not supported by currently available SNP-based GWAS data.

Taken together, the analysis of the survivin-containing REs paired to the DEGs sensitive to survivin inhibition revealed common features of transcriptional regulation. These features include a) long-range interactions between the survivin-containing RE and the promoters of target genes, b) location of the survivin-containing RE among several REs clustered into regulatory modules, and c) location of those REs on the repressed/poised chromatin. The activation of gene expression after survivin depletion highlights the role of survivin as the essential transcriptional repressor for these chromatin regions.

## Discussion

This study demonstrates that the oncoprotein survivin is an important regulator of transcription in CD4 cells that is required to maintain the phenotype of IFNγ-producing Th1 cells. We show that survivin provides a critical link between the increased anabolic activity of the effector Th1 cells and transcriptional control of the genes responsible for glycolysis, sugar sensing and transport. Analysing the predicted regulatory pattern of survivin on the chromatin, we identify common features of the survivin-binding DNA regions with clusters of REs located at a significant distance from the target gene, often in the poised chromatin.

We demonstrate that survivin position on chromatin is defined by its physical interaction with transcription factors IRF1 and SMAD3, which could explain its recruitment to the DNA regions containing IRF composite motifs and to the REs regulated by IRF1 and SMAD3. Concordantly, independent integration analysis of the publicly available human ChIPseq datasets predicted colocalization of survivin with IRF1 and SMAD3 and identified them among the most frequent and densely present TFs allocated to the survivin-containing REs paired to DEGs. Co-precipitation of survivin with IRF1 and SMAD3 provided strong evidence for the close interaction between these proteins while the YM155-inhibition studies pointed to the role of IRF1 and SMAD3 in competing cell programs. On the functional level, the effects of IRF1 and SMAD3 have been mostly studied through binding to the promoter regions of non-identical genes that often regulate opposing transcriptional programs (*58, 59*). IRF1 and SMAD3 were identified among the TFs binding to the distant regulatory regions and facilitating chromatin accessibility for other proteins (*60, 61*). SMAD3 has been attributed chromatin remodelling properties by coordination of super-enhancers (*62-64*). Thus, we propose that survivin functions as a bridge between the transcriptional programs governed by these two proteins.

Analysis of IRF1 and SMAD3 protein interactions identified histone acetyltransferase EP300 as a common interactor of human IRF1 and SMAD3. Notably, proteins of the EP300/CREB1 complex were significantly enriched in the survivin-containing REs paired to DEGs sensitive to YM155 treatment in their expression. Remarkably, EP300 integrates the immune processes initiated by IFNγ- and TGFβ-signalling. EP300 is activated by IFNγ through STAT1 and IRF1 dependent mechanism (*65*). Acetyltransferase activity of EP300 and its interaction with SMAD3 promotes the TGFβ-dependent effects (*66*).

The REs paired to DEGs were their predominantly located in the inactive/poised regions in CD4^+^ cells. EP300/CREB1 complex acts as transcriptional activator and participates in RNA polymerase II dependent recruitment of TF to the chromatin regulatory regions including distant enhancers. We propose that survivin inhibition converts these regions from inactive to active by enabling the interaction between EP300 and SMAD3. The subsequent activation of SMAD3 associated proteins FOXO1, RUNX2 and NOTCH1 and their transcriptional programs required for acquisition of the memory-like CD4^+^ phenotype. In this scenario, survivin functions as a guardian for the functional chromatin state that translates into the change of CD4^+^ cell phenotype. Activation of EP300/CREB1 complex is mediated by glucose (*67, 68*), which provides an additional support to its connection with survivin.

This study demonstrates that repression of the *PFKFB3* gene is central for the survivin-dependent metabolic effects in CD4^+^ cells. Acting in synergy with IRF1, survivin located on the REs paired to *PFKFB3* represses its activity. Expression of *PFKFB3* has a strong negative correlation with survivin levels in freshly isolated CD4^+^ cells. Production of PFKFB3 and the conventional aerobic glycolysis through the tricarboxylic acid cycle is restored early after survivin inhibition. This survivin-dependent switch in glucose utilisation forwards the logical connection between survivin and IRF1-dependent effector function of CD4^+^ cells and provides an important interventional input in regulation of autoimmunity (*17, 21*). Activation of SMAD3 has been reported as a functional consequence of the restored PFKFB3-dependent glycolytic activity (*69*). This notion finds new and stronger support in this study emphasizing the critical role of survivin in transcriptional control of PFKFB3. The chromatin locus of the *PFKFB3* gene and the paired REs contain numerous SNPs identified by GWAS studies on autoimmune diseases and the clinical traits associated with these SNPs point to a strong link between the *PFKFB3* genomic region and metabolic and autoimmune conditions. Identification of epigenetic mechanism connecting SNPs with the associated phenotype remains to be studied further.

The reset of TGFβ/SMAD3 signalling by survivin inhibition caused pronounced upregulation in transcription of the E3 ubiquitin ligases group of genes *SMURF2, SMAD7* and its co-repressors *SKI* and *SKIL* required for remodelling of cell function. *SMURF2* was the top gene activated after survivin inhibition in CD4^+^ T cells. In complex with SMAD3, SMURF2 represses the function of the central regulator of cell metabolism MYC with inevitable consequences for glucose metabolism and profound impairment of cell proliferation (*70, 71*). BIRC2 is another protein with ubiquitinase activity upregulated by survivin inhibition. Ubiquitinating MYC cofactor MAD1 (*56, 57*) BIRC2 accelerates MYC effects on glucose metabolism and provides a strong positive feedback to the metabolic processes affected by survivin inhibition.

The results of this study are produced in primary CD4^+^ cells from RA patients and healthy subjects. However, we believe that these findings make an important contribution to the understanding of general molecular mechanisms of autoimmunity. Survivin interaction with IRF1 maintain expression of the ISGs with established clinical relevance in several autoimmune diseases including RA(*72*), SLE (*73*) and Sjögren’s syndrome (*74*). Survivin dependent combination of the reverted glycolysis with high sugar uptake provides optimal energy supply required for the effector function of CD4^+^ cells. This supports the process of persistent chronic inflammation.

Taken together, the study presents experimental evidence for the fundamental role of survivin in the regulation of the IRF1 dependent transcription in CD4^+^ T cells. Through physical interaction with IRF1, survivin is recruited to the chromatin regulatory elements to repress the expression of the rate-limiting glycolytic enzyme PFKFB3 and facilitate glucose utilization through the pentose phosphate pathway. Inhibition of survivin restores the metabolic profile of CD4^+^ cells and reduced their effector activity. This function of survivin sheds new light on the regulation of the balance between IFNγ- and TGFβ-dependent processes. Pharmacological intervention that selectively targets these molecular interactions of survivin could present an attractive approach to improve control of IFNγ-dependent autoimmunity.

## Material and Methods

### Patients

Blood samples of 46 RA patients and 7 healthy female individuals were collected at the Rheumatology Clinic, Sahlgrenska Hospital, Gothenburg. Clinical characteristics of the patients are shown in Supplementary table 8S. All RA patients fulfilled the EULAR/ACR classification criteria (*75*) and gave their written informed consent prior to the blood sampling. The study was approved by the Swedish Ethical Evaluation Board (659-2011) and was performed in accordance with the Declaration of Helsinki. The trial is registered at ClinicalTrials.gov with ID NCT03449589.

### Isolation and stimulation of CD4^+^ cells

Human PBMC were isolated from venous peripheral blood using density gradient separation on Lymphoprep (Axis-Shield PoC As, Norway). CD4^+^ cells were isolated using positive selection (Invitrogen, 11331D), and cultured (1.25×10^6^ cells/ml) in complete RPMI-medium supplemented with concanavalin A (ConA, 0.625μg/ml, Sigma-Aldrich), and LPS (5μg/ml, Sigma-Aldrich), for 24 or 72h. In the inhibition experiments, CD4^+^ cells were cultured in aCD3 precoated wells (0.5μg/ml, OKT3, Sigma-Aldrich), in RPMI-medium supplemented with survivin inhibitor YM155 (0 or 10nM, Selleck chemicals, Houston, TX) for 24h or 72h. Stimulated with recombinant IFNγ (50ng/ml, Peprotech, Cranbury, NJ, USA) during last 2h.

### Flow cytometry analysis

Freshly isolated PBMC were stimulated overnight with ConA 0.625 *µ*g/ml, harvested and stained for flow cytometry as described (*37*) using antibodies to human surface antigens: CD4-APCH7 (SK3, BD Biosciences Franklin Lakes, NJ, USA), CD8-PerCP (SK1, BD), CD19-V500 (H1B19, BioLegend), CD62L-PECy7 (DREG-56, BD), CD27-APC (L128, BD) and CD45RA-BV421 (HI100, BioLegend). Cells were then fixed and permeabilized with Cytofix-Cytoperm permeabilization kit (BD) and stained with anti-Survivin (91630, R&D Systems) and isotype control (mouse IgG1κ, both R&D Systems, Minneapolis, MN, USA). The cells were analysed in FACSCantoII (BD) and data was evaluated in FlowJo software (BD, v.10.7) using appropriate FMO controls.

### Chromatin immunoprecipitation, library preparation and sequencing

CD4^+^ cells were isolated from 12 females stimulated with ConA+LPS for 72h and pooled in 4 independent samples for chromatin purification. CD4^+^ cells were cross-linked and lysed according to EpiTect ChIP OneDay kit (Qiagen). After sonication to shear the chromatin, cellular debris was removed by pelleting. After pre-clearing, 1% of the sample was saved as an input fraction and used as background nonspecific chromatin binding. The pre-cleared chromatin was incubated with 2μg anti-Survivin (*76*) (10811, SantaCruz Biotechnology). The immune complexes were washed, the cross-links reversed, and the DNA purified according to the EpiTect ChIP OneDay kit (Qiagen). Resulting purified DNA was quality controlled using TapeStation (Agilent, Santa Clara, CA, USA). DNA libraries were prepared using ThruPLEX (Rubicon). and sequenced using the Hiseq2000 (Illumina) following the manufacturer’s protocols. Bcl-files were then converted and demultiplexed to fastq using the bcl2fastq program from Illumina.

### Transcriptional sequencing (RNAseq)

RNA from CD4^+^ cell cultures was prepared using the Norgen Total micro mRNA kit (Norgen, Ontario, Canada). Quality control was done by Bioanalyzer RNA6000 Pico on Agilent2100 (Agilent, St.Clara, CA, USA). Deep sequencing was done by RNAseq (Hiseq2000, Illumina) at the LifeScience Laboratory, Huddinge, Sweden. Raw sequence data were obtained in Bcl-files and converted into fastq text format using the bcl2fastq program from Illumina. Validation of RNAseq was performed using qRT-PCR as described below.

### RNA-seq analysis

Mapping of transcripts was done using Genome UCSC annotation set for hg38 human genome assembly. Analysis was performed using the core Bioconductor packages in R-studio v.3.6.3. Differentially expressed genes (DEGs) were identified using DESeq2 (v.1.26.0) with Benjamini-Hochberg adjustment for multiple testing. Volcano plots were prepared with EnhancedVolcano (v.1.4.0). Correlation analysis was done with Hmisc (v.4.5), the correlation heatmap was built using Corrplot (v.0.85). Clustering of RNA-seq data was performed using Spearman correlation for distance (factoextra, v.1.0.7) and hierarchical clustering with WardD2.

### Conventional qPCR

RNA was isolated with Total RNA purification kit (Norgen, #17200). Concentration and quality of the RNA were evaluated with a NanoDrop spectrophotometer (ThermoFisher Scientific) and Experion electrophoresis system (Bio-Rad Laboratories). RNA (400 ng) was used for cDNA synthesis using High-Capacity cDNA Reverse Transcription Kit (Applied Biosystems, Foster City, CA, USA). Real-time amplification was performed with RT2 SYBR Green qPCR Mastermix (Qiagen) using a ViiA 7 Real-Time PCR System (ThermoFisher Scientific) as described(*44*). Primers used are shown in Supplementary Table 9S. The expression was calculated by the ddCt method.

### Affinity immunoprecipitation and western blotting

Human monocyte leukemia cell line (THP-1) was cultured at a density of 3-10×10^5^ cells/mL in RPMI1640 medium supplemented with 10% FBS at 37°C in a humidified atmosphere of 5% CO2. Cells were lysed in IP Lysis Buffer (87787, Pierce™) supplemented with protease inhibitors (Complete mini, Roche), and IP was performed with antibodies against survivin (RnD AF886) coupled to the Dynabeads Protein G Immunoprecipitation Kit (10007D, ThermoFisher Scientific), cross-linked with bis(sulfosuccinimidyl)suberate (A39266, Pierce™).

Western blots were performed by loading 30μg of total proteins from whole-cell lysates, and the immunoprecipitated (IP) material on NuPage 4–12% Bis–Tris gels (Novex). Proteins were transferred to polyvinylidene difluoride membranes (iBlot; Invitrogen). Membranes were blocked with TBST containing 3% BSA and incubated with antibodies against IRF1 (H-8, sc-74530), IRF8 (E-9, sc-365042), JUND (D-9, sc-271938), SMAD3 (38-Q, sc-101154), MAX (H-2, sc-8011), and MYC (9E10, sc-40) (all, Santa Cruz Biotechnology, USA) followed by peroxidase-conjugated anti-mouse antibodies (NA931, GE Healthcare, Chicago, IL). Bands were visualized using the ECL Select Western Blotting Detection Reagent (Amersham), and the ChemiDoc equipment with the Quantity One software (Bio-Rad Laboratories). Primary antibodies were used at 1:500 dilution, secondary at 1:4000.

### Cytokine measurement

Cytokine levels in supernatants of CD4^+^ cells were measured with sandwich enzyme-linked immune assay. Briefly, high-performance 384-well plates (Corning Plasticware, NY USA) were coated with capture antibody, blocked and developed according to the manufacturers’ instructions. IFNγ (PelikineM1933, detection limit 3pg/ml, Sanquin, Amsterdam, the Netherlands), IL10 (DY217B, detection limit 15pg/ml, RnD), IL9 (DY209, detection limit 1 pg/ml, RnD), IL13 (DY213, detection limit 50pg/ml, RnD), IL4 (DY204, detection limit 0.25 pg/ml, RnD) Developed plates were read in a SpectraMax340 Microplate reader (Molecular Devices, San Jose, CA, USA) and absolute protein levels were calculated after serial dilutions of the recombinant protein provided by the manufacturer.

### Glucose uptake assay

Glucose uptake was monitored using the fluorescent D-glucose derivate, 2-N-nitrobenz-2-oxa-1,3diazol-4-amino]-2 deoxy-D-glucose (2NBDG) (Abcam). CD4^+^ cells were cultured for 24h in anti-CD3 coated plates (0.5μg/ml) supplemented with IFNγ (50ng/ml) and YM155 (0 and 10 nM). Cells were starved in glucose-free RPMI-media for 2h and then supplemented with 2NBDG (100 *µ*M). 2NBDG uptake was registered after 30min using flow cytometry (Verse, BD) and quantified as a ratio between mean fluorescence intensity compared to baseline.

### ChIP-seq analysis

The fastq sequencing files were mapped to the human reference genome (hg38) using the STAR aligner (*77*) with default parameters apart from setting the alignIntronMax flag to 1 for end-to-end mapping. Quality control of the sequenced material was performed by FastQC tool using MultiQC v.0.9dev0 (Babraham Institute, Cambridge, U.K.). Peak calling was performed using the default parameters. Peaks were identified for survivin antibody immunoprecipitation fraction (IP) and unprocessed DNA (Input) and annotated by HOMER software (*78*) in standard mode to closest transcription start site (TSS) with no distance restriction. A set of peaks with enrichment versus surrounding region and input (adjusted p<10^−5^) was identified and quantified separately for each of sample. Peaks with overlapping localization by at least 1 nucleotide in several samples were merged and further on referred to as survivin-ChIP peaks. Peaks present in all samples were scored by number of tags of difference between IP and Input (average of these differences between samples). The HOMER program (findMotifsGenome.pl) with homer2 engine was used for the *de novo* motifs discovery and motif scanning. Most common *de novo* motifs were identified for each IP sample separately and searched detected motifs in the JASPAR database of human transcription factor binding sites (*79*). Each annotation motif was checked manually, and only highly aligned motifs were scored.

### Genome distribution of survivin-ChIP peaks

The whole interval set of survivin-ChIP peaks was compared with the set of functional genomic regions using Genome UCSC annotation hg38 (http://genome.ucsc.edu/, accessed 10apr2021). TSS were defined based on chromStart or chromEnd positions in GENCODE v36. Promoters were defined as regions 5kb upstream plus 1kb upstream of TSS annotated as above. The CTCF-binding sites were accessed according to ENSEMBL regulatory build (v103, 2020. http://www.ensembl.org/info/docs/funcgen/regulatory_build.html, accessed 01mars2021) (177376 elements). Insulator sites for all aggregated cells were defined according to ENCODE v5, 2020) (https://screen.encodeproject.org/, https://api.wenglab.org/screen_v13/fdownloads/GRCh38-ccREs.CTCF-only.bed file, accessed 01mars2021) (56766 elements). Enhancer selection was done using the integrated GeneHancer database (v4.4, https://www.genecards.org/GeneHancer_version_4-4, accessed 05jan2021. GH-score above 0.7).

For genomic interval datasets, including survivin-ChIP peaks and regulatory elements, the Table Browser for hg38 human genome assembly (http://genome.ucsc.edu/cgi-bin/hgTables) and Galaxy suite tools were applied (https://usegalaxy.org/) to estimate distances between nearest intervals, merging, overlapping, calculating genomic coverage and other standard procedures. Initial screening of genome wide distribution of survivin-ChIP peaks was carried out using the *cis*-regulatory annotation system (CEAS v0.9.8, accessed 01nov2020 via Cistrome Galaxy http://cistrome.org/ap/root). For enrichment analysis, total list of survivin-ChIP peaks and their fraction located within 100kb of the known genes was used. For pairwise distances estimation and statistical significance of pairwise interval overlaps for survivin-ChIP peaks with genome elements defined above, we used Bedtools suite (https://github.com/arq5x/bedtools2, accessed 01feb2021–15apr 2021). For each comparison the pairwise Fisher’s exact test was applied two-tailed. Comparison was based on initial survivin-ChIP peak positions as intervals and extended regions with 1kb, 10kb and 50kb flanks.

### Peak colocalization analysis with transcription regulators

To identify the transcription regulators in vicinity of survivin-ChIP peaks, we performed colocalization analysis of aggregated cell- and tissue-agnostic human ChIPseq datasets of 1034 transcriptional regulators using ReMap database (http://remap.univ-amu.fr/, accessed 15nov2020). Colocalization enrichment analysis was performed using ReMapEnrich R-script (https://github.com/remap-cisreg/ReMapEnrich, accessed 15nov2020). All comparisons were carried out using hg38 human genome assembly. Two-tailed p-values were estimated and normalized by Benjamini-Yekutielli, maximal allowed value of shuffled genomic regions for each dataset (n=15), kept on the same chromosome (shuffling genomic regions parameter byChrom=TRUE). Default fraction of minimal overlap for input and catalogue intervals was set to 10%. Additional analysis was performed with 30% and 50% overlap fraction for survivin-ChIP peaks. Bed interval files of survivin-ChIP peaks with 0 and 100kb flanks were prepared. The dataset with 0kb flanks was compared with the Universe sets of genomic regions, defined as 1Mb vicinity of the same ChIPseq peaks. For analysis of the regulatory chromatin paired with DEGs, input bedfiles were selected according to their distance from the genome region containing REs paired to DEGs and bedfiles for individual TFs downloaded from ReMap2020.

### Analysis for candidate partner transcription factors

The genome-wide analysis of the colocalized TFs was done for (1) survivin-ChIP peaks with 0kb flanks, minimal overlap fraction of 10%; (2) survivin-ChIP peaks with 100kb flanks in comparison to whole genome; (3) survivin-ChIP peaks with 0kb flanks in comparison to 1Mb vicinity to these peaks. TFs with statistically significant enrichment of overlaps (q-value < 0.05, number >100) were selected. Enriched TFs against the genomic background were identified within each RE using ReMap database, as described above. A subset of TFs enriched within the survivin-associated REs were calculated by chi-square test (chisq.test, R-studio) and FDR correction (R-studio). To explore the involvement of these TFs to regulation of DEGs, we prepared the presence matrix (1/0 type), excluded regions with 0 overlaps with top TFs, performed PCA analysis with singular value decomposition imputation. Hierarchical clustering of TFs was done with Canberra or Euclidean distances (prcomp, hclust, R-studio). FDR-adjusted p-values and a ratio between the survivin-associated and survivin-independent REs per 1Mb were estimated.

### Gene set enrichment analysis

Analysis of DEGs was performed in comparison to all protein-coding human genes (by default) using the Gene Set Enrichment Analysis (GSEA) (https://www.gsea-msigdb.org/gsea/index.jsp, accessed 15nov2020). TFs with significant overlap between ChIPseq and survivin-ChIP peaks were analysed in comparison to all 1034 transcriptional regulators within ReMap2020 (accessed 20nov2020). General functional categories for DEGs and TFs was annotated using the MetaScape service (https://metascape.org/gp/index.html, accessed 15nov2020). For the pathway analysis, Gene Ontology terms and TRRUST database were used. Pathways were selected by minimal gene overlap equal to 2, p-value <0.05 and minimal enrichment threshold of 1.3. Terms retrieved from the Gene Ontology Biological Processes(GO:BP) were grouped by medium term similarity of 0.7 and arranged on the semantic similarity scale coordinates using ReViGo service (http://revigo.irb.hr/, accessed 01dec2020).

### Genome-wide association data

Known genetic associations of the analysed regulatory regions of DEGs were examined using NHGRI’s collection of Genome-Wide Association Studies (http://genome.ucsc.edu/, accessed 01May2021). All published GWAS SNPs were included without p-value or ancestry filtering. A subset of relevant SNPs was selected by the keyword search for the trait for individual autoimmune disorders in the Table Browser(*80*).

### Chromatin functional segmentation data in CD4^+^ cells

Primary functional chromatin segmentation was accessed using NIH Roadmap Epigenomic Project data for activated CD4^+^ T cells (E042: CD4^+^_CD25^-^_IL17^+^_PMA/Ionomyc_stimulated_Th17_Primary_Cells) accessed via the Washington University Epigenomic Browser (http://epigenomegateway.wustl.edu/browser/roadmap/, accessed 01may2021). Default colour scheme was applied for chromatin segments of active enhancers, transcribed regions, repressed and poised loci.

## Supporting information

all supplementary figures and tables

## Author contributions

Conceiving the study (MB, KA, GK), collection of material (ME, KA, STS, ZE, MB), laboratory work (KA, ME, MJGB), statistical analysis (ME, KA, VC, NO, MB, AD), drafting the manuscript (MB, NO, MP, GK, ME). All authors contributed with interpretation of data, active discussions and feedback throughout the whole process of the manuscript preparation.

## Acknowledgements

This work has been funded by grants from the Swedish Research Council (MB, 521-2017-03025), the Röntgen-Ångström Cluster Framework of the Swedish Research Council (GK, 2015-06099), the Swedish Association against Rheumatism (MB, R-566961), the King Gustaf V:s 80-year Foundation (MB), the Regional agreement on medical training and clinical research between the Western Götaland county council and the University of Gothenburg (MB, ALFGBG-717681), the University of Gothenburg, Southern Älvsborg Hospital, Borås, Sweden (ZE).

## Supplementary table headings and figure legends

**Supplementary Figure S1**

A. Spearman correlation analysis of expression by normalized RNAseq values between *BIRC5* and Th1 signature genes

B. Spearman correlation analysis of expression by normalized RNAseq values between *PFKFB3*, and its ratio *PFKFB3/LDHA* and *PFKFB3/G6FD* with *BIRC5* and Th1 signature genes

**Supplementary Table S2**. The complete list of significantly enriched biological processes regulated by the proteins co-localized with survivin peaks (10% overlap). Functional annotation is done in MetaScape. Coordinates for the GO:BP correspond to the X and Y-axis of the semantic map in Figure 2F.

**Supplementary figure S3**.

A. Bar plot of immunologically relevant processes regulated by the proteins colocalized with survivin-ChIP peaks (10% overlap) identified in WikiPathway and KEGG.

B. Barcode plots of the IFNα and IFNγ signalling pathway for the DEG in YM155-treated CD4^+^ cells, by GSEA analysis

**Supplementary figure S4**

Heatmap of 77 protein-coding DEGs (RNASeq normalized value, basemean >0.5, DESeq2 unadjusted p-value <0.005) in YM155-treated CD4^+^ cells for 24h, last 2h stimulated with IFNγ (healthy, n=4).

**Supplementary figure 5S**

**A**. Forest plot of enrichment signifiance of bZIP family proteins in BIRC5^hi^ and BIRC5^lo^ CD4^+^ cells (blue, RA patients, n=24) and in YM155-treated CD4^+^ cells (red, healthy females, n=4). P-values are obtained by DESeq2. **B**. Box plots of normalized RNAseq values in BIRC5^hi^ and BIRC5^lo^ CD4^+^ cells (RA patients, n=24) and in YM155-treated CD4^+^ cells (healthy females, n=4). RNAseq were analysed by DESeq2. **C. D**. Forest plot of enrichment and p-values of IRF1 and SMAD3 interactors (BioGrid database, >1 physical evidence) in BIRC5^hi^ and BIRC5^lo^ CD4^+^ cells (red, RA patients, n=24). RNAseq were analysed by DESeq2.

**Supplementary Figure S6**

Violin plots of specific characteristics of 117 regulatory elements (REs) containing survivin-ChIP peaks and the remaining 852 RE by length, distance to TSS, GeneHancer score, GeneConnect score, number of TFs per RE.

**Supplementary Table S7**. Positions of immunologically and metabolically relevant GWAS SNPs and associated clinical traits in the vicinity of *PFKFB3, BIRC2* and *SMURF2* genes. according to NHGRI GWAS catalog.

**Supplementary Table S8**. Clinical characteristics RA patients

**Supplementary table S9**. Primers used in the cell experiments

