## Supplementary material for "Chromatin binding of survivin regulates glucose metabolism in the IFN-γ producing CD4^+^ T cells": all supplementary figures and tables

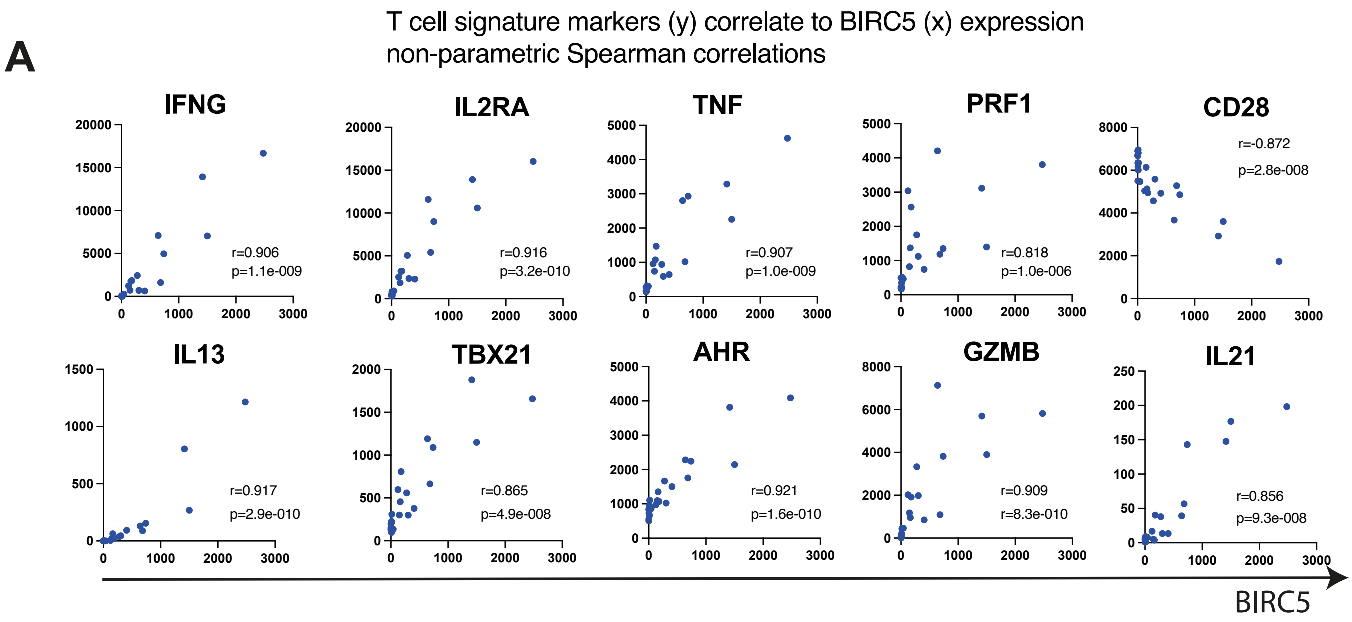


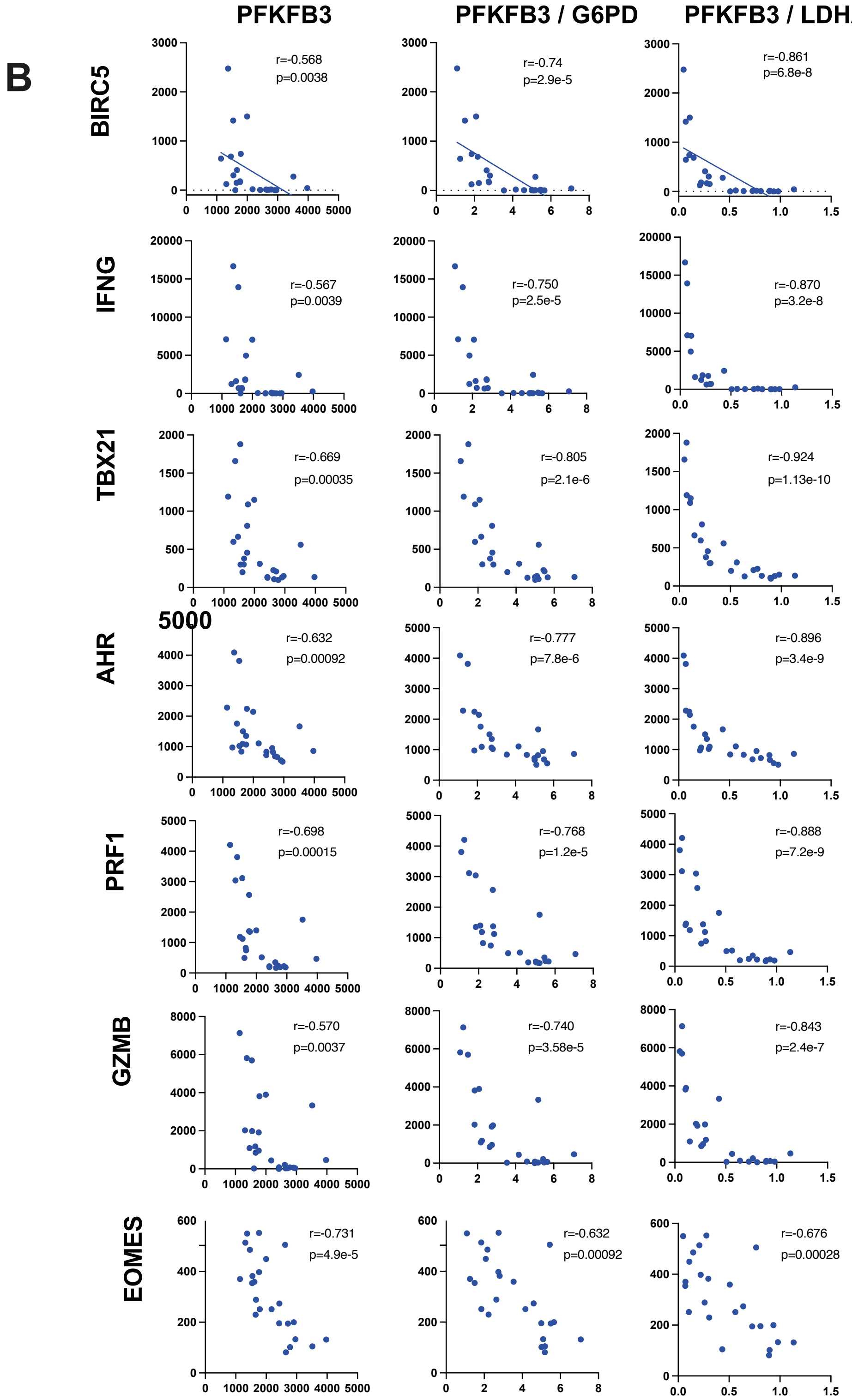


**Supplementary Figure S1**
A. Spearman correlation analysis of expression by normalized RNAseq values between BIRC5 and Th1 signature genes
B. Spearman correlation analysis of expression by normalized RNAseq values between PFKFB3, and its ratio PFKFB3/LDHA and PFKFB3/G6FD with BIRC5 and Th1 signature genes

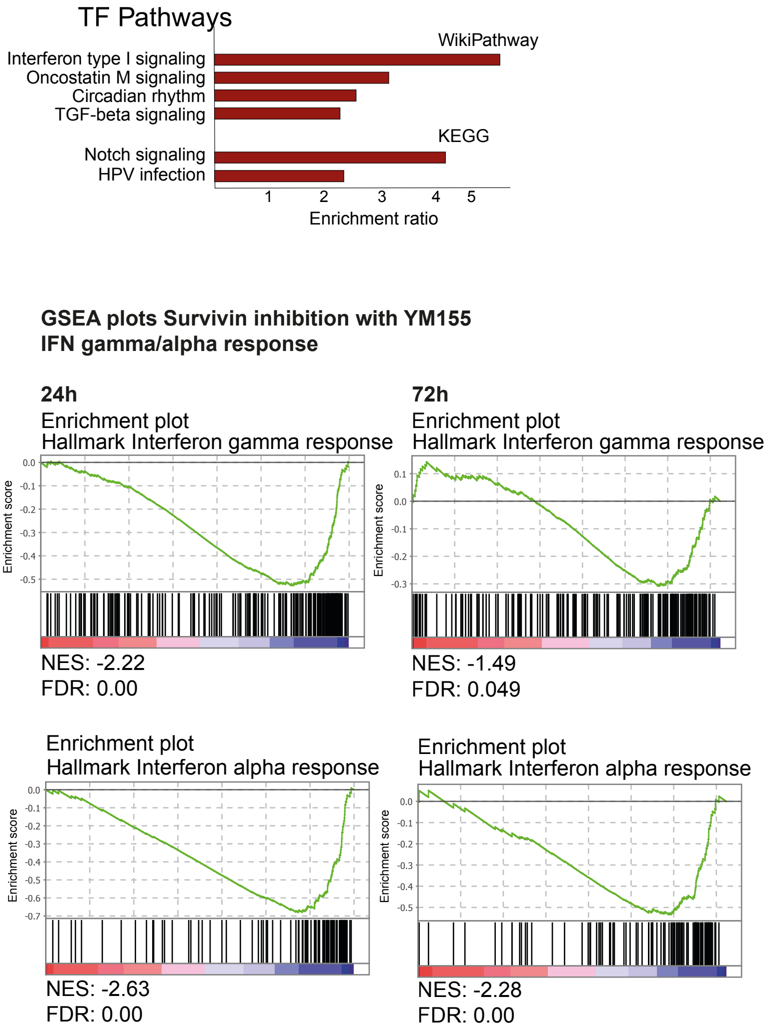


**Supplementary figure S3.**
A. Bar plot of immunologically relevant processes regulated by the proteins colocalized with survivin-ChIP peaks (10% overlap) identified in WikiPathway and KEGG.
B. Barcode plots of the IFN and IFN signalling pathway for the DEG in YM155-treated CD4+ cells, by GSEA analysis

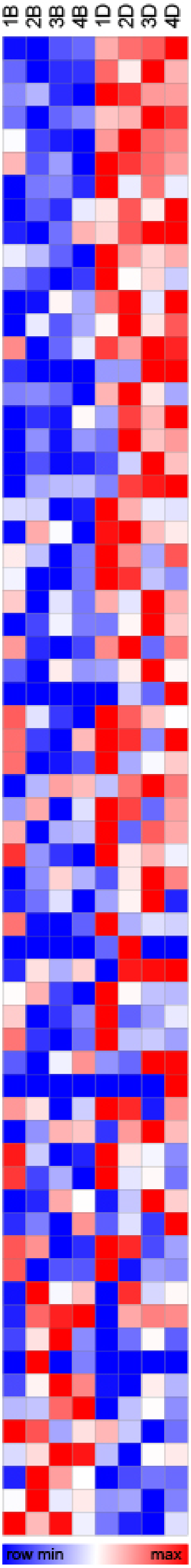


| Gene | pvalue | padj | FC YM IFN/ctr IFN |
|---|---|---|---|
| HNRNPA2B1 | 1,4E-03 | 1,0E-01 | 1,103 |
| CAMSAP2 | 1,2E-03 | 9,2E-02 | 1,281 |
| SMURF2 | 6,7E-12 | 5,1E-08 | 1,290 |
| PMEPA1 | 3,0E-08 | 5,3E-05 | 1,369 |
| FLT3LG | 1,3E-04 | 2,1E-02 | 1,698 |
| PGM2L1 | 6,4E-04 | 5,5E-02 | 1,293 |
| ESCO1 | 1,2E-04 | 2,0E-02 | 1,270 |
| ELK3 | 6,1E-05 | 1,2E-02 | 1,171 |
| ZFP90 | 2,2E-04 | 3,2E-02 | 1,207 |
| PDCD4 | 2,1E-04 | 3,1E-02 | 1,123 |
| KTN1 | 3,1E-04 | 3,9E-02 | 1,167 |
| HNRNPA3 | 4,0E-03 | 2,7E-01 | 1,120 |
| DIPK2A | 2,0E-03 | 1,4E-01 | 1,370 |
| SAMD1 | 5,5E-04 | 4,9E-02 | 1,209 |
| DTL | 2,9E-03 | 2,0E-01 | 12,019 |
| SKIL | 1,1E-03 | 8,6E-02 | 1,216 |
| MIDN | 2,4E-04 | 3,4E-02 | 1,278 |
| SKI | 2,6E-05 | 9,4E-03 | 1,252 |
| QPCT | 2,3E-03 | 1,6E-01 | 1,786 |
| SMAD7 | 2,1E-03 | 1,5E-01 | 1,185 |
| BIRC2 | 5,2E-07 | 5,6E-04 | 1,201 |
| EMB | 7,8E-04 | 6,4E-02 | 1,155 |
| TGFA | 1,0E-04 | 1,7E-02 | 1,390 |
| GJC3 | 4,2E-03 | 2,8E-01 | 3,510 |
| SPSB1 | 3,0E-04 | 3,8E-02 | 1,455 |
| LDLRAD4 | 6,2E-04 | 5,4E-02 | 1,143 |
| NKD1 | 2,4E-03 | 1,7E-01 | 1,280 |
| MYO1C | 3,3E-04 | 4,0E-02 | 1,369 |
| SEZ6L2 | 1,2E-11 | 7,1E-08 | |
| TES | 2,8E-03 | 1,9E-01 | 1,118 |
| ZFP36L1 | 1,5E-03 | 1,1E-01 | 1,131 |
| BPGM | 1,3E-03 | 9,8E-02 | 1,201 |
| PERP | 4,6E-03 | 3,1E-01 | 1,412 |
| RBMS1 | 4,4E-03 | 3,0E-01 | 1,127 |
| SLC25A53 | 1,5E-04 | 2,3E-02 | 1,272 |
| PFKFB3 | 7,1E-06 | 4,1E-03 | 1,208 |
| FAM102B | 8,9E-06 | 4,9E-03 | 1,314 |
| RGS16 | 4,1E-04 | 4,3E-02 | 1,517 |
| SLC20A2 | 1,0E-03 | 7,8E-02 | 1,173 |
| CCDC73 | 4,0E-06 | 2,7E-03 | |
| IL21R | 2,0E-03 | 1,4E-01 | 1,230 |
| STK17A | 4,3E-03 | 2,9E-01 | 1,128 |
| SIAH3 | 5,0E-04 | 4,5E-02 | 1,421 |
| OCIAD2 | 3,5E-03 | 2,4E-01 | 1,144 |
| CERK | 1,8E-03 | 1,3E-01 | 1,117 |
| RPA4 | 2,9E-04 | 3,8E-02 | |
| HVCN1 | 2,4E-04 | 3,3E-02 | 1,288 |
| FOXP4 | 3,9E-03 | 2,7E-01 | 1,359 |
| GRAP | 4,7E-03 | 3,1E-01 | 1,153 |
| FBLN7 | 8,3E-04 | 6,7E-02 | 1,224 |
| SLC1A4 | 4,0E-03 | 2,7E-01 | 1,253 |
| CDK5R1 | 2,2E-11 | 1,1E-07 | 1,432 |
| NET1 | 1,5E-03 | 1,1E-01 | 1,182 |
| CTSL | 1,7E-07 | 2,3E-04 | 1,260 |
| GNLY | 1,5E-03 | 1,1E-01 | 0,887 |
| PRF1 | 7,4E-05 | 1,3E-02 | 0,823 |
| TTYH2 | 1,0E-03 | 7,7E-02 | 0,739 |
| TMEM158 | 1,3E-09 | 3,7E-06 | 0,000 |
| ARID5B | 6,4E-05 | 1,2E-02 | 0,791 |
| SESN2 | 3,3E-03 | 2,3E-01 | 0,728 |
| TMEM25 | 2,0E-03 | 1,4E-01 | 0,790 |
| PYHIN1 | 2,1E-03 | 1,5E-01 | 0,833 |
| CXCL8 | 2,5E-03 | 1,8E-01 | 0,266 |
| UBE2T | 3,2E-04 | 4,0E-02 | 0,546 |
| NLRX1 | 4,1E-03 | 2,8E-01 | 0,763 |

**Supplementary figure S4**
Heatmap of 77 protein-coding DEGs (RNASeq normalized value, basemean >0.5, DESeq2 unadjusted p-value <0.005) in YM155-treated CD4+ cells for 24h, last 2h stimulated with IFN (healthy, n=4).

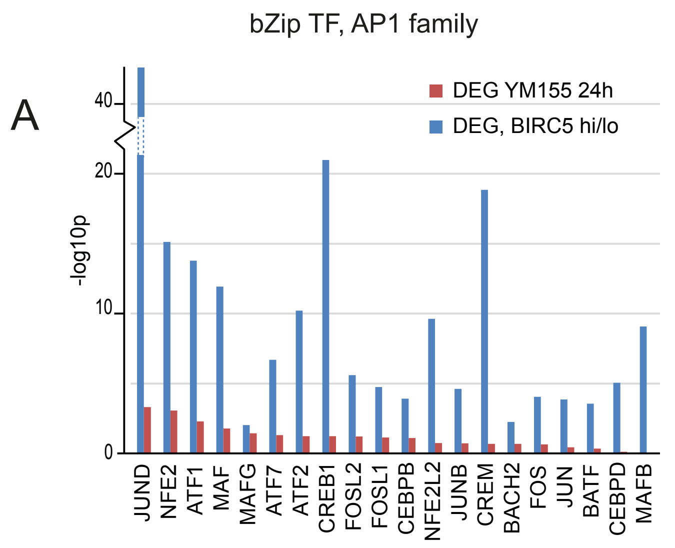


**Supplementary figure 5S**
A. Forest plot of enrichment signifiance of bZIP family proteins in BIRC5hi and BIRC5lo CD4+ cells (blue, RA patients, n=24) and in YM155-treated CD4+ cells (red, healthy females, n=4). P-values are obtained by DESeq2.
B. Box plots of normalized RNAseq values in BIRC5hi and BIRC5lo CD4+ cells (RA patients, n=24) and in YM155-treated CD4+ cells (healthy females, n=4). RNAseq were analysed by DESeq2. C. D. Forest plot of enrichment and p-values of IRF1 and SMAD3 interactors (BioGrid database, >1 physical evidence) in BIRC5hi and BIRC5lo CD4+ cells (red, RA patients, n=24). RNAseq were analysed by DESeq2.

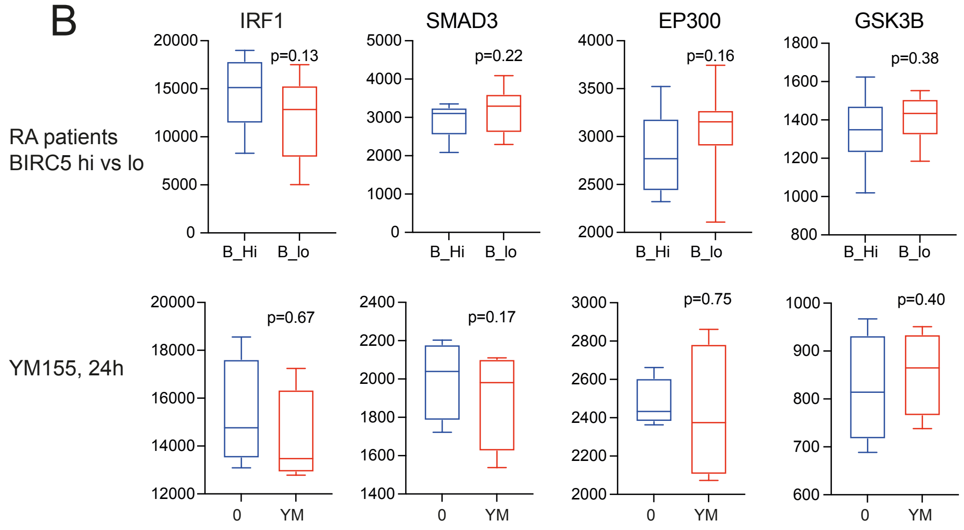


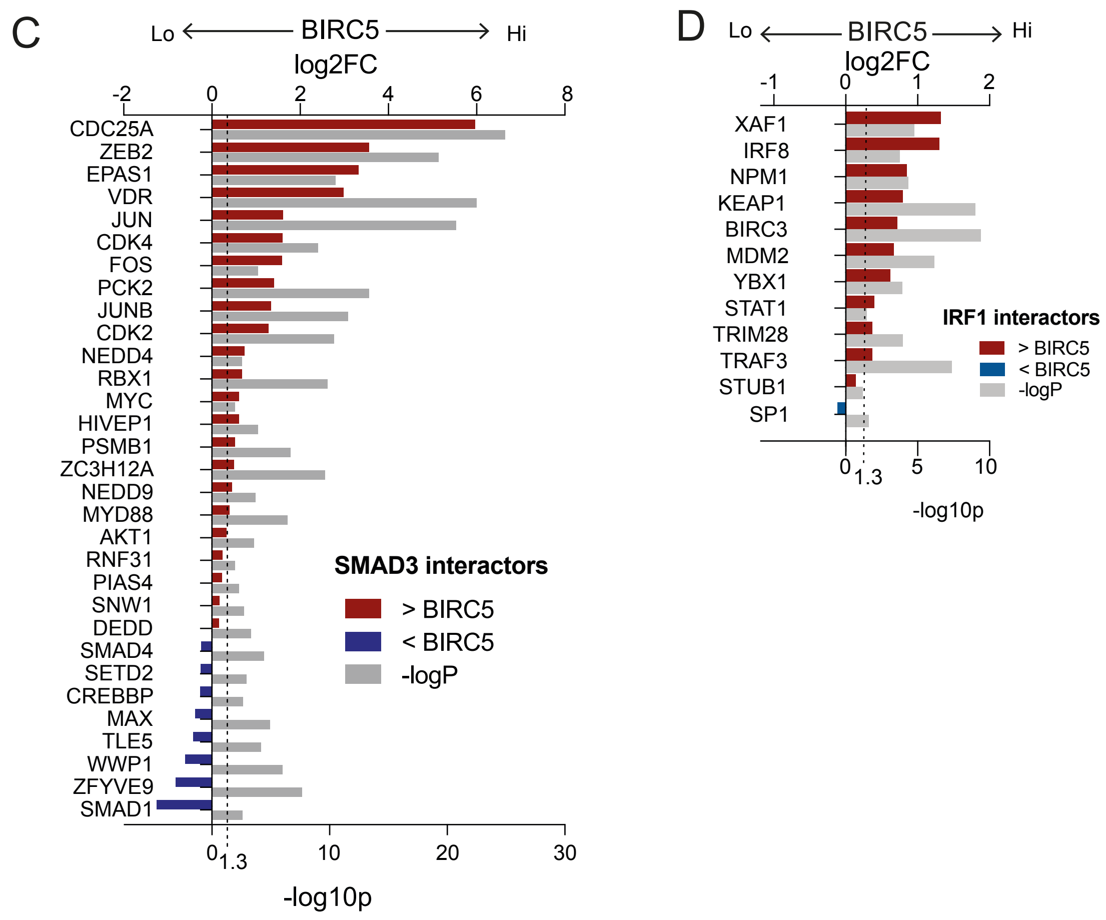


Suppl Figure 5

Descriptives of RE with and without survivin peaks

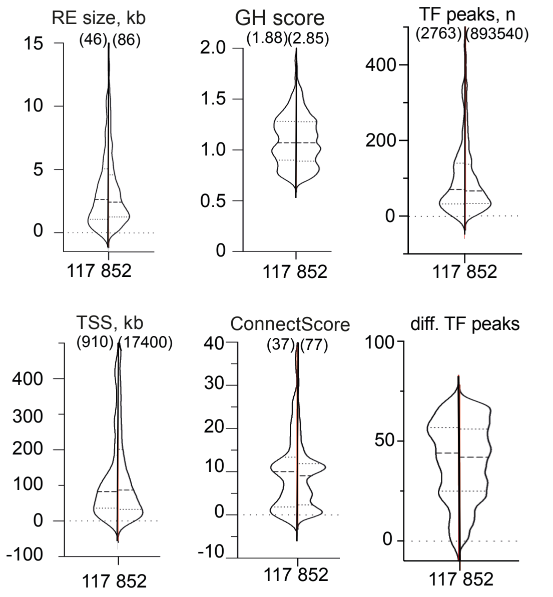


**Supplementary Figure S6**
Violin plots of specific characteristics of 117 regulatory elements (REs) containing survivin-ChIP peaks and the remaining 852 RE by length, distance to TSS, GeneHancer score, GeneConnect score, number of TFs per RE.

| Main//component | Group_ID | term_ID | description | pos_Y | pos_Y | log10 p-value | Functional category supergroup: label |
|---|---|---|---|---|---|---|---|
| Main | 48518_main | GO:0048518 | positive regulation of biological process | -8,426 | 0,293 | -12,2899 | 1 : Regulation of biological processes |
| Main | 48519_main | GO:0048519 | negative regulation of biological process | -8,42 | 0,1 | -8,2312 | 1 : Regulation of biological processes |
| Main | 48523_main | GO:0048523 | negative regulation of cellular process | -8,414 | -0,399 | -8,0696 | 1 : Regulation of biological processes |
| Component | 48523_subgroup | GO:0060968 | regulation of gene silencing | -8,414 | -0,399 | -1,9862 | 1 : Regulation of biological processes |
| Component | 48523_subgroup | GO:0031324 | negative regulation of cellular metabolic process | -8,414 | -0,399 | -6,2707 | 1 : Regulation of biological processes |
| Component | 48523_subgroup | GO:0045814 | negative regulation of gene expression, epigenetic | -8,414 | -0,399 | -2,4339 | 1 : Regulation of biological processes |
| Component | 48523_subgroup | GO:1902679 | negative regulation of RNA biosynthetic process | -8,414 | -0,399 | -6,523 | 1 : Regulation of biological processes |
| Component | 48523_subgroup | GO:0031327 | negative regulation of cellular biosynthetic process | -8,414 | -0,399 | -5,8855 | 1 : Regulation of biological processes |
| Component | 48523_subgroup | GO:0051253 | negative regulation of RNA metabolic process | -8,414 | -0,399 | -6,6744 | 1 : Regulation of biological processes |
| Component | 48523_subgroup | GO:0045934 | negative regulation of nucleobase-containing compound metabolic process | -8,414 | -0,399 | -6,3532 | 1 : Regulation of biological processes |
| Component | 48523_subgroup | GO:0045892 | negative regulation of transcription, DNA-templated | -8,414 | -0,399 | -6,6066 | 1 : Regulation of biological processes |
| Component | 48523_subgroup | GO:0010629 | negative regulation of gene expression | -8,414 | -0,399 | -7,0717 | 1 : Regulation of biological processes |
| Component | 48523_subgroup | GO:0010605 | negative regulation of macromolecule metabolic process | -8,414 | -0,399 | -6,3093 | 1 : Regulation of biological processes |
| Component | 48523_subgroup | GO:0010558 | negative regulation of macromolecule biosynthetic process | -8,414 | -0,399 | -5,9938 | 1 : Regulation of biological processes |
| Component | 48523_subgroup | GO:2000757 | negative regulation of peptidyl-lysine acetylation | -8,414 | -0,399 | -1,836 | 1 : Regulation of biological processes |
| Component | 48523_subgroup | GO:1901984 | negative regulation of protein acetylation | -8,414 | -0,399 | -1,836 | 1 : Regulation of biological processes |
| Component | 48523_subgroup | GO:2000113 | negative regulation of cellular macromolecule biosynthetic process | -8,414 | -0,399 | -6,0772 | 1 : Regulation of biological processes |
| Component | 48523_subgroup | GO:1903507 | negative regulation of nucleic acid-templated transcription | -8,414 | -0,399 | -6,523 | 1 : Regulation of biological processes |
| Component | 48523_subgroup | GO:0009892 | negative regulation of metabolic process | -8,414 | -0,399 | -6,4509 | 1 : Regulation of biological processes |
| Component | 48523_subgroup | GO:0009890 | negative regulation of biosynthetic process | -8,414 | -0,399 | -5,8855 | 1 : Regulation of biological processes |
| Component | 48523_subgroup | GO:0000122 | negative regulation of transcription from RNA polymerase II promoter | -8,414 | -0,399 | -4,6951 | 1 : Regulation of biological processes |
| Component | 48523_subgroup | GO:0016458 | gene silencing | -8,414 | -0,399 | -2,3613 | 1 : Regulation of biological processes |
| Component | 48523_subgroup | GO:0051129 | negative regulation of cellular component organization | -8,414 | -0,399 | -1,7627 | 1 : Regulation of biological processes |
| Component | 48523_subgroup | GO:0051172 | negative regulation of nitrogen compound metabolic process | -8,414 | -0,399 | -5,6451 | 1 : Regulation of biological processes |
| Main | 48522_main | GO:0048522 | positive regulation of cellular process | -8,359 | -0,343 | -12,2899 | 1 : Regulation of biological processes |
| Component | 48522_subgroup | GO:0032270 | positive regulation of cellular protein metabolic process | -8,359 | -0,343 | -3,2813 | 1 : Regulation of biological processes |
| Component | 48522_subgroup | GO:0010720 | positive regulation of cell development | -8,359 | -0,343 | -3,5186 | 1 : Regulation of biological processes |
| Component | 48522_subgroup | GO:1902680 | positive regulation of RNA biosynthetic process | -8,359 | -0,343 | -8,1412 | 1 : Regulation of biological processes |
| Component | 48522_subgroup | GO:0031331 | positive regulation of cellular catabolic process | -8,359 | -0,343 | -1,836 | 1 : Regulation of biological processes |
| Component | 48522_subgroup | GO:0031328 | positive regulation of cellular biosynthetic process | -8,359 | -0,343 | -7,8671 | 1 : Regulation of biological processes |
| Component | 48522_subgroup | GO:0031325 | positive regulation of cellular metabolic process | -8,359 | -0,343 | -9,8296 | 1 : Regulation of biological processes |
| Component | 48522_subgroup | GO:0051247 | positive regulation of protein metabolic process | -8,359 | -0,343 | -3,1871 | 1 : Regulation of biological processes |
| Component | 48522_subgroup | GO:0051240 | positive regulation of multicellular organismal process | -8,359 | -0,343 | -6,6349 | 1 : Regulation of biological processes |
| Component | 48522_subgroup | GO:0051254 | positive regulation of RNA metabolic process | -8,359 | -0,343 | -8,6604 | 1 : Regulation of biological processes |
| Component | 48522_subgroup | GO:0045935 | positive regulation of nucleobase-containing compound metabolic process | -8,359 | -0,343 | -8,5158 | 1 : Regulation of biological processes |
| Component | 48522_subgroup | GO:0010628 | positive regulation of gene expression | -8,359 | -0,343 | -9,2714 | 1 : Regulation of biological processes |
| Component | 48522_subgroup | GO:0045893 | positive regulation of transcription, DNA-templated | -8,359 | -0,343 | -8,9174 | 1 : Regulation of biological processes |
| Component | 48522_subgroup | GO:0010604 | positive regulation of macromolecule metabolic process | -8,359 | -0,343 | -10,0175 | 1 : Regulation of biological processes |
| Component | 48522_subgroup | GO:0045944 | positive regulation of transcription from RNA polymerase II promoter | -8,359 | -0,343 | -8,7674 | 1 : Regulation of biological processes |
| Component | 48522_subgroup | GO:0050769 | positive regulation of neurogenesis | -8,359 | -0,343 | -3,9553 | 1 : Regulation of biological processes |
| Component | 48522_subgroup | GO:0050767 | regulation of neurogenesis | -8,359 | -0,343 | -2,6495 | 1 : Regulation of biological processes |
| Component | 48522_subgroup | GO:0010557 | positive regulation of macromolecule biosynthetic process | -8,359 | -0,343 | -8,3543 | 1 : Regulation of biological processes |
| Component | 48522_subgroup | GO:0031401 | positive regulation of protein modification process | -8,359 | -0,343 | -2,6883 | 1 : Regulation of biological processes |
| Component | 48522_subgroup | GO:2000026 | regulation of multicellular organismal development | -8,359 | -0,343 | -6,3046 | 1 : Regulation of biological processes |
| Component | 48522_subgroup | GO:0045664 | regulation of neuron differentiation | -8,359 | -0,343 | -2,6011 | 1 : Regulation of biological processes |
| Component | 48522_subgroup | GO:0045666 | positive regulation of neuron differentiation | -8,359 | -0,343 | -2,7345 | 1 : Regulation of biological processes |
| Component | 48522_subgroup | GO:0051962 | positive regulation of nervous system development | -8,359 | -0,343 | -3,8439 | 1 : Regulation of biological processes |
| Component | 48522_subgroup | GO:0045597 | positive regulation of cell differentiation | -8,359 | -0,343 | -4,4708 | 1 : Regulation of biological processes |
| Component | 48522_subgroup | GO:0001525 | angiogenesis | -8,359 | -0,343 | -1,9899 | 1 : Regulation of biological processes |
| Component | 48522_subgroup | GO:1903508 | positive regulation of nucleic acid-templated transcription | -8,359 | -0,343 | -8,1412 | 1 : Regulation of biological processes |
| Component | 48522_subgroup | GO:0009893 | positive regulation of metabolic process | -8,359 | -0,343 | -9,8699 | 1 : Regulation of biological processes |
| Component | 48522_subgroup | GO:0060284 | regulation of cell development | -8,359 | -0,343 | -2,6243 | 1 : Regulation of biological processes |
| Component | 48522_subgroup | GO:0009891 | positive regulation of biosynthetic process | -8,359 | -0,343 | -8,1858 | 1 : Regulation of biological processes |
| Component | 48522_subgroup | GO:1904018 | positive regulation of vasculature development | -8,359 | -0,343 | -2,5891 | 1 : Regulation of biological processes |
| Component | 48522_subgroup | GO:0051094 | positive regulation of developmental process | -8,359 | -0,343 | -4,858 | 1 : Regulation of biological processes |
| Component | 48522_subgroup | GO:0051130 | positive regulation of cellular component organization | -8,359 | -0,343 | -4,1219 | 1 : Regulation of biological processes |
| Component | 48522_subgroup | GO:0051173 | positive regulation of nitrogen compound metabolic process | -8,359 | -0,343 | -9,8296 | 1 : Regulation of biological processes |
| Main | 51239_main | GO:0051239 | regulation of multicellular organismal process | -8,077 | 0,425 | -7,9068 | 1 : Regulation of biological processes |
| Component | 51239_subgroup | GO:0003008 | system process | -8,077 | 0,425 | -1,877 | 1 : Regulation of biological processes |
| Component | 51239_subgroup | GO:0010634 | positive regulation of epithelial cell migration | -8,077 | 0,425 | -1,836 | 1 : Regulation of biological processes |
| Component | 51239_subgroup | GO:0010631 | epithelial cell migration | -8,077 | 0,425 | -3,5954 | 1 : Regulation of biological processes |
| Component | 51239_subgroup | GO:0010632 | regulation of epithelial cell migration | -8,077 | 0,425 | -2,288 | 1 : Regulation of biological processes |
| Component | 51239_subgroup | GO:0071706 | tumor necrosis factor superfamily cytokine production | -8,077 | 0,425 | -2,1335 | 1 : Regulation of biological processes |
| Component | 51239_subgroup | GO:0010594 | regulation of endothelial cell migration | -8,077 | 0,425 | -2,3193 | 1 : Regulation of biological processes |
| Component | 51239_subgroup | GO:0010595 | positive regulation of endothelial cell migration | -8,077 | 0,425 | -1,8937 | 1 : Regulation of biological processes |
| Component | 51239_subgroup | GO:0001816 | cytokine production | -8,077 | 0,425 | -4,3355 | 1 : Regulation of biological processes |
| Component | 51239_subgroup | GO:0001819 | positive regulation of cytokine production | -8,077 | 0,425 | -2,2702 | 1 : Regulation of biological processes |
| Component | 51239_subgroup | GO:0001817 | regulation of cytokine production | -8,077 | 0,425 | -4,4267 | 1 : Regulation of biological processes |
| Component | 51239_subgroup | GO:0043542 | endothelial cell migration | -8,077 | 0,425 | -3,1508 | 1 : Regulation of biological processes |
| Component | 51239_subgroup | GO:0043534 | blood vessel endothelial cell migration | -8,077 | 0,425 | -3,0058 | 1 : Regulation of biological processes |
| Component | 51239_subgroup | GO:0043536 | positive regulation of blood vessel endothelial cell migration | -8,077 | 0,425 | -2,1335 | 1 : Regulation of biological processes |
| Component | 51239_subgroup | GO:0043535 | regulation of blood vessel endothelial cell migration | -8,077 | 0,425 | -2,6904 | 1 : Regulation of biological processes |
| Component | 51239_subgroup | GO:0014897 | striated muscle hypertrophy | -8,077 | 0,425 | -1,836 | 1 : Regulation of biological processes |
| Component | 51239_subgroup | GO:0014896 | muscle hypertrophy | -8,077 | 0,425 | -1,836 | 1 : Regulation of biological processes |
| Component | 51239_subgroup | GO:0003300 | cardiac muscle hypertrophy | -8,077 | 0,425 | -1,836 | 1 : Regulation of biological processes |
| Component | 51239_subgroup | GO:0032602 | chemokine production | -8,077 | 0,425 | -1,836 | 1 : Regulation of biological processes |
| Component | 51239_subgroup | GO:0002042 | cell migration involved in sprouting angiogenesis | -8,077 | 0,425 | -1,8576 | 1 : Regulation of biological processes |
| Component | 51239_subgroup | GO:0032640 | tumor necrosis factor production | -8,077 | 0,425 | -2,1335 | 1 : Regulation of biological processes |
| Component | 51239_subgroup | GO:0032680 | regulation of tumor necrosis factor production | -8,077 | 0,425 | -2,1335 | 1 : Regulation of biological processes |
| Component | 51239_subgroup | GO:1903555 | regulation of tumor necrosis factor superfamily cytokine production | -8,077 | 0,425 | -2,1335 | 1 : Regulation of biological processes |
| Component | 51239_subgroup | GO:0090130 | tissue migration | -8,077 | 0,425 | -3,2176 | 1 : Regulation of biological processes |
| Component | 51239_subgroup | GO:0090132 | epithelium migration | -8,077 | 0,425 | -3,4144 | 1 : Regulation of biological processes |
| Component | 51239_subgroup | GO:0001667 | ameboidal-type cell migration | -8,077 | 0,425 | -2,9428 | 1 : Regulation of biological processes |
| Main | 44093_main | GO:0044093 | positive regulation of molecular function | -7,778 | -0,291 | -2,4973 | 1 : Regulation of biological processes |
| Component | 44093_subgroup | GO:0051099 | positive regulation of binding | -7,778 | -0,291 | -2,297 | 1 : Regulation of biological processes |
| Main | 9605_main | GO:0009605 | response to external stimulus | -2,708 | -7,315 | -2,8203 | 2 : Response to organic substances |
| Main | 42221_main | GO:0042221 | response to chemical | -2,706 | -7,2 | -8,7674 | 2 : Response to organic substances |
| Component | 42221_subgroup | GO:0051716 | cellular response to stimulus | -2,706 | -7,2 | -9,2514 | 2 : Response to organic substances |
| Component | 42221_subgroup | GO:0071214 | cellular response to abiotic stimulus | -2,706 | -7,2 | -2,3551 | 2 : Response to organic substances |
| Component | 42221_subgroup | GO:0009611 | response to wounding | -2,706 | -7,2 | -2,2497 | 2 : Response to organic substances |
| Component | 42221_subgroup | GO:0070482 | response to oxygen levels | -2,706 | -7,2 | -2,8622 | 2 : Response to organic substances |
| Component | 42221_subgroup | GO:0036293 | response to decreased oxygen levels | -2,706 | -7,2 | -2,9428 | 2 : Response to organic substances |
| Component | 42221_subgroup | GO:0036294 | cellular response to decreased oxygen levels | -2,706 | -7,2 | -1,848 | 2 : Response to organic substances |
| Component | 42221_subgroup | GO:0033554 | cellular response to stress | -2,706 | -7,2 | -4,9046 | 2 : Response to organic substances |
| Component | 42221_subgroup | GO:0009314 | response to radiation | -2,706 | -7,2 | -2,4172 | 2 : Response to organic substances |
| Component | 42221_subgroup | GO:0071456 | cellular response to hypoxia | -2,706 | -7,2 | -1,9862 | 2 : Response to organic substances |
| Component | 42221_subgroup | GO:0006979 | response to oxidative stress | -2,706 | -7,2 | -3,5973 | 2 : Response to organic substances |
| Component | 42221_subgroup | GO:0009416 | response to light stimulus | -2,706 | -7,2 | -2,7543 | 2 : Response to organic substances |
| Component | 42221_subgroup | GO:0006952 | defense response | -2,706 | -7,2 | -3,7096 | 2 : Response to organic substances |
| Component | 42221_subgroup | GO:0006950 | response to stress | -2,706 | -7,2 | -7,9135 | 2 : Response to organic substances |
| Component | 42221_subgroup | GO:0006954 | inflammatory response | -2,706 | -7,2 | -1,8954 | 2 : Response to organic substances |
| Component | 42221_subgroup | GO:0071482 | cellular response to light stimulus | -2,706 | -7,2 | -2,2122 | 2 : Response to organic substances |
| Component | 42221_subgroup | GO:0071478 | cellular response to radiation | -2,706 | -7,2 | -2,8958 | 2 : Response to organic substances |
| Component | 42221_subgroup | GO:0001666 | response to hypoxia | -2,706 | -7,2 | -3,0701 | 2 : Response to organic substances |
| Main | 9628_main | GO:0009628 | response to abiotic stimulus | -2,586 | -7,443 | -4,5011 | 2 : Response to organic substances |
| Main | 9719_main | GO:0009719 | response to endogenous stimulus | -2,468 | -7,192 | -5,2489 | 2 : Response to organic substances |
| Main | 10033_main | GO:0010033 | response to organic substance | -2,174 | -7,269 | -9,3126 | 2 : Response to organic substances |
| Component | 10033_subgroup | GO:1901701 | cellular response to oxygen-containing compound | -2,174 | -7,269 | -5,7969 | 2 : Response to organic substances |
| Component | 10033_subgroup | GO:0042542 | response to hydrogen peroxide | -2,174 | -7,269 | -2,7407 | 2 : Response to organic substances |
| Component | 10033_subgroup | GO:1901698 | response to nitrogen compound | -2,174 | -7,269 | -2,5201 | 2 : Response to organic substances |
| Component | 10033_subgroup | GO:1901699 | cellular response to nitrogen compound | -2,174 | -7,269 | -2,4172 | 2 : Response to organic substances |
| Component | 10033_subgroup | GO:1901700 | response to oxygen-containing compound | -2,174 | -7,269 | -6,0772 | 2 : Response to organic substances |
| Component | 10033_subgroup | GO:0000302 | response to reactive oxygen species | -2,174 | -7,269 | -2,2885 | 2 : Response to organic substances |
| Component | 10033_subgroup | GO:0048545 | response to steroid hormone | -2,174 | -7,269 | -2,422 | 2 : Response to organic substances |
| Component | 10033_subgroup | GO:0042493 | response to drug | -2,174 | -7,269 | -5,2444 | 2 : Response to organic substances |
| Component | 10033_subgroup | GO:0032870 | cellular response to hormone stimulus | -2,174 | -7,269 | -3,5562 | 2 : Response to organic substances |
| Component | 10033_subgroup | GO:0019221 | cytokine-mediated signaling pathway | -2,174 | -7,269 | -2,2497 | 2 : Response to organic substances |
| Component | 10033_subgroup | GO:0009636 | response to toxic substance | -2,174 | -7,269 | -3,4297 | 2 : Response to organic substances |
| Component | 10033_subgroup | GO:0033993 | response to lipid | -2,174 | -7,269 | -5,4266 | 2 : Response to organic substances |
| Component | 10033_subgroup | GO:0071363 | cellular response to growth factor stimulus | -2,174 | -7,269 | -1,8176 | 2 : Response to organic substances |
| Component | 10033_subgroup | GO:0070887 | cellular response to chemical stimulus | -2,174 | -7,269 | -8,9465 | 2 : Response to organic substances |
| Component | 10033_subgroup | GO:0071383 | cellular response to steroid hormone stimulus | -2,174 | -7,269 | -2,3551 | 2 : Response to organic substances |
| Component | 10033_subgroup | GO:0070301 | cellular response to hydrogen peroxide | -2,174 | -7,269 | -2,8226 | 2 : Response to organic substances |
| Component | 10033_subgroup | GO:0071396 | cellular response to lipid | -2,174 | -7,269 | -3,5928 | 2 : Response to organic substances |
| Component | 10033_subgroup | GO:0071407 | cellular response to organic cyclic compound | -2,174 | -7,269 | -3,7727 | 2 : Response to organic substances |
| Component | 10033_subgroup | GO:1901652 | response to peptide | -2,174 | -7,269 | -2,6059 | 2 : Response to organic substances |
| Component | 10033_subgroup | GO:0071310 | cellular response to organic substance | -2,174 | -7,269 | -8,617 | 2 : Response to organic substances |
| Component | 10033_subgroup | GO:0010035 | response to inorganic substance | -2,174 | -7,269 | -2,4295 | 2 : Response to organic substances |
| Component | 10033_subgroup | GO:0071345 | cellular response to cytokine stimulus | -2,174 | -7,269 | -2,9712 | 2 : Response to organic substances |
| Component | 10033_subgroup | GO:0009755 | hormone-mediated signaling pathway | -2,174 | -7,269 | -1,8618 | 2 : Response to organic substances |
| Component | 10033_subgroup | GO:0009749 | response to glucose | -2,174 | -7,269 | -2,0168 | 2 : Response to organic substances |
| Component | 10033_subgroup | GO:0009746 | response to hexose | -2,174 | -7,269 | -2,0168 | 2 : Response to organic substances |
| Component | 10033_subgroup | GO:0009743 | response to carbohydrate | -2,174 | -7,269 | -1,894 | 2 : Response to organic substances |
| Component | 10033_subgroup | GO:0009725 | response to hormone | -2,174 | -7,269 | -4,9424 | 2 : Response to organic substances |
| Component | 10033_subgroup | GO:0060759 | regulation of response to cytokine stimulus | -2,174 | -7,269 | -1,836 | 2 : Response to organic substances |

| | | | | | | | |
|---|---|---|---|---|---|---|---|
| Component | 10033_subgroup | GO:0034097 | response to cytokine | -2,174 | -7,269 | -2,9712 | 2 : Response to organic substances |
| Component | 10033_subgroup | GO:0043401 | steroid hormone mediated signaling pathway | -2,174 | -7,269 | -1,8618 | 2 : Response to organic substances |
| Component | 10033_subgroup | GO:0046677 | response to antibiotic | -2,174 | -7,269 | -3,8466 | 2 : Response to organic substances |
| Component | 10033_subgroup | GO:0034284 | response to monosaccharide | -2,174 | -7,269 | -2,0168 | 2 : Response to organic substances |
| Component | 10033_subgroup | GO:0071417 | cellular response to organonitrogen compound | -2,174 | -7,269 | -2,3309 | 2 : Response to organic substances |
| Component | 10033_subgroup | GO:0034599 | cellular response to oxidative stress | -2,174 | -7,269 | -3,1081 | 2 : Response to organic substances |
| Component | 10033_subgroup | GO:0010243 | response to organonitrogen compound | -2,174 | -7,269 | -2,4882 | 2 : Response to organic substances |
| Component | 10033_subgroup | GO:0034614 | cellular response to reactive oxygen species | -2,174 | -7,269 | -2,1335 | 2 : Response to organic substances |
| Component | 10033_subgroup | GO:0014070 | response to organic cyclic compound | -2,174 | -7,269 | -4,49 | 2 : Response to organic substances |
| Component | 10033_subgroup | GO:0071495 | cellular response to endogenous stimulus | -2,174 | -7,269 | -4,6232 | 2 : Response to organic substances |
| Main | 35690_main | GO:0035690 | cellular response to drug | -1,874 | -7,168 | -4,9046 | 2 : Response to organic substances |
| Main | 97237_main | GO:0097237 | cellular response to toxic substance | -1,492 | -7,147 | -3,274 | 2 : Response to organic substances |
| Main | 71236_main | GO:0071236 | cellular response to antibiotic | -1,142 | -7,411 | -4,3956 | 2 : Response to organic substances |
| Main | 32940_main | GO:0032940 | secretion by cell | -0,74 | 6,239 | -3,5928 | 3: Cellular trafficking |
| Component | 32940_subgroup | GO:0023061 | signal release | -0,74 | 6,239 | -2,6338 | 3: Cellular trafficking |
| Component | 32940_subgroup | GO:0046903 | secretion | -0,74 | 6,239 | -3,9553 | 3: Cellular trafficking |
| Component | 32940_subgroup | GO:0046879 | hormone secretion | -0,74 | 6,239 | -2,3551 | 3: Cellular trafficking |
| Component | 32940_subgroup | GO:0046883 | regulation of hormone secretion | -0,74 | 6,239 | -3,0466 | 3: Cellular trafficking |
| Component | 32940_subgroup | GO:0051046 | regulation of secretion | -0,74 | 6,239 | -3,2949 | 3: Cellular trafficking |
| Component | 32940_subgroup | GO:1903530 | regulation of secretion by cell | -0,74 | 6,239 | -3,4297 | 3: Cellular trafficking |
| Main | 71705_main | GO:0071705 | nitrogen compound transport | -0,731 | 7,259 | -2,956 | 3: Cellular trafficking |
| Main | 8104_main | GO:0008104 | protein localization | -0,542 | 7,062 | -2,8793 | 3: Cellular trafficking |
| Component | 8104_subgroup | GO:0015031 | protein transport | -0,542 | 7,062 | -2,2108 | 3: Cellular trafficking |
| Component | 8104_subgroup | GO:0045184 | establishment of protein localization | -0,542 | 7,062 | -2,1335 | 3: Cellular trafficking |
| Main | 15833_main | GO:0015833 | peptide transport | -0,351 | 6,812 | -2,4384 | 3: Cellular trafficking |
| Component | 15833_subgroup | GO:0090087 | regulation of peptide transport | -0,351 | 6,812 | -1,8287 | 3: Cellular trafficking |
| Component | 15833_subgroup | GO:0042886 | amide transport | -0,351 | 6,812 | -2,4384 | 3: Cellular trafficking |
| Main | 46907_main | GO:0046907 | intracellular transport | -0,293 | 7,084 | -2,0774 | 3: Cellular trafficking |
| Component | 46907_subgroup | GO:0006913 | nucleocytoplasmic transport | -0,293 | 7,084 | -1,9549 | 3: Cellular trafficking |
| Component | 46907_subgroup | GO:0051170 | nuclear import | -0,293 | 7,084 | -1,9763 | 3: Cellular trafficking |
| Component | 46907_subgroup | GO:0051169 | nuclear transport | -0,293 | 7,084 | -1,848 | 3: Cellular trafficking |
| Main | 7049_main | GO:0007049 | cell cycle | -0,456 | 4,711 | -3,7632 | 4: Cell cycle |
| Main | 8219_main | GO:0008219 | cell death | 0,044 | 4,331 | -6,7963 | 4: Cell cycle |
| Main | 6915_main | GO:0006915 | apoptotic process | 0,144 | 3,698 | -6,6066 | 4: Cell cycle |
| Component | 6915_subgroup | GO:0060548 | negative regulation of cell death | 0,144 | 3,698 | -2,7244 | 4: Cell cycle |
| Component | 6915_subgroup | GO:0043069 | negative regulation of programmed cell death | 0,144 | 3,698 | -2,4553 | 4: Cell cycle |
| Component | 6915_subgroup | GO:0043066 | negative regulation of apoptotic process | 0,144 | 3,698 | -2,5218 | 4: Cell cycle |
| Component | 6915_subgroup | GO:0043067 | regulation of programmed cell death | 0,144 | 3,698 | -3,7732 | 4: Cell cycle |
| Component | 6915_subgroup | GO:0012501 | programmed cell death | 0,144 | 3,698 | -6,9031 | 4: Cell cycle |
| Component | 6915_subgroup | GO:0010941 | regulation of cell death | 0,144 | 3,698 | -3,7071 | 4: Cell cycle |
| Component | 6915_subgroup | GO:0042981 | regulation of apoptotic process | 0,144 | 3,698 | -3,5297 | 4: Cell cycle |
| Main | 6476_main | GO:0006476 | protein deacetylation | 3,403 | 3,134 | -2,6876 | 5: Protein modification |
| Component | 6476_subgroup | GO:0016575 | histone deacetylation | 3,403 | 3,134 | -2,4066 | 5: Protein modification |
| Component | 6476_subgroup | GO:0035601 | protein deacylation | 3,403 | 3,134 | -2,6876 | 5: Protein modification |
| Main | 98732_main | GO:0098732 | macromolecule deacylation | 3,555 | 3,103 | -2,6876 | 5: Protein modification |
| Main | 46777_main | GO:0046777 | protein autophosphorylation | 3,756 | 2,609 | -1,836 | 5: Protein modification |
| Main | 43412_main | GO:0043412 | macromolecule modification | 3,95 | 1,316 | -8,6516 | 5: Protein modification |
| Main | 43543_main | GO:0043543 | protein acylation | 4,181 | 3,423 | -1,7627 | 5: Protein modification |
| Main | 18205_main | GO:0018205 | peptidyl-lysine modification | 4,555 | 2,931 | -3,8543 | 5: Protein modification |
| Main | 19538_main | GO:0019538 | protein metabolic process | 4,651 | 1,476 | -8,2312 | 5: Protein modification |
| Main | 44267_main | GO:0044267 | cellular protein metabolic process | 4,666 | 1,483 | -8,4132 | 5: Protein modification |
| Main | 1901564_main | GO:1901564 | organonitrogen compound metabolic process | 4,499 | -2,552 | -8,5134 | 6: Metabolism |
| Main | 16310_main | GO:0016310 | phosphorylation | 4,716 | -0,754 | -3,7046 | 6: Metabolism |
| Component | 16310_subgroup | GO:0006796 | phosphate-containing compound metabolic process | 4,716 | -0,754 | -3,2155 | 6: Metabolism |
| Main | 6259_main | GO:0006259 | DNA metabolic process | 4,717 | -0,083 | -1,9019 | 6: Metabolism |
| Main | 6793_main | GO:0006793 | phosphorus metabolic process | 5,035 | -2,309 | -3,2155 | 6: Metabolism |
| Main | 23051_main | GO:0023051 | regulation of signaling | -8,071 | -1,499 | -6,2561 | |
| Component | 23051_subgroup | GO:0023056 | positive regulation of signaling | -8,071 | -1,499 | -2,422 | |
| Component | 23051_subgroup | GO:0023057 | negative regulation of signaling | -8,071 | -1,499 | -2,5894 | |
| Component | 23051_subgroup | GO:0031347 | regulation of defense response | -8,071 | -1,499 | -2,6059 | |
| Component | 23051_subgroup | GO:0048584 | positive regulation of response to stimulus | -8,071 | -1,499 | -3,7213 | |
| Component | 23051_subgroup | GO:0035556 | intracellular signal transduction | -8,071 | -1,499 | -2,6598 | |
| Component | 23051_subgroup | GO:0048585 | negative regulation of response to stimulus | -8,071 | -1,499 | -2,6186 | |
| Component | 23051_subgroup | GO:0010648 | negative regulation of cell communication | -8,071 | -1,499 | -2,5894 | |
| Component | 23051_subgroup | GO:0010647 | positive regulation of cell communication | -8,071 | -1,499 | -2,422 | |
| Component | 23051_subgroup | GO:0007165 | signal transduction | -8,071 | -1,499 | -6,4163 | |
| Component | 23051_subgroup | GO:0008593 | regulation of Notch signaling pathway | -8,071 | -1,499 | -1,836 | |
| Component | 23051_subgroup | GO:0030518 | intracellular steroid hormone receptor signaling pathway | -8,071 | -1,499 | -2,1588 | |
| Component | 23051_subgroup | GO:0030521 | androgen receptor signaling pathway | -8,071 | -1,499 | -2,3097 | |
| Component | 23051_subgroup | GO:0060765 | regulation of androgen receptor signaling pathway | -8,071 | -1,499 | -3,0147 | |
| Component | 23051_subgroup | GO:0032101 | regulation of response to external stimulus | -8,071 | -1,499 | -2,0795 | |
| Component | 23051_subgroup | GO:1902531 | regulation of intracellular signal transduction | -8,071 | -1,499 | -2,5671 | |
| Component | 23051_subgroup | GO:0033143 | regulation of intracellular steroid hormone receptor signaling pathway | -8,071 | -1,499 | -2,733 | |
| Component | 23051_subgroup | GO:0009968 | negative regulation of signal transduction | -8,071 | -1,499 | -2,4778 | |
| Component | 23051_subgroup | GO:0009966 | regulation of signal transduction | -8,071 | -1,499 | -4,9046 | |
| Component | 23051_subgroup | GO:0080134 | regulation of response to stress | -8,071 | -1,499 | -2,3657 | |
| Main | 42127_main | GO:0042127 | regulation of cell proliferation | -7,961 | -1,023 | -4,4982 | |
| Component | 42127_subgroup | GO:0050679 | positive regulation of epithelial cell proliferation | -7,961 | -1,023 | -2,733 | |
| Component | 42127_subgroup | GO:0008284 | positive regulation of cell proliferation | -7,961 | -1,023 | -3,2194 | |
| Main | 50878_main | GO:0050878 | regulation of body fluid levels | -7,925 | 1,844 | -1,9964 | |
| Component | 44093_subgroup | GO:0051098 | regulation of binding | -7,778 | -0,291 | -1,8176 | |
| Main | 65008_main | GO:0065008 | regulation of biological quality | -7,76 | 0,156 | -7,9803 | |
| Main | 10646_main | GO:0010646 | regulation of cell communication | -7,707 | -1,492 | -6,2561 | |
| Main | 65009_main | GO:0065009 | regulation of molecular function | -7,667 | 0,018 | -3,7632 | |
| Main | 10817_main | GO:0010817 | regulation of hormone levels | -7,397 | 1,158 | -3,5973 | |
| Component | 10817_subgroup | GO:0042592 | homeostatic process | -7,397 | 1,158 | -3,0562 | |
| Main | 48872_main | GO:0048872 | homeostasis of number of cells | -7,375 | 1,331 | -2,0774 | |
| Main | 43900_main | GO:0043900 | regulation of multi-organism process | -7,254 | -1,008 | -3,8443 | |
| Component | 43900_subgroup | GO:0052312 | modulation of transcription in other organism involved in symbiotic interaction | -7,254 | -1,008 | -2,007 | |
| Component | 43900_subgroup | GO:0016032 | viral process | -7,254 | -1,008 | -3,4297 | |
| Component | 43900_subgroup | GO:0050792 | regulation of viral process | -7,254 | -1,008 | -3,3814 | |
| Component | 43900_subgroup | GO:0043921 | modulation by host of viral transcription | -7,254 | -1,008 | -2,007 | |
| Component | 43900_subgroup | GO:0043903 | regulation of symbiosis, encompassing mutualism through parasitism | -7,254 | -1,008 | -3,3814 | |
| Component | 43900_subgroup | GO:0044419 | interspecies interaction between organisms | -7,254 | -1,008 | -3,4297 | |
| Component | 43900_subgroup | GO:0052472 | modulation by host of symbiont transcription | -7,254 | -1,008 | -2,007 | |
| Component | 43900_subgroup | GO:0019080 | viral gene expression | -7,254 | -1,008 | -2,1335 | |
| Component | 43900_subgroup | GO:0019083 | viral transcription | -7,254 | -1,008 | -2,4384 | |
| Component | 43900_subgroup | GO:0044403 | symbiosis, encompassing mutualism through parasitism | -7,254 | -1,008 | -3,4297 | |
| Component | 43900_subgroup | GO:0046782 | regulation of viral transcription | -7,254 | -1,008 | -3,0562 | |
| Main | 51726_main | GO:0051726 | regulation of cell cycle | -6,951 | 1,984 | -3,0527 | |
| Component | 51726_subgroup | GO:0010564 | regulation of cell cycle process | -6,951 | 1,984 | -2,1049 | |
| Component | 51726_subgroup | GO:0022402 | cell cycle process | -6,951 | 1,984 | -2,0986 | |
| Component | 51726_subgroup | GO:0045786 | negative regulation of cell cycle | -6,951 | 1,984 | -1,8218 | |
| Main | 33044_main | GO:0033044 | regulation of chromosome organization | -6,771 | -0,352 | -2,7058 | |
| Component | 33044_subgroup | GO:2001252 | positive regulation of chromosome organization | -6,771 | -0,352 | -2,8083 | |
| Component | 33044_subgroup | GO:0010638 | positive regulation of organelle organization | -6,771 | -0,352 | -2,5898 | |
| Component | 33044_subgroup | GO:0033043 | regulation of organelle organization | -6,771 | -0,352 | -2,3811 | |
| Main | 48583_main | GO:0048583 | regulation of response to stimulus | -6,115 | -3,226 | -5,4266 | |
| Main | 30522_main | GO:0030522 | intracellular receptor signaling pathway | -6,075 | -3,817 | -3,781 | |
| Main | 32879_main | GO:0032879 | regulation of localization | -6,024 | 3,124 | -2,9569 | |
| Main | 40029_main | GO:0040029 | regulation of gene expression, epigenetic | -6,005 | -0,411 | -3,4297 | |
| Main | 7259_main | GO:0007259 | JAK-STAT cascade | -5,888 | -4,262 | -3,0058 | |
| Component | 7259_subgroup | GO:0046425 | regulation of JAK-STAT cascade | -5,888 | -4,262 | -1,836 | |
| Component | 7259_subgroup | GO:1904892 | regulation of STAT cascade | -5,888 | -4,262 | -1,836 | |
| Main | 51246_main | GO:0051246 | regulation of protein metabolic process | -5,873 | 0,98 | -4,0064 | |
| Main | 97696_main | GO:0097696 | STAT cascade | -5,87 | -4,03 | -3,0058 | |
| Main | 51049_main | GO:0051049 | regulation of transport | -5,81 | 3,247 | -2,0987 | |
| Main | 1901532_main | GO:1901532 | regulation of hematopoietic progenitor cell differentiation | -5,779 | 1,807 | -2,1335 | |
| Main | 9914_main | GO:0009914 | hormone transport | -5,702 | 3,777 | -2,3551 | |
| Component | 9914_subgroup | GO:0032350 | regulation of hormone metabolic process | -5,702 | 3,777 | -1,8937 | |
| Component | 9914_subgroup | GO:0042445 | hormone metabolic process | -5,702 | 3,777 | -2,3193 | |
| Main | 31399_main | GO:0031399 | regulation of protein modification process | -5,136 | 1,158 | -4,5011 | |
| Component | 31399_subgroup | GO:0018193 | peptidyl-amino acid modification | -5,136 | 1,158 | -4,4267 | |
| Component | 31399_subgroup | GO:0032268 | regulation of cellular protein metabolic process | -5,136 | 1,158 | -4,0166 | |
| Component | 31399_subgroup | GO:0070647 | protein modification by small protein conjugation or removal | -5,136 | 1,158 | -2,2356 | |
| Component | 31399_subgroup | GO:0036211 | protein modification process | -5,136 | 1,158 | -8,974 | |
| Component | 31399_subgroup | GO:0006464 | cellular protein modification process | -5,136 | 1,158 | -8,974 | |
| Component | 31399_subgroup | GO:0006468 | protein phosphorylation | -5,136 | 1,158 | -3,4394 | |
| Main | 43620_main | GO:0043620 | regulation of DNA-templated transcription in response to stress | -4,881 | -3,897 | -3,0701 | |
| Main | 43618_main | GO:0043618 | regulation of transcription from RNA polymerase II promoter in response to stress | -4,842 | -4,016 | -3,4796 | |
| Component | 43618_subgroup | GO:0061418 | regulation of transcription from RNA polymerase II promoter in response to hypoxia | -4,842 | -4,016 | -2,6904 | |
| Component | 43618_subgroup | GO:0006367 | transcription initiation from RNA polymerase II promoter | -4,842 | -4,016 | -1,7762 | |
| Main | 6996_main | GO:0006996 | organelle organization | -2,082 | 1,841 | -4,939 | |
| Component | 6996_subgroup | GO:0016043 | cellular component organization | -2,082 | 1,841 | -5,4266 | |
| Component | 6996_subgroup | GO:0051276 | chromosome organization | -2,082 | 1,841 | -4,382 | |
| Component | 6996_subgroup | GO:0006325 | chromatin organization | -2,082 | 1,841 | -4,9046 | |

| | | | | | | |
|---|---|---|---|---|---|---|
| Component | 6996_subgroup | GO:1902275 | regulation of chromatin organization | -2,082 | 1,841 | -3,6599 |
| Component | 6996_subgroup | GO:0018394 | peptidyl-lysine acetylation | -2,082 | 1,841 | -1,9019 |
| Component | 6996_subgroup | GO:0018393 | internal peptidyl-lysine acetylation | -2,082 | 1,841 | -1,7478 |
| Component | 6996_subgroup | GO:0031056 | regulation of histone modification | -2,082 | 1,841 | -2,7728 |
| Component | 6996_subgroup | GO:1905269 | positive regulation of chromatin organization | -2,082 | 1,841 | -2,297 |
| Component | 6996_subgroup | GO:0016573 | histone acetylation | -2,082 | 1,841 | -1,7478 |
| Component | 6996_subgroup | GO:0016570 | histone modification | -2,082 | 1,841 | -4,7233 |
| Component | 6996_subgroup | GO:0016569 | covalent chromatin modification | -2,082 | 1,841 | -5,3006 |
| Component | 6996_subgroup | GO:0006473 | protein acetylation | -2,082 | 1,841 | -1,7627 |
| Component | 6996_subgroup | GO:0006475 | internal protein amino acid acetylation | -2,082 | 1,841 | -1,7478 |
| Component | 6996_subgroup | GO:0051128 | regulation of cellular component organization | -2,082 | 1,841 | -3,6411 |
| Main | 48869_main | GO:0048869 | cellular developmental process | -1,341 | 4,396 | -8,8855 |
| Component | 48869_subgroup | GO:0048513 | animal organ development | -1,341 | 4,396 | -6,492 |
| Component | 48869_subgroup | GO:0007399 | nervous system development | -1,341 | 4,396 | -3,8513 |
| Component | 48869_subgroup | GO:0048534 | hematopoietic or lymphoid organ development | -1,341 | 4,396 | -4,8855 |
| Component | 48869_subgroup | GO:0002520 | immune system development | -1,341 | 4,396 | -5,0795 |
| Component | 48869_subgroup | GO:0002521 | leukocyte differentiation | -1,341 | 4,396 | -1,9158 |
| Component | 48869_subgroup | GO:1903706 | regulation of hemopoiesis | -1,341 | 4,396 | -5,3235 |
| Component | 48869_subgroup | GO:0048598 | embryonic morphogenesis | -1,341 | 4,396 | -2,6243 |
| Component | 48869_subgroup | GO:0050793 | regulation of developmental process | -1,341 | 4,396 | -6,6412 |
| Component | 48869_subgroup | GO:0030218 | erythrocyte differentiation | -1,341 | 4,396 | -2,4066 |
| Component | 48869_subgroup | GO:0030219 | megakaryocyte differentiation | -1,341 | 4,396 | -1,894 |
| Component | 48869_subgroup | GO:0050778 | positive regulation of immune response | -1,341 | 4,396 | -1,9964 |
| Component | 48869_subgroup | GO:0050776 | regulation of immune response | -1,341 | 4,396 | -2,2497 |
| Component | 48869_subgroup | GO:0007275 | multicellular organism development | -1,341 | 4,396 | -7,8478 |
| Component | 48869_subgroup | GO:0061061 | muscle structure development | -1,341 | 4,396 | -2,0712 |
| Component | 48869_subgroup | GO:0045444 | fat cell differentiation | -1,341 | 4,396 | -2,1985 |
| Component | 48869_subgroup | GO:0048468 | cell development | -1,341 | 4,396 | -2,4759 |
| Component | 48869_subgroup | GO:0022008 | neurogenesis | -1,341 | 4,396 | -2,7078 |
| Component | 48869_subgroup | GO:0009653 | anatomical structure morphogenesis | -1,341 | 4,396 | -2,3334 |
| Component | 48869_subgroup | GO:0030855 | epithelial cell differentiation | -1,341 | 4,396 | -2,6243 |
| Component | 48869_subgroup | GO:0045646 | regulation of erythrocyte differentiation | -1,341 | 4,396 | -2,9455 |
| Component | 48869_subgroup | GO:0045648 | positive regulation of erythrocyte differentiation | -1,341 | 4,396 | -2,2122 |
| Component | 48869_subgroup | GO:0045637 | regulation of myeloid cell differentiation | -1,341 | 4,396 | -3,9159 |
| Component | 48869_subgroup | GO:0045639 | positive regulation of myeloid cell differentiation | -1,341 | 4,396 | -1,848 |
| Component | 48869_subgroup | GO:0030099 | myeloid cell differentiation | -1,341 | 4,396 | -3,8127 |
| Component | 48869_subgroup | GO:0030097 | hemopoiesis | -1,341 | 4,396 | -4,7486 |
| Component | 48869_subgroup | GO:0034101 | erythrocyte homeostasis | -1,341 | 4,396 | -2,4066 |
| Component | 48869_subgroup | GO:0002684 | positive regulation of immune system process | -1,341 | 4,396 | -2,6011 |
| Component | 48869_subgroup | GO:0002682 | regulation of immune system process | -1,341 | 4,396 | -4,5869 |
| Component | 48869_subgroup | GO:0002244 | hematopoietic progenitor cell differentiation | -1,341 | 4,396 | -2,2416 |
| Component | 48869_subgroup | GO:0045088 | regulation of innate immune response | -1,341 | 4,396 | -1,894 |
| Component | 48869_subgroup | GO:0030154 | cell differentiation | -1,341 | 4,396 | -9,3126 |
| Component | 48869_subgroup | GO:0051960 | regulation of nervous system development | -1,341 | 4,396 | -2,5218 |
| Component | 48869_subgroup | GO:0035295 | tube development | -1,341 | 4,396 | -1,7829 |
| Component | 48869_subgroup | GO:0048863 | stem cell differentiation | -1,341 | 4,396 | -2,0975 |
| Component | 48869_subgroup | GO:0048856 | anatomical structure development | -1,341 | 4,396 | -8,4132 |
| Component | 48869_subgroup | GO:0002262 | myeloid cell homeostasis | -1,341 | 4,396 | -2,1985 |
| Component | 48869_subgroup | GO:0009790 | embryo development | -1,341 | 4,396 | -1,9735 |
| Component | 48869_subgroup | GO:0030182 | neuron differentiation | -1,341 | 4,396 | -3,3421 |
| Component | 48869_subgroup | GO:0045595 | regulation of cell differentiation | -1,341 | 4,396 | -6,4163 |
| Component | 48869_subgroup | GO:0048699 | generation of neurons | -1,341 | 4,396 | -2,8874 |
| Component | 48869_subgroup | GO:0006955 | immune response | -1,341 | 4,396 | -4,2971 |
| Component | 48869_subgroup | GO:0007423 | sensory organ development | -1,341 | 4,396 | -2,3283 |
| Component | 48869_subgroup | GO:0007417 | central nervous system development | -1,341 | 4,396 | -2,9388 |
| Component | 48869_subgroup | GO:0007420 | brain development | -1,341 | 4,396 | -1,8954 |
| Component | 48869_subgroup | GO:0048731 | system development | -1,341 | 4,396 | -7,2271 |
| Component | 48869_subgroup | GO:0009913 | epidermal cell differentiation | -1,341 | 4,396 | -2,1424 |
| Component | 48869_subgroup | GO:0098751 | bone cell development | -1,341 | 4,396 | -1,836 |
| Main | 32502_main | GO:0032502 | developmental process | -1,109 | -0,152 | -9,0812 |
| Main | 51641_main | GO:0051641 | cellular localization | -0,97 | 6,803 | -2,4553 |
| Main | 71702_main | GO:0071702 | organic substance transport | -0,964 | 7,249 | -3,3708 |
| Component | 71702_subgroup | GO:0051234 | establishment of localization | -0,964 | 7,249 | -3,1151 |
| Component | 71702_subgroup | GO:0006810 | transport | -0,964 | 7,249 | -3,0149 |
| Main | 33036_main | GO:0033036 | macromolecule localization | -0,842 | 6,981 | -4,5869 |
| Main | 60992_main | GO:0060992 | response to fungicide | -0,751 | -7,372 | -2,3811 |
| Main | 2376_main | GO:0002376 | immune system process | -0,733 | -1,901 | -5,2489 |
| Main | 9299_main | GO:0009299 | mRNA transcription | -0,091 | -4,196 | -2,733 |
| Main | 8283_main | GO:0008283 | cell proliferation | 0,206 | 0,645 | -6,3332 |
| Main | 7623_main | GO:0007623 | circadian rhythm | 0,294 | -0,754 | -2,251 |
| Main | 32501_main | GO:0032501 | multicellular organismal process | 0,934 | -3,139 | -9,8699 |
| Main | 7154_main | GO:0007154 | cell communication | 1,484 | -1,519 | -6,7794 |
| Main | 51179_main | GO:0051179 | localization | 1,674 | 0,528 | -4,8735 |
| Main | 50896_main | GO:0050896 | response to stimulus | 2,1 | -4,92 | -10,3121 |
| Main | 51704_main | GO:0051704 | multi-organism process | 2,318 | -3,956 | -4,3234 |
| Main | 48511_main | GO:0048511 | rhythmic process | 2,599 | -5,955 | -3,5054 |
| Main | 71840_main | GO:0071840 | cellular component organization or biogenesis | 3,149 | -6,279 | -5,1037 |
| Main | 23052_main | GO:0023052 | signaling | 3,27 | -4,723 | -6,9592 |
| Main | 6366_main | GO:0006366 | transcription from RNA polymerase II promoter | 4,202 | -1,532 | -1,836 |

### SMURF2

| LOCUS: chr1 | chrom | chromStart | chromEnd | name | pubMedID | author | pubDate | journal | title | trait | region | pValue | Autoimmune relevant | Overlapping survivin-associated | Overlapping other SMURF2 RE |
|---|---|---|---|---|---|---|---|---|---|---|---|---|---|---|---|
| SMURF2 | chr17 | 64483155 | 64483156 | **rs17650301** | 32042192 | Ruth KS | 20-02-10 | Nat Med | Using human genetics to unde | Sex hormone-binding globulin levels adjusted for BMI | 17q23.3 | 1,00E-09 | | | |
| SMURF2 | chr17 | 64483155 | 64483156 | **rs17650301** | 32042192 | Ruth KS | 20-02-10 | Nat Med | Using human genetics to unde | Sex hormone-binding globulin levels | 17q23.3 | 3,00E-07 | | | |
| SMURF2 | chr17 | 64483155 | 64483156 | **rs17650301** | 30598549 | Morris JA | 18-12-31 | Nat Genet | An atlas of genetic influences | Heel bone mineral density | 17q23.3 | 2,00E-17 | | | |
| SMURF2 | chr17 | 64483155 | 64483156 | **rs17650301** | 30595370 | Kichaev G | 18-12-27 | Am J Hum G | Leveraging Polygenic Function | Heel bone mineral density | 17q23.3 | 3,00E-23 | | | |
| SMURF2 | chr17 | 64483155 | 64483156 | **rs17650301** | 30595370 | Kichaev G | 18-12-27 | Am J Hum G | Leveraging Polygenic Function | Age at menopause | 17q23.3 | 6,00E-20 | | | |
| SMURF2 | chr17 | 64483155 | 64483156 | **rs17650301** | 30048462 | Kim SK | 18-07-26 | PLoS One | Identification of 613 new loci | Heel bone mineral density | 17q23.3 | 2,00E-17 | | | |
| SMURF2 | chr17 | 64487813 | 64487814 | **rs113252144** | 30595370 | Kichaev G | 18-12-27 | Am J Hum G | Leveraging Polygenic Function | Height | 17q23.3 | 2,00E-27 | | | |
| SMURF2 | chr17 | 64501845 | 64501846 | **rs77672322** | 30804560 | Shrine N | 19-02-25 | Nat Genet | New genetic signals for lung f | Lung function (FVC) | 17q23.3 | 3,00E-09 | | | |
| SMURF2 | chr17 | 64506316 | 64506317 | **rs1991401** | 32296059 | Han Y | 20-04-15 | Nat Commun | Genome-wide analysis highlig | Asthma | 17q23.3 | 1,00E-08 | Y | | |
| SMURF2 | chr17 | 64506316 | 64506317 | **rs1991401** | 32231278 | Hysi PG | 20-03-30 | Nat Genet | Meta-analysis of 542,934 subj | Refractive error | 17q23.3 | 2,00E-13 | | | |
| SMURF2 | chr17 | 64506316 | 64506317 | **rs1991401** | 32042192 | Ruth KS | 20-02-10 | Nat Med | Using human genetics to unde | Sex hormone-binding globulin levels adjusted for BMI | 17q23.3 | 1,00E-08 | | | |
| SMURF2 | chr17 | 64506316 | 64506317 | **rs1991401** | 32042192 | Ruth KS | 20-02-10 | Nat Med | Using human genetics to unde | Sex hormone-binding globulin levels | 17q23.3 | 2,00E-07 | | | |
| SMURF2 | chr17 | 64506316 | 64506317 | **rs1991401** | 30595370 | Kichaev G | 18-12-27 | Am J Hum G | Leveraging Polygenic Function | Systolic blood pressure | 17q23.3 | 3,00E-12 | | | |
| SMURF2 | chr17 | 64510161 | 64510162 | **rs57166100** | 24324551 | Deng X | 13-11-20 | PLoS One | Genome wide association stu | PR interval in Tripanosoma cruzi seropositivity | 17q23.3 | 3,00E-06 | | | |
| SMURF2 | chr17 | 64520840 | 64520841 | **rs148050755** | 31014085 | Welsh P | 19-04-24 | Circulation | Cardiac Troponin T and Tropor | Cardiac troponin-I levels | 17q23.3 | 3,00E-11 | | | |
| SMURF2 | chr17 | 64539477 | 64539478 | **rs139126928** | 32572223 | Hughes DA | 20-06-22 | Nat Microbio | Genome-wide associations of | Gut microbiota (bacterial taxa, hurdle binary method) | 17q23.3 | 2,00E-06 | | | |
| SMURF2 | chr17 | 64593459 | 64593460 | **rs80215473** | 30595370 | Kichaev G | 18-12-27 | Am J Hum G | Leveraging Polygenic Function | Height | 17q23.3 | 8,00E-10 | | | |
| SMURF2 | chr17 | 64604816 | 64604817 | **rs111854052** | 24024966 | Teumer A | 13-08-14 | J Clin Periodo | Genome-wide association stu | Periodontitis (Mean PAL) | 17q24.1 | 9,00E-06 | | | |
| SMURF2 | chr17 | 64680030 | 64680031 | **rs6504249** | 27182965 | Pickrell JK | 16-05-16 | Nat Genet | Detection and interpretation d | Joint mobility (Beighton score) | 17q24.1 | 9,00E-09 | Y | Y: SMURF2_GH17F064678 | |
| SMURF2 | chr17 | 64685386 | 64685387 | **rs11656678** | 30595370 | Kichaev G | 18-12-27 | Am J Hum G | Leveraging Polygenic Function | Lung function (FVC) | 17q24.1 | 4,00E-16 | | | |
| SMURF2 | chr17 | 64690611 | 64690612 | **rs11653958** | 30804560 | Shrine N | 19-02-25 | Nat Genet | New genetic signals for lung f | Lung function (FVC) | 17q24.1 | 3,00E-10 | | | |
| SMURF2 | chr17 | 64690611 | 64690612 | **rs11653958** | 30804560 | Shrine N | 19-02-25 | Nat Genet | New genetic signals for lung f | Lung function (FEV1/FVC) | 17q24.1 | 9,00E-13 | | | |
| SMURF2 | chr17 | 64743455 | 64743456 | **rs9900062** | 30595370 | Kichaev G | 18-12-27 | Am J Hum G | Leveraging Polygenic Function | Lung function (FEV1/FVC) | 17q24.1 | 4,00E-17 | | | |
| SMURF2 | chr17 | 64743455 | 64743456 | **rs9900062** | 29213071 | Mendez-Gira | 17-12-06 | Sci Rep | GWAS of the electrocardiogra | QT interval | 17q24.1 | 5,00E-06 | | | |

### BIRC2

| LOCUS: chr1 | chrom | chromStart | chromEnd | name | pubMedID | author | pubDate | journal | title | trait | region | pValue | Autoimmune relevant | Overlapping survivin-associated | Overlapping other BIRC2 RE |
|---|---|---|---|---|---|---|---|---|---|---|---|---|---|---|---|
| BIRC2 | chr11 | 102240709 | 102240710 | **rs117751356** | 30589442 | Whitfield JB | 18-12-27 | Alcohol Clin | Biomarker and genomic risk fa | Alanine aminotransferase levels in low alcohol consumption | 11q22.1 | 4,00E-06 | | | |
| BIRC2 | chr11 | 102284742 | 102284743 | **rs2852894** | 21116278 | Furney SJ | 10-11-30 | Mol Psychiat | Genome-wide association wit | Whole-brain volume (Alzheimer's disease interaction) | 11q22.1 | 1,00E-06 | | | |
| BIRC2 | chr11 | 102287168 | 102287169 | **rs4754839** | 32042192 | Ruth KS | 20-02-10 | Nat Med | Using human genetics to unde | Total testosterone levels | 11q22.1 | 1,00E-08 | | | |
| BIRC2 | chr11 | 102320216 | 102320217 | **rs186356272** | 33067605 | Folkersen L | 20-10-16 | Nat Metab | Genomic and drug target eval | Matrix metalloproteinase-1 levels | 11q22.2 | 2,00E-15 | | | Y: BIRC2_GH11F102311 |
| BIRC2 | chr11 | 102410019 | 102410020 | **rs11225247** | 25881214 | Shah AA | 15-02-27 | Heart Surg F | Genetic variants associated w | Vein graft stenosis in coronary artery bypass grafting | 11q22.2 | 8,00E-06 | | | |
| BIRC2 | chr11 | 102503235 | 102503236 | **rs7131148** | 32579418 | Maukonen M | 20-06-24 | J Biol Rhythn | Genetic Associations of Chron | Chronotype (sMEQ score) | 11q22.2 | 2,00E-06 | | | |
| BIRC2 | chr11 | 102510592 | 102510593 | **rs578627** | 33632238 | Song Y | 21-02-25 | J Transl Med | Identification of susceptibility | Cardio-cerebrovascular disease in dyslipidemia | 11q22.2 | 1,00E-05 | | | |
| BIRC2 | chr11 | 102525875 | 102525876 | **rs12285347** | 28139693 | Hoffmann TJ | 17-01-31 | Nat Commun | Genome-wide association stu | Prostate-specific antigen levels | 11q22.2 | 8,00E-12 | | | |
| BIRC2 | chr11 | 102527702 | 102527703 | **rs17884405** | 33067605 | Folkersen L | 20-10-16 | Nat Metab | Genomic and drug target eval | Matrix metalloproteinase-7 levels | 11q22.2 | 6,00E-28 | | | |
| BIRC2 | chr11 | 102530901 | 102530902 | **rs11568819** | 30072576 | Emilsson V | 18-08-02 | Science | Co-regulatory networks of hur | Blood protein levels | 11q22.2 | 5,00E-21 | | | |
| BIRC2 | chr11 | 102530901 | 102530902 | **rs11568819** | 28915241 | Ahsan M | 17-09-15 | PLoS Genet | The relative contribution of DI | Blood protein levels | 11q22.2 | 4,00E-12 | | | |
| BIRC2 | chr11 | 102530901 | 102530902 | **rs11568819** | 28240269 | Suhre K | 17-02-27 | Nat Commun | Connecting genetic risk to dise | Blood protein levels | 11q22.2 | 6,00E-14 | | | |
| BIRC2 | chr11 | 102530901 | 102530902 | **rs11568819** | 33067605 | Folkersen L | 20-10-16 | Nat Metab | Genomic and drug target eval | Matrix metalloproteinase-7 levels | 11q22.2 | 2,00E-222 | | | |
| BIRC2 | chr11 | 102530929 | 102530930 | **rs11568818** | 28915241 | Ahsan M | 17-09-15 | PLoS Genet | The relative contribution of DI | Blood protein levels | 11q22.2 | 4,00E-12 | | | |
| BIRC2 | chr11 | 102530929 | 102530930 | **rs11568818** | 33398198 | Conti DV | 21-01-04 | Nat Genet | Trans-ancestry genome-wide | Prostate cancer | 11q22.2 | 4,00E-26 | | | |
| BIRC2 | chr11 | 102530929 | 102530930 | **rs11568818** | 29892016 | Schumacher | 18-06-11 | Nat Genet | Association analyses of more | Prostate cancer | 11q22.2 | 2,00E-19 | | | |
| BIRC2 | chr11 | 102530929 | 102530930 | **rs11568818** | 23535732 | Eeles RA | 13-04-01 | Nat Genet | Identification of 23 new prost | Prostate cancer | 11q22.2 | 2,00E-11 | | | |
| BIRC2 | chr11 | 102544269 | 102544270 | **rs7946641** | 33067605 | Folkersen L | 20-10-16 | Nat Metab | Genomic and drug target eval | Matrix metalloproteinase-7 levels | 11q22.2 | 1,00E-44 | | | |
| BIRC2 | chr11 | 102575642 | 102575643 | **rs11607749** | 33067605 | Folkersen L | 20-10-16 | Nat Metab | Genomic and drug target eval | Matrix metalloproteinase-7 levels | 11q22.2 | 1,00E-13 | | | |
| BIRC2 | chr11 | 102591594 | 102591595 | **rs61893841** | 30067105 | Chang LC | 18-08-01 | Expert Opin | Thyrotropin receptor antibodie | Adverse response to antithyroid drugs in Graves disease | 11q22.2 | 9,00E-07 | | | |
| BIRC2 | chr11 | 102599524 | 102599525 | **rs10895322** | 28924153 | Chang X | 17-09-18 | Nat Commun | Common variants in MMP20 a | Neuroblastoma (11q deletion) | 11q22.2 | 3,00E-10 | | | Y: BIRC2_GH11F102597 |
| BIRC2 | chr11 | 102609658 | 102609659 | **rs17098961** | 27126917 | Spada J | 16-04-29 | J Sleep Res | Genome-wide association ana | Night sleep phenotypes | 11q22.2 | 7,00E-07 | | | |
| BIRC2 | chr11 | 102617341 | 102617342 | **rs1317947** | 29545352 | Li Y | 18-03-15 | J Am Soc Ne | Genome-Wide Association St | Urinary metabolite levels in chronic kidney disease | 11q22.2 | 5,00E-08 | | | |
| BIRC2 | chr11 | 102620517 | 102620517 | **rs11383572** | 33067605 | Folkersen L | 20-10-16 | Nat Metab | Genomic and drug target eval | Matrix metalloproteinase-1 levels | 11q22.2 | 7,00E-18 | | | |
| BIRC2 | chr11 | 102630933 | 102630934 | **rs948138** | 30166627 | Massey J | 18-08-31 | Pharmacoge | Genome-wide association stu | Response to TNF inhibitor in rheumatoid arthritis (change in patient global heat | 11q22.2 | 8,00E-08 | Y | | |
| BIRC2 | chr11 | 102632171 | 102632172 | **rs183501923** | 31250787 | Fawns-Ritch | 19-06-01 | Twin Res Hu | Genetic Contributions to Healt | Health literacy | 11q22.2 | 2,00E-06 | | | |
| BIRC2 | chr11 | 102636683 | 102636684 | **rs2260433** | 28054174 | Leslie EJ | 17-01-04 | Hum Genet | Genome-wide meta-analyses | Cleft palate | 11q22.2 | 9,00E-06 | | | |
| BIRC2 | chr11 | 102638968 | 102638969 | **rs139780693** | 31591132 | Wu W | 19-10-07 | Genetics | Retrospective Association Ana | Longitudinal cocaine use | 11q22.2 | 4,00E-07 | | | |
| BIRC2 | chr11 | 102638968 | 102638969 | **rs139780693** | 31591132 | Wu W | 19-10-07 | Genetics | Retrospective Association Ana | Longitudinal cocaine use | 11q22.2 | 2,00E-07 | | | |
| BIRC2 | chr11 | 102644129 | 102644130 | **rs4433535** | 27532455 | Sun W | 16-08-17 | PLoS Genet | Common Genetic Polymorphis | Blood protein levels | 11q22.2 | 7,00E-07 | | | |
| BIRC2 | chr11 | 102655878 | 102655879 | **rs2846714** | 30072576 | Emilsson V | 18-08-02 | Science | Co-regulatory networks of hur | Blood protein levels | 11q22.2 | 2,00E-14 | | | |
| BIRC2 | chr11 | 102685589 | 102685590 | **rs1291326** | 33067605 | Folkersen L | 20-10-16 | Nat Metab | Genomic and drug target eval | Matrix metalloproteinase-3 levels | 11q22.2 | 2,00E-38 | | | |
| BIRC2 | chr11 | 102724403 | 102724404 | **rs1939012** | 25087078 | Anney RJ | 14-07-30 | Lancet Neuro | Genetic determinants of comr | Epilepsy | 11q22.2 | 2,00E-08 | | | |
| BIRC2 | chr11 | 102738267 | 102738268 | **rs55930209** | 33067605 | Folkersen L | 20-10-16 | Nat Metab | Genomic and drug target eval | Matrix metalloproteinase-10 levels | 11q22.2 | 4,00E-18 | | | |
| BIRC2 | chr11 | 102757320 | 102757321 | **rs7924357** | 23412934 | Gourraud PA | 13-02-13 | Brain | A genome-wide association st | Multiple sclerosis | 11q22.2 | 1,00E-06 | Y | | |
| BIRC2 | chr11 | 102771529 | 102771530 | **rs12290253** | 33067605 | Folkersen L | 20-10-16 | Nat Metab | Genomic and drug target eval | Matrix metalloproteinase-1 levels | 11q22.2 | 1,00E-48 | | | |
| BIRC2 | chr11 | 102771778 | 102771779 | **rs12807063** | 30072576 | Emilsson V | 18-08-02 | Science | Co-regulatory networks of hur | Blood protein levels | 11q22.2 | 4,00E-21 | | | |
| BIRC2 | chr11 | 102777325 | 102777326 | **rs17860967** | 30072576 | Emilsson V | 18-08-02 | Science | Co-regulatory networks of hur | Blood protein levels | 11q22.2 | 3,00E-15 | | | |
| BIRC2 | chr11 | 102778750 | 102778751 | **rs17860955** | 29875488 | Sun BB | 18-06-06 | Nature | Genomic atlas of the human p | Blood protein levels | 11q22.2 | 1,00E-23 | | | |
| BIRC2 | chr11 | 102778750 | 102778751 | **rs17860955** | 28915241 | Ahsan M | 17-09-15 | PLoS Genet | The relative contribution of DI | Blood protein levels | 11q22.2 | 5,00E-10 | | | |
| BIRC2 | chr11 | 102778750 | 102778751 | **rs17860955** | 33067605 | Folkersen L | 20-10-16 | Nat Metab | Genomic and drug target eval | Matrix metalloproteinase-10 levels | 11q22.2 | 5,00E-155 | | | |
| BIRC2 | chr11 | 102779692 | 102779693 | **rs486055** | 29875488 | Sun BB | 18-06-06 | Nature | Genomic atlas of the human p | Blood protein levels | 11q22.2 | 1,00E-21 | | | |
| BIRC2 | chr11 | 102779692 | 102779693 | **rs486055** | 28915241 | Ahsan M | 17-09-15 | PLoS Genet | The relative contribution of DI | Blood protein levels | 11q22.2 | 9,00E-10 | | | |
| BIRC2 | chr11 | 102779692 | 102779693 | **rs486055** | 33067605 | Folkersen L | 20-10-16 | Nat Metab | Genomic and drug target eval | Matrix metalloproteinase-10 levels | 11q22.2 | 9,00E-142 | | | |
| BIRC2 | chr11 | 102779692 | 102779693 | **rs486055** | 32641083 | Hillary RF | 20-07-08 | Genome Me | Multi-method genome- and e | Matrix metalloproteinase-10 levels | 11q22.2 | 2,00E-11 | | | |
| BIRC2 | chr11 | 102795923 | 102795924 | **rs17878931** | 30072576 | Emilsson V | 18-08-02 | Science | Co-regulatory networks of hur | Blood protein levels | 11q22.2 | 1,00E-42 | | | |
| BIRC2 | chr11 | 102800965 | 102800966 | **rs534191** | 32641083 | Hillary RF | 20-07-08 | Genome Me | Multi-method genome- and e | Matrix metalloproteinase levels | 11q22.2 | 6,00E-16 | | | |

### PFKFB3

| LOCUS: chr1 | chrom | chromStart | chromEnd | name | pubMedID | author | pubDate | journal | title | trait | region | pValue | Autoimmune relevant | Overlapping survivin-associated | Overlapping other PFKFB3 RE |
|---|---|---|---|---|---|---|---|---|---|---|---|---|---|---|---|
| PFKFB3 | chr10 | 6000051 | 6000052 | **rs76913543** | 26974007 | Ellinghaus D | 16-03-14 | Nat Genet | Analysis of five chronic inflam | Chronic inflammatory diseases (ankylosing spondylitis, Crohn's disease, psoriasis | 10p15.1 | 4,00E-12 | Y | | |
| PFKFB3 | chr10 | 6011199 | 6011200 | **rs12722605** | 22228203 | Aslibekyan S | 12-01-05 | Pharmacoge | A genome-wide association st | Inflammatory biomarkers | 10p15.1 | 3,00E-07 | | | |
| PFKFB3 | chr10 | 6013258 | 6013259 | **rs11256335** | 31596458 | Greenwood T | 19-10-09 | JAMA Psychi | Genome-wide Association of | Degraded stimulus continuous performance test score | 10p15.1 | 3,00E-06 | | | |
| PFKFB3 | chr10 | 6016359 | 6016360 | **rs4749894** | 32296059 | Han Y | 20-04-15 | Nat Commun | Genome-wide analysis highlig | Asthma | 10p15.1 | 4,00E-17 | Y | | |
| PFKFB3 | chr10 | 6018469 | 6018470 | **rs12722588** | 27989323 | Ahola-Olli A | 16-12-13 | Am J Hum G | Genome-wide Association Stu | Interleukin-2 receptor antagonist levels | 10p15.1 | 9,00E-20 | Y | | |
| PFKFB3 | chr10 | 6024498 | 6024499 | **rs12722574** | 31604244 | International | 19-09-01 | Science | Multiple sclerosis genomic ma | Multiple sclerosis | 10p15.1 | 4,00E-07 | Y | | |
| PFKFB3 | chr10 | 6025724 | 6025725 | **rs2025345** | 23817569 | Hinds DA | 13-06-30 | Nat Genet | A genome-wide association m | Self-reported allergy | 10p15.1 | 3,00E-06 | Y | | |
| PFKFB3 | chr10 | 6027929 | 6027930 | **rs12722561** | 26819262 | Zhou Y | 16-01-27 | Mult Scler | Genetic loci for Epstein-Barr v | Epstein Barr virus nuclear antigen 1 IgG levels or multiple sclerosis | 10p15.1 | 6,00E-08 | Y | | |
| PFKFB3 | chr10 | 6027929 | 6027930 | **rs12722561** | 32953199 | Liu Z | 19-03-01 | J BioX Res | Integrative omics analysis ide | Liver fibrogenesis (alpha smooth muscle actin levels) | 10p15.1 | 2,00E-06 | | | |
| PFKFB3 | chr10 | 6028309 | 6028310 | **rs12722559** | 31604244 | International | 19-09-01 | Science | Multiple sclerosis genomic ma | Multiple sclerosis | 10p15.1 | 2,00E-15 | Y | | |
| PFKFB3 | chr10 | 6028575 | 6028576 | **rs12722556** | 25918132 | Yucesoy B | 15-04-26 | Toxicol Sci | Genome-Wide Association St | Diisocyanate-induced asthma | 10p15.1 | 2,00E-06 | | | |
| PFKFB3 | chr10 | 6030129 | 6030130 | **rs12722547** | 31361310 | Johansson A | 19-07-30 | Hum Mol Ge | Genome-wide association ana | Asthma | 10p15.1 | 6,00E-21 | Y | | |
| PFKFB3 | chr10 | 6030129 | 6030130 | **rs12722547** | 31669095 | Zhu Z | 19-10-24 | J Allergy Clin | Shared Genetic and Experime | Asthma (childhood onset) | 10p15.1 | 4,00E-09 | Y | | |
| PFKFB3 | chr10 | 6030129 | 6030130 | **rs12722547** | 32888493 | Chen MH | 20-09-01 | Cell | Trans-ethnic and Ancestry-Spe | Eosinophil counts | 10p15.1 | 6,00E-20 | Y | | |
| PFKFB3 | chr10 | 6030129 | 6030130 | **rs12722547** | 32888493 | Chen MH | 20-09-01 | Cell | Trans-ethnic and Ancestry-Spe | Eosinophil counts | 10p15.1 | 4,00E-23 | Y | | |
| PFKFB3 | chr10 | 6030129 | 6030130 | **rs12722547** | 32888494 | Vuckovic D | 20-09-01 | Cell | The Polygenic and Monogenic | Eosinophil percentage of white cells | 10p15.1 | 2,00E-15 | Y | | |
| PFKFB3 | chr10 | 6030129 | 6030130 | **rs12722547** | 31669095 | Zhu Z | 19-10-24 | J Allergy Clin | Shared Genetic and Experime | Nonatopic asthma | 10p15.1 | 6,00E-09 | Y | | |
| PFKFB3 | chr10 | 6030129 | 6030130 | **rs12722547** | 30595370 | Kichaev G | 18-12-27 | Am J Hum G | Leveraging Polygenic Function | Respiratory diseases | 10p15.1 | 3,00E-11 | | | |
| PFKFB3 | chr10 | 6032487 | 6032488 | **rs4747846** | 29083406 | Ferreira MA | 17-10-30 | Nat Genet | Shared genetic origin of asthm | Allergic disease (asthma, hay fever or eczema) | 10p15.1 | 1,00E-11 | Y | | |
| PFKFB3 | chr10 | 6037071 | 6037072 | **rs7893467** | 30595370 | Kichaev G | 18-12-27 | Am J Hum G | Leveraging Polygenic Function | Hypothyroidism | 10p15.1 | 3,00E-08 | Y | | |
| PFKFB3 | chr10 | 6037071 | 6037072 | **rs7893467** | 31015401 | Wu Y | 19-04-23 | Nat Commun | Genome-wide association stu | Medication use (thyroid preparations) | 10p15.1 | 3,00E-09 | | | |
| PFKFB3 | chr10 | 6039266 | 6039267 | **rs12722515** | 26192919 | Liu JZ | 15-07-20 | Nat Genet | Association analyses identify 3 | Crohn's disease | 10p15.1 | 8,00E-13 | Y | | |
| PFKFB3 | chr10 | 6039266 | 6039267 | **rs12722515** | 26192919 | Liu JZ | 15-07-20 | Nat Genet | Association analyses identify 3 | Inflammatory bowel disease | 10p15.1 | 5,00E-12 | Y | | |
| PFKFB3 | chr10 | 6039266 | 6039267 | **rs12722515** | 23128233 | Jostins L | 12-11-01 | Nature | Host-microbe interactions hav | Inflammatory bowel disease | 10p15.1 | 4,00E-10 | Y | | |
| PFKFB3 | chr10 | 6040594 | 6040595 | **rs11256464** | 28928442 | Tian C | 17-09-19 | Nat Commun | Genome-wide association and | Tonsillectomy | 10p15.1 | 7,00E-06 | | | |
| PFKFB3 | chr10 | 6040989 | 6040990 | **rs6602398** | 27643478 | Powers A | 16-09-09 | J Psychiatr R | A genome-wide association st | Emotional dysregulation | 10p15.1 | 1,00E-08 | | | |
| PFKFB3 | chr10 | 6045331 | 6045332 | **rs1107345** | 31407831 | Din L | 19-08-13 | Genet Epide | Genetic overlap between auto | Marginal zone lymphoma or multiple sclerosis | 10p15.1 | 3,00E-08 | Y | | |
| PFKFB3 | chr10 | 6047877 | 6047878 | **rs7909519** | 23603761 | Hinks A | 13-04-21 | Nat Genet | Dense genotyping of immune- | Juvenile idiopathic arthritis (oligoarticular or rheumatoid factor-negative polyart | 10p15.1 | 8,00E-10 | Y | | |
| PFKFB3 | chr10 | 6048358 | 6048359 | **rs791590** | 26293465 | Durda P | 15-08-20 | Arterioscler T | Plasma Levels of Soluble Inter | Soluble interleukin-2 receptor subunit alpha | 10p15.1 | 7,00E-35 | Y | | |
| PFKFB3 | chr10 | 6049053 | 6049054 | **rs7090504** | 27863252 | Astle WJ | 16-11-17 | Cell | The Allelic Landscape of Huma | Lymphocyte counts | 10p15.1 | 2,00E-15 | Y | | |
| PFKFB3 | chr10 | 6050382 | 6050383 | **rs12253981** | 28445677 | Steri M | 17-04-27 | N Engl J Med | Overexpression of the Cytokin | Multiple sclerosis | 10p15.1 | 1,00E-09 | Y | | |
| PFKFB3 | chr10 | 6051175 | 6051176 | **rs12722502** | 32296059 | Han Y | 20-04-15 | Nat Commun | Genome-wide analysis highlig | Asthma | 10p15.1 | 1,00E-24 | Y | | |
| PFKFB3 | chr10 | 6051175 | 6051176 | **rs12722502** | 31959851 | Olafsdottir T | 20-01-20 | Nat Commun | Eighty-eight variants highlight | Asthma | 10p15.1 | 1,00E-17 | Y | | |
| PFKFB3 | chr10 | 6051175 | 6051176 | **rs12722502** | 31619474 | Zhu Z | 19-10-17 | Eur Respir J | Shared Genetics of Asthma ar | Asthma | 10p15.1 | 6,00E-17 | Y | | |
| PFKFB3 | chr10 | 6051175 | 6051176 | **rs12722502** | 30929738 | Ferreira MA | 19-03-28 | Am J Hum G | Genetic Architectures of Child | Asthma | 10p15.1 | 2,00E-18 | Y | | |
| PFKFB3 | chr10 | 6051175 | 6051176 | **rs12722502** | 31669095 | Zhu Z | 19-10-24 | J Allergy Clin | Shared Genetic and Experime | Asthma (adult onset) | 10p15.1 | 3,00E-10 | Y | | |
| PFKFB3 | chr10 | 6051175 | 6051176 | **rs12722502** | 30929738 | Ferreira MA | 19-03-28 | Am J Hum G | Genetic Architectures of Child | Asthma (adult onset) | 10p15.1 | 4,00E-12 | Y | | |
| PFKFB3 | chr10 | 6051175 | 6051176 | **rs12722502** | 30929738 | Ferreira MA | 19-03-28 | Am J Hum G | Genetic Architectures of Child | Asthma (childhood onset) | 10p15.1 | 7,00E-14 | Y | | |
| PFKFB3 | chr10 | 6051175 | 6051176 | **rs12722502** | 31669095 | Zhu Z | 19-10-24 | J Allergy Clin | Shared Genetic and Experime | Atopic asthma | 10p15.1 | 9,00E-14 | Y | | |
| PFKFB3 | chr10 | 6051175 | 6051176 | **rs12722502** | 32888494 | Vuckovic D | 20-09-01 | Cell | The Polygenic and Monogenic | Eosinophil counts | 10p15.1 | 1,00E-17 | Y | | |
| PFKFB3 | chr10 | 6051175 | 6051176 | **rs12722502** | 30595370 | Kichaev G | 18-12-27 | Am J Hum G | Leveraging Polygenic Function | Eosinophil counts | 10p15.1 | 8,00E-24 | Y | | |
| PFKFB3 | chr10 | 6051175 | 6051176 | **rs12722502** | 31361310 | Johansson A | 19-07-30 | Hum Mol Ge | Genome-wide association ana | Hay fever and/or eczema | 10p15.1 | 3,00E-18 | Y | | |
| PFKFB3 | chr10 | 6052733 | 6052734 | **rs61839660** | 31361310 | Johansson A | 19-07-30 | Hum Mol Ge | Genome-wide association ana | Allergic disease (asthma, hay fever or eczema) | 10p15.1 | 3,00E-28 | Y | | |
| PFKFB3 | chr10 | 6052733 | 6052734 | **rs61839660** | 29785011 | Zhu Z | 18-05-21 | Nat Genet | A genome-wide cross-trait an | Allergic disease (asthma, hay fever or eczema) | 10p15.1 | 4,00E-12 | Y | | |
| PFKFB3 | chr10 | 6052733 | 6052734 | **rs61839660** | 29083406 | Ferreira MA | 17-10-30 | Nat Genet | Shared genetic origin of asthm | Allergic disease (asthma, hay fever or eczema) | 10p15.1 | 4,00E-19 | Y | | |
| PFKFB3 | chr10 | 6052733 | 6052734 | **rs61839660** | 29785011 | Zhu Z | 18-05-21 | Nat Genet | A genome-wide cross-trait an | Asthma or allergic disease (pleiotropy) | 10p15.1 | 2,00E-11 | Y | | |
| PFKFB3 | chr10 | 6052733 | 6052734 | **rs61839660** | 26974007 | Ellinghaus D | 16-03-14 | Nat Genet | Analysis of five chronic inflam | Chronic inflammatory diseases (ankylosing spondylitis, Crohn's disease, psoriasis | 10p15.1 | 1,00E-15 | Y | | |
| PFKFB3 | chr10 | 6052733 | 6052734 | **rs61839660** | 28067908 | de Lange KM | 17-01-09 | Nat Genet | Genome-wide association stu | Crohn's disease | 10p15.1 | 2,00E-08 | Y | | |
| PFKFB3 | chr10 | 6052733 | 6052734 | **rs61839660** | 31361310 | Johansson A | 19-07-30 | Hum Mol Ge | Genome-wide association ana | Eczema | 10p15.1 | 6,00E-14 | Y | | |
| PFKFB3 | chr10 | 6052733 | 6052734 | **rs61839660** | 30595370 | Kichaev G | 18-12-27 | Am J Hum G | Leveraging Polygenic Function | Eczema | 10p15.1 | 4,00E-38 | Y | | |
| PFKFB3 | chr10 | 6052733 | 6052734 | **rs61839660** | 31361310 | Johansson A | 19-07-30 | Hum Mol Ge | Genome-wide association ana | Hay fever and/or eczema | 10p15.1 | 1,00E-32 | Y | | |
| PFKFB3 | chr10 | 6052733 | 6052734 | **rs61839660** | 32888494 | Vuckovic D | 20-09-01 | Cell | The Polygenic and Monogenic | Lymphocyte counts | 10p15.1 | 3,00E-23 | Y | | |
| PFKFB3 | chr10 | 6052733 | 6052734 | **rs61839660** | 32888494 | Vuckovic D | 20-09-01 | Cell | The Polygenic and Monogenic | Lymphocyte percentage of white cells | 10p15.1 | 5,00E-29 | Y | | |
| PFKFB3 | chr10 | 6052733 | 6052734 | **rs61839660** | 27863252 | Astle WJ | 16-11-17 | Cell | The Allelic Landscape of Huma | Lymphocyte percentage of white cells | 10p15.1 | 2,00E-10 | Y | | |
| PFKFB3 | chr10 | 6052733 | 6052734 | **rs61839660** | 32888494 | Vuckovic D | 20-09-01 | Cell | The Polygenic and Monogenic | Monocyte count | 10p15.1 | 3,00E-23 | Y | | |
| PFKFB3 | chr10 | 6052733 | 6052734 | **rs61839660** | 32888494 | Vuckovic D | 20-09-01 | Cell | The Polygenic and Monogenic | Neutrophil percentage of white cells | 10p15.1 | 3,00E-14 | Y | | |
| PFKFB3 | chr10 | 6052733 | 6052734 | **rs61839660** | 28714469 | Langefeld CD | 17-07-17 | Nat Commun | Transancestral mapping and g | Systemic lupus erythematosus | 10p15.1 | 9,00E-06 | Y | | |
| PFKFB3 | chr10 | 6052733 | 6052734 | **rs61839660** | 25751624 | Onengut-Gu | 15-04-01 | Nat Genet | Fine mapping of type 1 diabet | Type 1 diabetes | 10p15.1 | 3,00E-39 | Y | | |
| PFKFB3 | chr10 | 6052733 | 6052734 | **rs61839660** | 22293688 | Huang J | 12-02-01 | Eur J Hum G | 1000 Genomes-based imputa | Type 1 diabetes | 10p15.1 | 5,00E-09 | Y | | |
| PFKFB3 | chr10 | 6053446 | 6053447 | **rs2476491** | 23603761 | Hinks A | 13-04-21 | Nat Genet | Dense genotyping of immune- | Juvenile idiopathic arthritis (oligoarticular or rheumatoid factor-negative polyart | 10p15.1 | 1,00E-06 | Y | | |
| PFKFB3 | chr10 | 6053964 | 6053965 | **rs12722497** | 28915241 | Ahsan M | 17-09-15 | PLoS Genet | The relative contribution of DI | Blood protein levels | 10p15.1 | 8,00E-16 | Y | | |
| PFKFB3 | chr10 | 6053964 | 6053965 | **rs12722497** | 27989323 | Ahola-Olli A | 16-12-13 | Am J Hum G | Genome-wide Association Stu | Interleukin-2 receptor antagonist levels | 10p15.1 | 2,00E-38 | Y | | |
| PFKFB3 | chr10 | 6053964 | 6053965 | **rs12722497** | 28928442 | Tian C | 17-09-19 | Nat Commun | Genome-wide association and | Strep throat | 10p15.1 | 5,00E-07 | | | |
| PFKFB3 | chr10 | 6054235 | 6054236 | **rs10795763** | 28714469 | Langefeld CD | 17-07-17 | Nat Commun | Transancestral mapping and g | Systemic lupus erythematosus | 10p15.1 | 2,00E-06 | Y | | |
| PFKFB3 | chr10 | 6055319 | 6055320 | **rs12722495** | 21829393 | Plagnol V | 11-08-04 | PLoS Genet | Genome-wide association ana | Type 1 diabetes | 10p15.1 | 1,00E-38 | Y | | |
| PFKFB3 | chr10 | 6056860 | 6056861 | **rs706779** | 22922229 | Cooper JD | 12-12-01 | Hum Mol Ge | Seven newly identified loci fo | Autoimmune thyroid diseases (Graves disease or Hashimoto's thyroiditis) | 10p15.1 | 3,00E-07 | Y | | |
| PFKFB3 | chr10 | 6056860 | 6056861 | **rs706779** | 22922229 | Cooper JD | 12-12-01 | Hum Mol Ge | Seven newly identified loci fo | Graves' disease | 10p15.1 | 2,00E-06 | Y | | |
| PFKFB3 | chr10 | 6056860 | 6056861 | **rs706779** | 27723757 | Jin Y | 16-11-01 | Nat Genet | Genome-wide association stu | Vitiligo | 10p15.1 | 7,00E-27 | | | |
| PFKFB3 | chr10 | 6056860 | 6056861 | **rs706779** | 20410501 | Jin Y | 10-04-21 | N Engl J Med | Variant of TYR and autoimmu | Vitiligo | 10p15.1 | 3,00E-09 | | | |
| PFKFB3 | chr10 | 6056985 | 6056986 | **rs706778** | 33106285 | Lopez-Isac E | 20-10-26 | Ann Rheum | Combined genetic analysis of | Arthritis (juvenile idiopathic) | 10p15.1 | 1,00E-12 | Y | | |
| PFKFB3 | chr10 | 6056985 | 6056986 | **rs706778** | 30595370 | Kichaev G | 18-12-27 | Am J Hum G | Leveraging Polygenic Function | Autoimmune traits | 10p15.1 | 4,00E-18 | Y | | |
| PFKFB3 | chr10 | 6056985 | 6056986 | **rs706778** | 26301688 | Li YR | 15-08-24 | Nat Med | Meta-analysis of shared gene | Pediatric autoimmune diseases | 10p15.1 | 6,00E-09 | Y | | |
| PFKFB3 | chr10 | 6056985 | 6056986 | **rs706778** | 30423114 | Laufer VA | 18-11-13 | Hum Mol Ge | Genetic Influences on Suscept | Rheumatoid arthritis | 10p15.1 | 2,00E-11 | Y | | |
| PFKFB3 | chr10 | 6056985 | 6056986 | **rs706778** | 30423114 | Laufer VA | 18-11-13 | Hum Mol Ge | Genetic Influences on Suscept | Rheumatoid arthritis | 10p15.1 | 7,00E-12 | Y | | |
| PFKFB3 | chr10 | 6056985 | 6056986 | **rs706778** | 24390342 | Okada Y | 13-12-25 | Nature | Genetics of rheumatoid arthri | Rheumatoid arthritis | 10p15.1 | 5,00E-14 | Y | | |
| PFKFB3 | chr10 | 6056985 | 6056986 | **rs706778** | 24390342 | Okada Y | 13-12-25 | Nature | Genetics of rheumatoid arthri | Rheumatoid arthritis | 10p15.1 | 5,00E-15 | Y | | |
| PFKFB3 | chr10 | 6056985 | 6056986 | **rs706778** | 20453842 | Stahl EA | 10-05-09 | Nat Genet | Genome-wide association stu | Rheumatoid arthritis | 10p15.1 | 1,00E-11 | Y | | |
| PFKFB3 | chr10 | 6057081 | 6057082 | **rs2104286** | 26974007 | Ellinghaus D | 16-03-14 | Nat Genet | Analysis of five chronic inflam | Chronic inflammatory diseases (ankylosing spondylitis, Crohn's disease, psoriasis | 10p15.1 | 4,00E-08 | Y | | |
| PFKFB3 | chr10 | 6057081 | 6057082 | **rs2104286** | 24076602 | Beecham AH | 13-11-01 | Nat Genet | Analysis of immune-related lo | Multiple sclerosis | 10p15.1 | 2,00E-47 | Y | | |
| PFKFB3 | chr10 | 6057081 | 6057082 | **rs2104286** | 19525955 | Bahlo M | 09-06-14 | Nat Genet | Genome-wide association stu | Multiple sclerosis | 10p15.1 | 7,00E-06 | Y | | |
| PFKFB3 | chr10 | 6057081 | 6057082 | **rs2104286** | 19525953 | De Jager PL | 09-06-14 | Nat Genet | Meta-analysis of genome sca | Multiple sclerosis | 10p15.1 | 9,00E-08 | Y | | |
| PFKFB3 | chr10 | 6057081 | 6057082 | **rs2104286** | 24449572 | Orozco G | 14-01-01 | Arthritis Rhe | Novel rheumatoid arthritis sus | Rheumatoid arthritis | 10p15.1 | 1,00E-06 | Y | | |
| PFKFB3 | chr10 | 6058761 | 6058762 | **rs3134883** | 33310728 | Ha E | 20-12-11 | Ann Rheum | Large-scale meta-analysis acr | Rheumatoid arthritis | 10p15.1 | 2,00E-15 | Y | | |
| PFKFB3 | chr10 | 6059165 | 6059166 | **rs3118469** | 32888494 | Vuckovic D | 20-09-01 | Cell | The Polygenic and Monogenic | Lymphocyte counts | 10p15.1 | 5,00E-25 | Y | | |
| PFKFB3 | chr10 | 6059165 | 6059166 | **rs3118469** | 32888493 | Chen MH | 20-09-01 | Cell | Trans-ethnic and Ancestry-Spe | Lymphocyte counts | 10p15.1 | 2,00E-43 | Y | | |
| PFKFB3 | chr10 | 6059165 | 6059166 | **rs3118469** | 32888493 | Chen MH | 20-09-01 | Cell | Trans-ethnic and Ancestry-Spe | Lymphocyte counts | 10p15.1 | 2,00E-45 | Y | | |
| PFKFB3 | chr10 | 6059165 | 6059166 | **rs3118469** | 32888494 | Vuckovic D | 20-09-01 | Cell | The Polygenic and Monogenic | Monocyte count | 10p15.1 | 5,00E-25 | Y | | |
| PFKFB3 | chr10 | 6059165 | 6059166 | **rs3118469** | 32888494 | Vuckovic D | 20-09-01 | Cell | The Polygenic and Monogenic | White blood cell count | 10p15.1 | 5,00E-10 | Y | | |

| | | | | | | | | | | | | | | | |
|---|---|---|---|---|---|---|---|---|---|---|---|---|---|---|---|
| PFKFB3 | chr10 | 6059165 | 6059166 | **rs3118469** | 30595370 | Kichaev G | 18-12-27 | Am J Hum G | Leveraging Polygenic Function | White blood cell count | 10p15.1 | 8,00E-10 | Y | | |
| PFKFB3 | chr10 | 6059749 | 6059750 | **rs3118470** | 25608926 | Betz RC | 15-01-22 | Nat Commur | Genome-wide meta-analysis | Alopecia areata | 10p15.1 | 8,00E-21 | | | |
| PFKFB3 | chr10 | 6059749 | 6059750 | **rs3118470** | 20596022 | Petukhova L | 10-07-01 | Nature | Genome-wide association stu | Alopecia areata | 10p15.1 | 2,00E-12 | | | |
| PFKFB3 | chr10 | 6059749 | 6059750 | **rs3118470** | 30572963 | Marquez A | 18-12-20 | Genome Me | Meta-analysis of Immunochip | Autoimmune traits (pleiotropy) | 10p15.1 | 6,00E-09 | Y | | |
| PFKFB3 | chr10 | 6059749 | 6059750 | **rs3118470** | 21833088 | Sawcer S | 11-08-11 | Nature | Genetic risk and a primary rol | Multiple sclerosis | 10p15.1 | 3,00E-11 | Y | | |
| PFKFB3 | chr10 | 6059749 | 6059750 | **rs3118470** | 30572963 | Marquez A | 18-12-20 | Genome Me | Meta-analysis of Immunochip | Rheumatoid arthritis | 10p15.1 | 1,00E-06 | Y | | |
| PFKFB3 | chr10 | 6059749 | 6059750 | **rs3118470** | 30572963 | Marquez A | 18-12-20 | Genome Me | Meta-analysis of Immunochip | Type 1 diabetes | 10p15.1 | 1,00E-10 | Y | | |
| PFKFB3 | chr10 | 6060048 | 6060049 | **rs12722489** | 27532455 | Sun W | 16-08-17 | PLoS Genet | Common Genetic Polymorphis | Blood protein levels | 10p15.1 | 3,00E-20 | Y | | |
| PFKFB3 | chr10 | 6060048 | 6060049 | **rs12722489** | 21102463 | Franke A | 10-11-21 | Nat Genet | Genome-wide meta-analysis | Crohn's disease | 10p15.1 | 3,00E-09 | Y | | |
| PFKFB3 | chr10 | 6060048 | 6060049 | **rs12722489** | 22190364 | Patsopoulos | 11-12-01 | Ann Neurol | Genome-wide meta-analysis | Multiple sclerosis | 10p15.1 | 4,00E-08 | Y | | |
| PFKFB3 | chr10 | 6060048 | 6060049 | **rs12722489** | 17660530 | Hafler DA | 07-07-29 | N Engl J Med | Risk alleles for multiple scler | Multiple sclerosis | 10p15.1 | 3,00E-08 | Y | | |
| PFKFB3 | chr10 | 6060793 | 6060794 | **rs3118471** | 26974007 | Ellinghaus D | 16-03-14 | Nat Genet | Analysis of five chronic inflam | Chronic inflammatory diseases (ankylosing spondylitis, Crohn's disease, psoriasis | 10p15.1 | 4,00E-13 | Y | | |
| PFKFB3 | chr10 | 6060793 | 6060794 | **rs3118471** | 32888493 | Chen MH | 20-09-01 | Cell | Trans-ethnic and Ancestry-Spe | White blood cell count | 10p15.1 | 2,00E-11 | Y | | |
| PFKFB3 | chr10 | 6061798 | 6061799 | **rs12722486** | 24554482 | Leger PD | 14-02-20 | J Neurovirol | Genome-wide association stu | Response to anti-retroviral therapy (ddI/d4T) in HIV-1 infection (Grade 3 periphe | 10p15.1 | 2,00E-09 | | | |
| PFKFB3 | chr10 | 6064302 | 6064303 | **rs7072793** | 28100725 | Chen J | 17-01-18 | Neurology | Genome-wide association stu | Plasma t-tau levels | 10p15.1 | 8,00E-06 | | | |
| PFKFB3 | chr10 | 6066376 | 6066377 | **rs10795791** | 32888494 | Vuckovic D | 20-09-01 | Cell | The Polygenic and Monogenic | Eosinophil counts | 10p15.1 | 5,00E-13 | | | |
| PFKFB3 | chr10 | 6066376 | 6066377 | **rs10795791** | 32888494 | Vuckovic D | 20-09-01 | Cell | The Polygenic and Monogenic | Eosinophil percentage of white cells | 10p15.1 | 5,00E-09 | | | |
| PFKFB3 | chr10 | 6066376 | 6066377 | **rs10795791** | 23143596 | Eyre S | 12-12-01 | Nat Genet | High-density genetic mapping | Rheumatoid arthritis | 10p15.1 | 1,00E-09 | Y | | |
| PFKFB3 | chr10 | 6066376 | 6066377 | **rs10795791** | 24532676 | Kim K | 14-02-14 | Ann Rheum | High-density genotyping of im | Rheumatoid arthritis (ACPA-positive) | 10p15.1 | 1,00E-10 | Y | | |
| PFKFB3 | chr10 | 6066376 | 6066377 | **rs10795791** | 23143596 | Eyre S | 12-12-01 | Nat Genet | High-density genetic mapping | Rheumatoid arthritis (ACPA-positive) | 10p15.1 | 1,00E-09 | Y | | |
| PFKFB3 | chr10 | 6066376 | 6066377 | **rs10795791** | 25751624 | Onengut-Gu | 15-04-01 | Nat Genet | Fine mapping of type 1 diabet | Type 1 diabetes | 10p15.1 | 6,00E-11 | Y | | |
| PFKFB3 | chr10 | 6066475 | 6066476 | **rs4147359** | 27992413 | Ji SG | 16-12-19 | Nat Genet | Genome-wide association stu | Primary sclerosing cholangitis | 10p15.1 | 8,00E-17 | | | |
| PFKFB3 | chr10 | 6066475 | 6066476 | **rs4147359** | 23603763 | Liu JZ | 13-06-01 | Nat Genet | Dense genotyping of immune- | Primary sclerosing cholangitis | 10p15.1 | 8,00E-17 | | | |
| PFKFB3 | chr10 | 6068865 | 6068866 | **rs7090512** | 21833088 | Sawcer S | 11-08-11 | Nature | Genetic risk and a primary rol | Multiple sclerosis | 10p15.1 | 5,00E-20 | Y | | |
| PFKFB3 | chr10 | 6068911 | 6068912 | **rs7090530** | 32581359 | Saevarsdotti | 20-06-24 | Nature | FLT3 stop mutation increases | Autoimmune thyroid disease | 10p15.1 | 2,00E-26 | Y | | |
| PFKFB3 | chr10 | 6068911 | 6068912 | **rs7090530** | 30595370 | Kichaev G | 18-12-27 | Am J Hum G | Leveraging Polygenic Function | Hypothyroidism | 10p15.1 | 5,00E-19 | Y | | |
| PFKFB3 | chr10 | 6068911 | 6068912 | **rs7090530** | 31015401 | Wu Y | 19-04-23 | Nat Commur | Genome-wide association stu | Medication use (thyroid preparations) | 10p15.1 | 7,00E-24 | | | |
| PFKFB3 | chr10 | 6069111 | 6069112 | **rs7893324** | 31669095 | Zhu Z | 19-10-24 | J Allergy Clin | Shared Genetic and Experime | Atopic asthma | 10p15.1 | 3,00E-09 | | | |
| PFKFB3 | chr10 | 6072046 | 6072047 | **rs10905716** | 26293465 | Durda P | 15-08-20 | Arterioscler | Plasma Levels of Soluble Inter | Soluble interleukin-2 receptor subunit alpha | 10p15.1 | 3,00E-09 | Y | | |
| PFKFB3 | chr10 | 6072892 | 6072893 | **rs10905718** | 32888494 | Vuckovic D | 20-09-01 | Cell | The Polygenic and Monogenic | Lymphocyte percentage of white cells | 10p15.1 | 5,00E-10 | Y | | |
| PFKFB3 | chr10 | 6072892 | 6072893 | **rs10905718** | 32888494 | Vuckovic D | 20-09-01 | Cell | The Polygenic and Monogenic | Neutrophil percentage of white cells | 10p15.1 | 2,00E-10 | Y | | |
| PFKFB3 | chr10 | 6072892 | 6072893 | **rs10905718** | 28714469 | Langefeld CD | 17-07-17 | Nat Commur | Transancestral mapping and g | Systemic lupus erythematosus | 10p15.1 | 4,00E-06 | Y | | |
| PFKFB3 | chr10 | 6073675 | 6073676 | **rs62626322** | 31619474 | Zhu Z | 19-10-17 | Eur Respir J | Shared Genetics of Asthma ar | Asthma | 10p15.1 | 4,00E-08 | Y | | |
| PFKFB3 | chr10 | 6073675 | 6073676 | **rs62626322** | 31361310 | Johansson A | 19-07-30 | Hum Mol Ge | Genome-wide association ana | Asthma | 10p15.1 | 3,00E-09 | Y | | |
| PFKFB3 | chr10 | 6074290 | 6074291 | **rs72776098** | 30572963 | Marquez A | 18-12-20 | Genome Me | Meta-analysis of Immunochip | Autoimmune traits (pleiotropy) | 10p15.1 | 7,00E-10 | Y | | |
| PFKFB3 | chr10 | 6074290 | 6074291 | **rs72776098** | 30572963 | Marquez A | 18-12-20 | Genome Me | Meta-analysis of Immunochip | Type 1 diabetes | 10p15.1 | 6,00E-09 | Y | | |
| PFKFB3 | chr10 | 6075358 | 6075359 | **rs11256593** | 31604244 | International | 19-09-01 | Science | Multiple sclerosis genomic ma | Multiple sclerosis | 10p15.1 | 3,00E-65 | Y | | |
| PFKFB3 | chr10 | 6079321 | 6079322 | **rs62626325** | 24076602 | Beecham AH | 13-11-01 | Nat Genet | Analysis of immune-related lo | Multiple sclerosis | 10p15.1 | 9,00E-06 | Y | | |
| PFKFB3 | chr10 | 6080045 | 6080046 | **rs11594656** | 33310728 | Ha E | 20-12-11 | Ann Rheum | Large-scale meta-analysis acr | Rheumatoid arthritis | 10p15.1 | 4,00E-09 | Y | | |
| PFKFB3 | chr10 | 6080705 | 6080706 | **rs2182410** | 27386562 | Andlauer TF | 16-06-17 | Sci Adv | Novel multiple sclerosis susce | Multiple sclerosis | 10p15.1 | 4,00E-09 | Y | | |
| PFKFB3 | chr10 | 6081531 | 6081532 | **rs12251307** | 19430480 | Barrett JC | 09-05-10 | Nat Genet | Genome-wide association stu | Type 1 diabetes | 10p15.1 | 1,00E-13 | Y | | |
| PFKFB3 | chr10 | 6081531 | 6081532 | **rs12251307** | 18978792 | Cooper JD | 08-11-02 | Nat Genet | Meta-analysis of genome-wid | Type 1 diabetes | 10p15.1 | 2,00E-06 | Y | | |
| PFKFB3 | chr10 | 6082657 | 6082658 | **rs41295115** | 24388013 | Ferreira MA | 13-12-30 | J Allergy Clin | Genome-wide association ana | Asthma and hay fever | 10p15.1 | 5,00E-07 | Y | | |
| PFKFB3 | chr10 | 6087346 | 6087347 | **rs75641219** | 32493714 | Klimentidis Y | 20-06-03 | Diabetes | Phenotypic and Genetic Chara | Low density lipoprotein cholesterol levels | 10p15.1 | 3,00E-08 | | | |
| PFKFB3 | chr10 | 6087679 | 6087680 | **rs41295121** | 25751624 | Onengut-Gu | 15-04-01 | Nat Genet | Fine mapping of type 1 diabet | Type 1 diabetes | 10p15.1 | 5,00E-08 | Y | | |
| PFKFB3 | chr10 | 6107073 | 6107074 | **rs12778662** | 30134952 | Traglia M | 18-08-22 | Genome Me | Cross-genetic determination c | Midgestational cytokine/chemokine levels (maternal genetic effect) | 10p15.1 | 4,00E-09 | | | |
| PFKFB3 | chr10 | 6122346 | 6122347 | **rs1073646** | 30886212 | Eszlari N | 19-03-18 | Transl Psychi | Genome-wide association ana | Brooding (response to stress) | 10p15.1 | 3,00E-06 | | | |
| PFKFB3 | chr10 | 6125160 | 6125161 | **rs10905856** | 31194736 | Tiensuu H | 19-06-13 | PLoS Genet | Risk of spontaneous preterm l | Spontaneous preterm birth (preterm birth) | 10p15.1 | 5,00E-06 | | | |
| PFKFB3 | chr10 | 6127176 | 6127177 | **rs11256909** | 29678681 | Matana A | 18-04-18 | Genomics | Genome-wide meta-analysis | Plasma anti-thyroid peroxidase levels | 10p15.1 | 1,00E-06 | | | |
| PFKFB3 | chr10 | 6129671 | 6129672 | **rs5014463** | 31926482 | Shen H | 20-01-11 | Neuropsycho | Polygenic prediction and GWA | Post-traumatic stress disorder | 10p15.1 | 5,00E-06 | | | |
| PFKFB3 | chr10 | 6131729 | 6131730 | **rs7924005** | 26293465 | Durda P | 15-08-20 | Arterioscler | Plasma Levels of Soluble Inter | Soluble interleukin-2 receptor subunit alpha | 10p15.1 | 4,00E-10 | Y | | |
| PFKFB3 | chr10 | 6131813 | 6131814 | **rs10905875** | 27723758 | Bronson PG | 16-10-10 | Nat Genet | Common variants at PVT1, AT | Selective IgA deficiency | 10p15.1 | 1,00E-06 | Y | | |
| PFKFB3 | chr10 | 6136650 | 6136651 | **rs1983890** | 30254083 | Cousminer D | 18-09-25 | Diabetes Car | First Genome-Wide Associati | Latent autoimmune diabetes | 10p15.1 | 3,00E-08 | Y | | |
| PFKFB3 | chr10 | 6166047 | 6166048 | **rs186003188** | 32141793 | Garman L | 20-03-06 | Ocul Immuno | Genome-Wide Association Stu | Ocular sarcoidosis | 10p15.1 | 1,00E-07 | | | |
| PFKFB3 | chr10 | 6171300 | 6171301 | **rs184694238** | 28915241 | Ahsan M | 17-09-15 | PLoS Genet | The relative contribution of DI | Blood protein levels | 10p15.1 | 4,00E-11 | | | |
| PFKFB3 | chr10 | 6186262 | 6186263 | **rs666595** | 31888951 | Schlauch KA | 19-12-30 | G3 (Bethesd | A Comprehensive Genome-W | Obesity (extreme) | 10p15.1 | 6,00E-06 | | | |
| PFKFB3 | chr10 | 6187610 | 6187611 | **rs631902** | 25608926 | Betz RC | 15-01-22 | Nat Commur | Genome-wide meta-analysis | Alopecia areata | 10p15.1 | 5,00E-06 | | | |
| PFKFB3 | chr10 | 6189821 | 6189822 | **rs186795497** | 27989323 | Ahola-Olli A | 16-12-13 | Am J Hum G | Genome-wide Association Stu | Interleukin-2 receptor antagonist levels | 10p15.1 | 3,00E-07 | Y | | |
| PFKFB3 | chr10 | 6195192 | 6195193 | **rs11257188** | 25117820 | Schneider BP | 14-08-12 | Br J Cancer | Genetic variant predicts bevac | Response to chemotherapy in breast cancer hypertensive cases (cumulative dose | 10p15.1 | 9,00E-06 | | | |
| PFKFB3 | chr10 | 6236347 | 6236348 | **rs113152482** | 28644415 | Guenard F | 17-06-23 | Nutrients | Genome-Wide Association Stu | Western dietary pattern | 10p15.1 | 8,00E-07 | | | |
| PFKFB3 | chr10 | 6273830 | 6273831 | **rs7077335** | 23049088 | Meng W | 12-10-09 | Invest Ophth | A genome-wide association st | Myopia (pathological) | 10p15.1 | 2,00E-12 | | | |
| PFKFB3 | chr10 | 6299719 | 6299720 | **rs185254058** | 28915241 | Ahsan M | 17-09-15 | PLoS Genet | The relative contribution of DI | Blood protein levels | 10p15.1 | 1,00E-10 | | | |
| PFKFB3 | chr10 | 6348229 | 6348230 | **rs2387397** | 22057235 | Trynka G | 11-11-06 | Nat Genet | Dense genotyping identifies a | Celiac disease | 10p15.1 | 2,00E-08 | Y | | Y: PFKFB3_GH10F006346 |
| PFKFB3 | chr10 | 6348487 | 6348488 | **rs947474** | 30572963 | Marquez A | 18-12-20 | Genome Me | Meta-analysis of Immunochip | Autoimmune traits (pleiotropy) | 10p15.1 | 1,00E-08 | Y | | Y: PFKFB3_GH10F006346 |
| PFKFB3 | chr10 | 6348487 | 6348488 | **rs947474** | 30572963 | Marquez A | 18-12-20 | Genome Me | Meta-analysis of Immunochip | Celiac disease | 10p15.1 | 2,00E-07 | Y | | Y: PFKFB3_GH10F006346 |
| PFKFB3 | chr10 | 6348487 | 6348488 | **rs947474** | 26546613 | Gutierrez-Ac | 16-01-01 | Hum Mol Ge | Functional implications of dis | Celiac disease and Rheumatoid arthritis | 10p15.1 | 8,00E-11 | Y | | Y: PFKFB3_GH10F006346 |
| PFKFB3 | chr10 | 6348487 | 6348488 | **rs947474** | 30572963 | Marquez A | 18-12-20 | Genome Me | Meta-analysis of Immunochip | Rheumatoid arthritis | 10p15.1 | 2,00E-07 | Y | | Y: PFKFB3_GH10F006346 |
| PFKFB3 | chr10 | 6348487 | 6348488 | **rs947474** | 30423114 | Laufer VA | 18-11-13 | Hum Mol Ge | Genetic Influences on Suscept | Rheumatoid arthritis | 10p15.1 | 3,00E-08 | Y | | Y: PFKFB3_GH10F006346 |
| PFKFB3 | chr10 | 6348487 | 6348488 | **rs947474** | 30423114 | Laufer VA | 18-11-13 | Hum Mol Ge | Genetic Influences on Suscept | Rheumatoid arthritis | 10p15.1 | 3,00E-09 | Y | | Y: PFKFB3_GH10F006346 |
| PFKFB3 | chr10 | 6348487 | 6348488 | **rs947474** | 24390342 | Okada Y | 13-12-25 | Nature | Genetics of rheumatoid arthri | Rheumatoid arthritis | 10p15.1 | 4,00E-10 | Y | | Y: PFKFB3_GH10F006346 |
| PFKFB3 | chr10 | 6348487 | 6348488 | **rs947474** | 24390342 | Okada Y | 13-12-25 | Nature | Genetics of rheumatoid arthri | Rheumatoid arthritis | 10p15.1 | 3,00E-10 | Y | | Y: PFKFB3_GH10F006346 |
| PFKFB3 | chr10 | 6348487 | 6348488 | **rs947474** | 24532676 | Kim K | 14-02-14 | Ann Rheum | High-density genotyping of im | Rheumatoid arthritis (ACPA-positive) | 10p15.1 | 1,00E-08 | Y | | Y: PFKFB3_GH10F006346 |
| PFKFB3 | chr10 | 6348487 | 6348488 | **rs947474** | 18978792 | Cooper JD | 08-11-02 | Nat Genet | Meta-analysis of genome-wid | Type 1 diabetes | 10p15.1 | 4,00E-09 | Y | | Y: PFKFB3_GH10F006346 |
| PFKFB3 | chr10 | 6349098 | 6349099 | **rs2181622** | 30595370 | Kichaev G | 18-12-27 | Am J Hum G | Leveraging Polygenic Function | Autoimmune traits | 10p15.1 | 6,00E-11 | Y | | Y: PFKFB3_GH10F006346 |
| PFKFB3 | chr10 | 6351046 | 6351047 | **rs744253** | 31015401 | Wu Y | 19-04-23 | Nat Commur | Genome-wide association stu | Medication use (thyroid preparations) | 10p15.1 | 4,00E-15 | | | |
| PFKFB3 | chr10 | 6351297 | 6351298 | **rs4750316** | 20453842 | Stahl EA | 10-05-09 | Nat Genet | Genome-wide association stu | Rheumatoid arthritis | 10p15.1 | 2,00E-06 | Y | | |
| PFKFB3 | chr10 | 6351297 | 6351298 | **rs4750316** | 18794853 | Raychaudhur | 08-09-14 | Nat Genet | Common variants at CD40 and | Rheumatoid arthritis | 10p15.1 | 4,00E-06 | Y | | |
| PFKFB3 | chr10 | 6359662 | 6359663 | **rs4558075** | 24999842 | Garner C | 14-07-07 | PLoS One | Genome-wide association stu | Celiac disease | 10p15.1 | 2,00E-07 | Y | | |
| PFKFB3 | chr10 | 6363571 | 6363572 | **rs11258303** | 30595370 | Kichaev G | 18-12-27 | Am J Hum G | Leveraging Polygenic Function | Hypothyroidism | 10p15.1 | 7,00E-13 | Y | | |
| PFKFB3 | chr10 | 6365774 | 6365775 | **rs11258317** | 20889312 | Wang KS | 10-10-01 | Schizophr Re | A genome-wide meta-analysi | Bipolar disorder and schizophrenia | 10p15.1 | 6,00E-06 | | | |
| PFKFB3 | chr10 | 6406931 | 6406932 | **rs586511** | 31015401 | Wu Y | 19-04-23 | Nat Commur | Genome-wide association stu | Medication use (thyroid preparations) | 10p15.1 | 4,00E-09 | | | |
| PFKFB3 | chr10 | 6419032 | 6419033 | **rs11258664** | 30642921 | de Vries PS | 19-01-14 | Blood | A genome-wide association st | Factor VII activity | 10p15.1 | 2,00E-06 | | | |
| PFKFB3 | chr10 | 6430928 | 6430929 | **rs11258747** | 19430480 | Barrett JC | 09-05-10 | Nat Genet | Genome-wide association stu | Type 1 diabetes | 10p15.1 | 1,00E-07 | Y | | |
| PFKFB3 | chr10 | 6013640 | 6013641 | **rs9663421** | 31619474 | Zhu Z | 19-10-17 | Eur Respir J | Shared Genetics of Asthma ar | Asthma | 10p15.1 | 3,00E-10 | | | |

Supplementary Table 8S. Clinical characteristics RA patients

| | 22 patients FACS | 24 patients RNAseq |
|---|---|---|
| Age, years Average[min-max] | 62.1[37-73] | 55 [46.5-66] |
| gender, F | 16 (72.7%) | 24 (100%) |
| DD, years Average[min-max] | 19.5[9-47] | 5.5 [1.75-11.25] |
| seropositive | 17 | 16 (+3?) |
| Treatment | | |
| MTX | 20 | 16 |
| aTNF | 20 | 5 |
| Other biologics | 2 | 4 |

Supplementary table 9S. Primers used in the cell experiments

| Target | Forward | Reverse |
|---|---|---|
| SLC2A1 | ACTGGAGTCATCAATGCCCC | AGAAGGAGCCAATCATGCCC |
| PFKFB3 | CCTACAACTTCTTCCGCCCC | CCGCAATTTGTCCCCCTTCT |
| BIRC5 | GACCACCGCATCTCTACATTC | TGCTTTTTATGTTCCTCTATGGG |
